# Evaluating Transferability and Robustness of Process-Guided Neural Networks in Forest Carbon Flux Modelling

**DOI:** 10.64898/2026.02.24.707715

**Authors:** Hannah Habenicht, Hanne Raum, Joschka Boedecker, Carsten F. Dormann

**Author notes:** Corresponding author *Email address:* (Hannah Habenicht). These authors contributed equally to this work (Hannah Habenicht, Hanne Raum).

## Abstract

Making robust and generalisable predictions within ecological systems such as forests remains a challenge due to limited data availability and the slow pace of environmental change. To address this, we integrated a semi-empirical environmental process model (PRELES) to support deep learning approaches, specifically artificial neural networks (ANNs). We replicated and extended previous work on process-guided neural networks (PGNN) by introducing new model types and conducting a comprehensive hyperparameter optimisation. These were then evaluated within systematic nested cross-validation analyses covering data-thinning and extrapolative scenarios. Data-driven ANNs and PGNNs alike outperformed the stand-alone process model on average, while certain PGNNs provided additional advantages over plain ANNs when training data were scarce and when models were transferred to climatically novel sites. We further estimated the generalisation error for data-driven models as a function of training data size, giving practical guidance on model suitability under different data availability and distribution. A variable-importance analysis using accumulated local effects (ALE) revealed that both PGNNs and ANNs learn simple, physically plausible relationships. Further analyses show where the process model (PRELES) provides useful guidance to the PGNNs and where its limitations may constrain the latter’s performance.

**Graphical Abstract:** 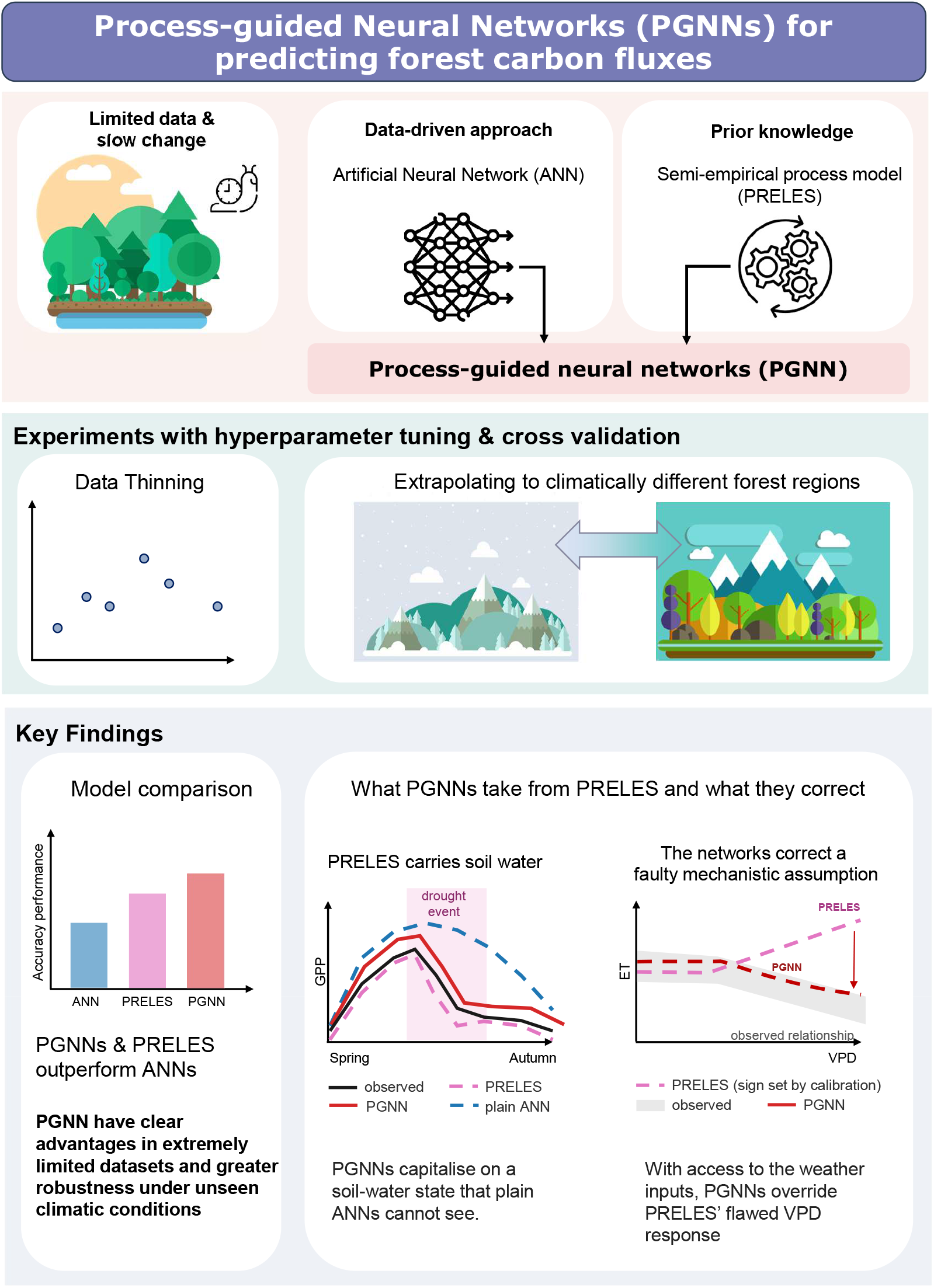

**Highlights:**

- Process-guided, and plain neural networks outperform a calibrated process-based model (PRELES) in predicting forest ecosystem carbon fluxes.
- Process-guided neural networks provide advantages over plain ANN in sparse-data settings and show greater robustness under transferable scenarios with unseen changing climatic conditions.
- Variable-importance analyses using accumulated local effects show that both process-guided and plain neural networks learn simple yet physically plausible relationships between meteorological drivers and target responses, while PGNNs (bias) benefited from the process model (PRELES) during drought conditions but also inherited some of its physically implausible constraints on ET, which they could partially correct.

## 1. Introduction

Environmental systems are inherently challenging to predict, as they emerge from many interacting drivers whose relationships can vary across both space and time. Forest ecosystems, for instance, exhibit highly diverse patterns across different climatic regions. Mechanisms that appear stable in the pine-dominated boreal forest in Finland may not generalise to oceanic forests in coastal France or Mediterranean beech forests in Italy. This ecological variability underscores a broader challenge: how can we construct modelling approaches that remain reliable across complex and heterogeneous environmental settings?

To tackle this complexity, researchers have increasingly turned to data-driven approaches such as machine learning (Hsieh, 2009), as artificial neural networks (plain ANNs) can capture complex non-linear relationships when sufficient high-quality data are available. However, their performance tends to degrade in data-scarce settings, which can produce unreliable or scientifically inconsistent results (Karimi et al., 2020; Zhou et al., 2022; Pichler and Hartig, 2023; Hackenberg et al., 2025). This is often the case in ecological modelling, where long time horizons and costly data collection limit available training data. Additionally, data-driven models tend to struggle when applied to conditions that differ from their training setting, which limits their practical use for generalising to out-of-distribution (OOD) contexts (Willard et al., 2022; Karpatne et al., 2024).

Efforts to overcome these limitations have introduced hybrid or informed machine learning methods (Reichstein et al., 2019; Von Rueden et al., 2021), where prior scientific knowledge is incorporated into the training process, most commonly via the loss function. These methods emerged primarily in physics and physics-adjacent domains and have shown promise in environmental applications where governing equations are well-understood. For example: predicting lake temperature using energy conservation laws (Jia et al., 2021), modelling climate and weather patterns using neural ordinary differential equations that integrate conservation-law-based transport dynamics (Verma et al., 2024), or forecasting air quality using graph-based differential equation networks that capture pollutant dispersion and air flow dynamics (Hettige et al., 2024). Although these approaches differ in their technical implementations, they share a common foundation: encoding known system dynamics to improve model robustness and maintain physical consistency.

Forest carbon flux modelling, however, does not offer this convenience. Many drivers and parameters are collinear, interact non-linearly, and are only partially observable (Dormann et al., 2013), so governing equations that ODE-based hybrids rely on are rarely available in closed form. Process models of forest systems instead *approximate* key mechanisms as simplified algebraic forms. The process model (PREdict Light-use efficiency, Evapotranspiration and Soil water, PRELES) is one such model: a calibrated semi-empirical process model used to estimate gross primary production (GPP), evapotranspiration (ET), and soil water dynamics (Peltoniemi et al., 2015; Mäkelä et al., 2008). This structural difference may explain why hybrid approaches in forest carbon flux modelling remain rare. Only a few studies (Koppa et al., 2022; Fang and Gentine, 2024) embed process knowledge within machine learning frameworks for carbon and water fluxes. Those that exist either train and evaluate on the same global pool of sites without testing transfer to unseen locations or evaluate performance within the same site across time (Koppa et al., 2022; Fang and Gentine, 2024), leaving open how hybrid strategies behave under data scarcity or when transferred to climatically novel forest sites.

Here, the process model (PRELES) serves as a process-based constraint whose predictions are incorporated into plain ANNs using five different strategies. This produces a set of process-guided neural networks (PGNNs), collectively referred to as hybrid models. Previous work by Wesselkamp et al. (2024) showed that hybrid models can outperform both the process model and plain ANNs under sparse data scenarios. However, it remains unclear how each modelling strategy – plain ANN, hybrid or process – systematically responds to varying levels of data availability. Additionally, process models are often assumed to generalise better in OOD settings due to their structural rigidity (Willard et al., 2022; Karpatne et al., 2024). However, the OOD analysis by Wesselkamp et al. (2024) tested this assumption only in a limited setting, evaluating trained models at Hyytiälä (Finland), the site where the process model (PRELES) was originally developed. Consequently, we still lack a clear understanding of how these models behave when confronted with a broader range of environmental conditions found across European forests.

In this study, we address the issue of data availability and predictions in OOD settings through a series of nested cross-validations (NCV). Specifically, we vary the amount of training data and test model performance at the training sites (in-distribution, ID) and across unseen test sites (out-of-distribution, OOD). These experiments allow us to estimate and compare the generalisation errors for each modelling strategy, and assess the amount of training data required to reliably predict under both data-thinning and OOD conditions. We further assess whether incorporating the mechanistic knowledge encoded in the process model (PRELES) improves the predictions of hybrid strategies relative to purely data-driven neural networks and the process model (PRELES) itself. Finally, to better understand the differences in prediction patterns between the process-based model (PRELES) and the PGNNs, and to identify their respective strengths and limitations, we analysed cases of poor performance using tools such as variable associations. Two research questions guide this analysis: (1) How do modelling strategies perform when training data are systematically reduced, evaluated separately under in-distribution and out-of-distribution scenarios? (2) Can changes in environmental conditions explain differences in predictive performance across modelling strategies? Ultimately, our analysis broadens the perspective of model transferability and extrapolative ability by linking performance differences to varying sites and ecological contexts. Our results show that (1) data scarcity is not inherently limiting, (2) hybrids relying on the process model can inherit useful information not supplied by the data. (3) Variable importance analyses can point towards inaccurate mechanistic assumptions in the process model, and (4) overall, hybrid models achieve more reliable performance across diverse climates, site characteristics, and data regimes found in forest ecosystems.

## 2. Background and Methods

The following sections introduce the process model (PRELES), the datasets, and strategies for integrating process-based predictions with neural networks. We outline the evaluation framework, covering training strategies, approximations of generalisation errors across models, and methods used to analyse model behaviour using accumulated local effects (ALE).

### 2.1. Process Model (PM): PRELES

The mechanistic component, which introduces prior scientific knowledge into our hybrid frameworks, is provided by the environmental process model (PRELES) (Peltoniemi et al., 2015; Mäkelä et al., 2008). PRELES is a semi-empirical (i.e. the parameters of the mechanistic model components are empirically informed) process model that simulates daily gross primary production (GPP), evapotranspiration (ET) and soil water (SW) using daily meteorological drivers: air temperature (Tair), precipitation (Precip), vapour pressure deficit (VPD), photo-active radiation (PAR) and the fraction of absorbed PAR (fAPAR) by the canopy (as detailed by Peltoniemi et al., 2015; Mäkelä et al., 2008). As a relatively simple model with a limited number of parameters, the process model (PRELES) is easier to apply than more complex mechanistic models. It was originally developed in Finland and validated for boreal forests, including a default parameter set tuned to the latter’s region (Minunno et al., 2016). For our experiments, we inversely calibrated key site-specific parameters, known for their sensitivity to water and carbon cycling processes, for each site and dataset subset (Van Oijen et al., 2005; Sun and Sun, 2015) (details are provided in the Appendix A.2.2). The process model (PRELES) predictions (GPP and ET) from these inversely calibrated mechanistic models were then used as a baseline in various hybrid integration strategies with neural networks, as described below.

### 2.2. Datasets

Data were retrieved from the PROFOUND database (Reyer et al., 2020), which provides aggregated meteorological and remote sensing records at nine European forest sites compiled from other sources (FLUXNET, MODIS etc.). Specific variable requirements of the process model (PRELES) (Minunno et al., 2016) resulted in a final selection of four forest sites: Hyytiälä (Finland), Sorø (Denmark), Collelongo (Italy) and Le Bray (France). They represent the major climatic gradients and forest types throughout Europe, from boreal coniferous forests to Mediterranean broadleaf forests, enabling robust testing of cross-site generalisation (summarised in Table 1).

**Table 1:** Descriptives and site characteristics for the four PROFOUND forest sites. All variables are reported as mean standard deviation from harmonised daily-resolution data across the full dataset. Units follow PROFOUND/FLUXNET conventions.

|  | **Collelongo** | **Hyytiälä** | **Le Bray** | **Sorø** |
|---|---|---|---|---|
| **Site metadata** |  |  |  |  |
| Site ID | 5 | 12 | 14 | 21 |
| Years available (used) | 2001–2013 (13) | 2004–2012 (8) | 2001–2008 (8) | 2002–2012 (8) |
| Tree type | Deciduous Broadleaf | Evergreen Needleleaf | Evergreen Needleleaf | Deciduous Broadleaf |
| Climate | Mediterranean montane | Boreal | Moderate oceanic | Warm temperate, humid |
| **Environmental drivers (mean ± std)** |  |  |  |  |
| PAR (*µ*mol m<sup>−2</sup> s<sup>−1</sup>) | 27.79 ± 16.40 | 15.34 ± 14.56 | 23.48 ± 15.04 | 18.60 ± 15.28 |
| Tair (°C) | 7.29 ± 6.98 | 4.37 ± 9.53 | 13.18 ± 6.30 | 8.45 ± 6.71 |
| VPD (kPa) | 0.32 ± 0.31 | 0.29 ± 0.33 | 0.47 ± 0.41 | 0.24 ± 0.22 |
| fAPAR (-) | 0.55 ± 0.30 | 0.46 ± 0.32 | 0.66 ± 0.12 | 0.54 ± 0.24 |
| Precip (mm d<sup>−1</sup>) | 3.16 ± 7.30 | 1.86 ± 4.14 | 2.58 ± 5.96 | 2.20 ± 5.61 |
| **Fluxes (mean ± std per site)** |  |  |  |  |
| GPP (gC m<sup>−2</sup> d<sup>−1</sup>) | 4.39 ± 4.41 | 2.87 ± 3.52 | 4.15 ± 3.07 | 5.30 ± 5.65 |
| ET (mm d<sup>−1</sup>) | 1.44 ± 1.52 | 1.09 ± 1.18 | 1.96 ± 1.33 | 1.28 ± 1.37 |

Climate variables (air temperature, precipitation, and global radiation) were sourced from the CLIMATE_LOCAL table at daily resolution. Vapour pressure deficit, latent heat flux and gross primary production (GPP, daytime-partitioned product derived from MDS gap-filled net ecosystem exchange) were obtained from 30-minute records and aggregated to daily means. Latent heat flux was converted to evapotranspiration (ET, mm day^−1^) following (Stull, 1988). Global radiation was converted to photosynthetically active radiation (PAR) following Taiz (2015). GPP was converted to g C m^−2^ d^−1^.

Across the four sites, the provider’s quality flags indicate that approximately 58% of half-hourly GPP values, 37% of latent heat flux, and 16% of VPD are gap-filled (of which 19%, 18%, and 10%, respectively, carry medium- or poor-quality flags). We used GPP and VPD as provided. Before converting heat flux to ET, we corrected implausible negative values of heat flux by the mean of the direct non-zero neighbours, single valid neighbour where only one was available or zero otherwise. We applied no further temporal interpolation to any of these three variables; incomplete years were discarded rather than imputed (see below). Since a substantial fraction of our target variables are gap-filled rather than directly measured, model skill on the gap-filled periods may reflect the gap-filling procedure itself, instead of a genuine prediction of the ecosystem observations. We address this by evaluating model performance on the subset of days with directly measured (non-gap-filled) GPP, LE, and VPD values; see Appendix C.2 for details and results. This analysis shows no evidence that model performance is substantially affected by gap-filling.

We obtained fAPAR from PROFOUND’s MODIS database, which bundles the FPAR/LAI, NDVI/EVI, and other MODIS products available in PROFOUND into a single table joined by site and date. This joined view resulted in a 16-day resolution series, despite the native FPAR/LAI product (MOD15A2) being generated as an 8-day series. Visual inspection revealed many isolated drops to implausibly low values during otherwise active periods (e.g. fAPAR 0.15 during mid-green-up). We opted against the MODIS quality flag, which removed too many winter data points (for instance, 100% at Hyytiälä), and instead applied two de-spiking criteria: a neighbour-based notch test following the logic of the Best Index Slope Extraction (BISE) algorithm (Viovy et al., 1992), and a robust local outlier test following the Hampel identifier (Pearson et al., 2016). A value was flagged as implausible if it fell more than 0.20 below both temporal neighbours, or more than three median absolute deviations (MAD) below a local rolling median with a window length of 7. Together, these criteria identified local dips that were implausible relative to either their immediate neighbours or the broader seasonal pattern. Flagged values were blanked and refilled from the mean of their neighbours; genuine winter lows, which agree with their neighbours, were retained. The threshold and window length were chosen empirically, and their practical impact is assessed in a sensitivity analysis (see Appendix C.1).

All variables were merged and trimmed to complete calendar years in which all essential driver and target variables were fully observed. For reduced-data experiments, we thinned the datasets by retaining one time point per fixed interval (weekly, monthly, or every 50th day) from the full dataset (details in Table 2).

**Table 2:** Overview of training data regimes and thinning strategies for temporal (in-distribution) and spatial experiments (out of distribution). ID experiments were carried out at Hyytiälä and Collelongo. OOD experiments used one to four full-resolution years per training site (a total of three sites) taken from Hyytiälä, Collelongo, Le Bray and Sorø. Thinning intervals (7, 30, 50 days) were applied to the training data to form reduced-data regimes.

| **Training regimes** | **ID: Hyytiälä** | **ID: Collelongo** | **OOD: 3 training sites** |
|---|---|---|---|
| **Sparse regimes** (samples/from) in [days/days] |  |  |  |
| Every 7th day (weekly) | 52/365 | 52/365 | 157/1095 |
| Every 30th day (monthly) | 12/365 | 12/365 | 37/1095 |
| Every 50th day | 7/365 | 7/365 | 22/1095 |
| **Full regimes** with daily resolution in [years] |  |  |  |
| Number of years | 1,2,3 | 1,2,3,5 | 3, 6, 9, 12 (1-4 per site) |
| **Total number of training regimes** | 6 | 7 | 7 |

### 2.3. Hybrid Modelling Strategies

Building on the work of Wesselkamp et al. (2024), we implemented five hybrid strategies to integrate the process model (PRELES) predictions into plain ANNs (*naive* models) as process-guided neural networks (PGNNs). All neural networks are fully connected multi-layer perceptrons (MLPs) trained to predict ET and GPP, using observed values as ground truth (Figure 1). The mathematical formulations of all modelling strategies and associated loss functions can be found in Appendix A.1.

**Figure 1:**
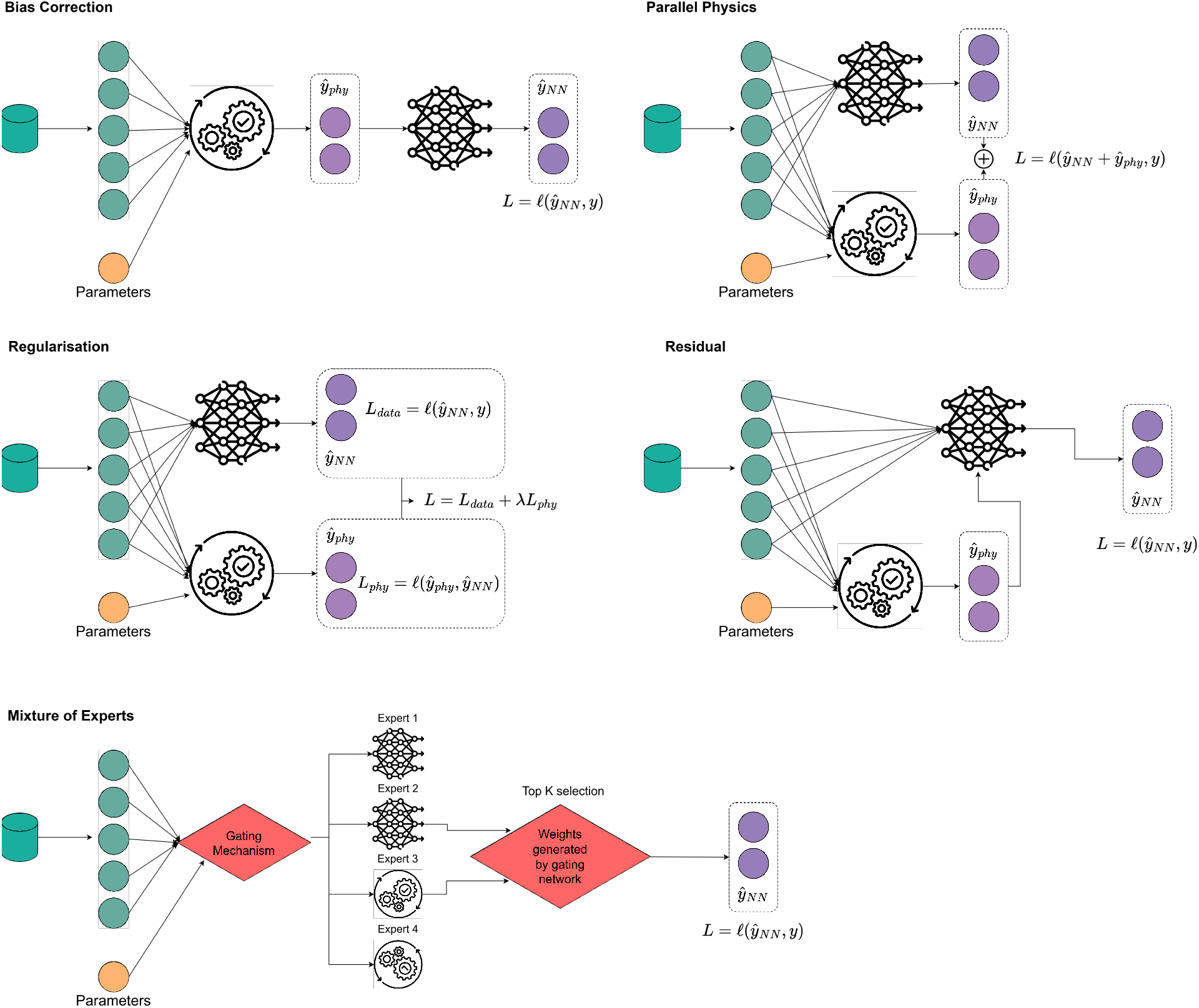
Schematic visualisation, adapted from and expanded from Wesselkamp et al. (2024), showing the hybrid modelling structures of process prediction integration into a neural network, as well as the associated loss function used during training.

The bias correction strategy (*bias*) receives the process model predictions as inputs to an MLP, which learns to correct systematic errors by minimising the loss against observed values. The parallel physics strategy (*parallel*) combines the neural network’s output with the process model predictions before computing the loss against the observed ET and GPP values. The regularized strategy (*regularized*) includes the process model outputs in the loss function by constraining the original loss (difference between measured values and neural network outputs) with an additional regularisation loss term, which defines the correction strength of the process model output relative to the MLP’s output. In addition, we introduced two new strategies:

- **Residual Network:** The residual approach provides an alternative to the parallel, and is an extension of the bias. Instead of adding process model predictions to the outputs, the network receives them as additional inputs alongside the environmental drivers, allowing it to selectively integrate process knowledge and potentially improve learning efficiency.
- **Noisy Mixture of Experts** The Noisy Mixture of Experts (Shazeer et al., 2017) (*mixture*) combines multiple candidate models (experts) through a learnable gating network that assigns weights to each expert’s prediction based on the input. In our setup, the *mixture* considers an expert pool consisting of the process model (PRELES) and up to two plain ANNs (different architectures), and selects the two highest-weighted experts to form the final prediction. The experts and the gating network are trained jointly, such that the gating mechanism learns how to combine expert outputs across the input space. This design allows the model to specialise experts for different input regimes, potentially improving performance in complex or heterogeneous conditions.

All PGNN strategies, except the *residual* strategy, receive the same five input drivers as the process model (PRELES) to make their predictions. We excluded the domain adaptation and embedding strategies of Wesselkamp et al. (2024). Domain adaptation requires arbitrary choices of simulated conditions and is computationally intensive despite limited effectiveness (Wesselkamp et al., 2024). Embedding was also excluded due to slow, unstable convergence under sparse data and the computational cost of repeated joint calibration. Appendix A.1 provides further technical details and architecture specifications.

### 2.4. Experimental Setup and Evaluation

We evaluated the potential of integrating the process model with neural networks by two guiding questions:

1. How do modelling strategies perform when training data are systematically reduced, evaluated separately under in-distribution (ID) and out-of-distribution (OOD) scenarios?
2. Can changes in environmental conditions explain differences in predictive performance?

To address these questions, we designed experiments covering both ID and OOD scenarios within a systematic nested cross-validation framework (see 2.5). The ID experiments focus on data efficiency by systematically reducing the amount of training data within a single forest station. The OOD experiments evaluate the extent to which models trained on multiple forest sites can generalise to unseen (test) sites with different variable associations, and how this capacity changes with training data size. Table 2 details the experiments, including thinning strategies.

#### Effects of training size – Small & sparse data in-distribution (ID) settings

To isolate the effect of training size, we ran ID experiments at two contrasting sites: Hyytiälä (Finland), where the process model (PRELES) was originally developed, and Collelongo (Italy), a climatically divergent site. The training datasets ranged from coarse subsampling to 5 years. All models were evaluated on two test years to assess their ability to predict from limited signals. The results were used to estimate the generalisation error of each modelling strategy, and the amount of training data each model requires (details on the generalisation error in subsection 2.6).

#### Robustness & generalisation – out-of-distribution (OOD)

To evaluate model performance under OOD conditions, we tested each model’s ability to extrapolate beyond its training domain and remain robust to structural and climatic shifts, as well as data limitations. We trained each model on three forest sites, using increasing training size ranging from 22 samples to 4 years, and tested performance on a previously unseen site. This setup assesses each strategy’s capacity to generalise to novel environments not represented during training.

Together, these scenarios test both the external validity and data efficiency of process-guided neural networks (PGNNs).

### 2.5. Nested cross-validation & Hyperparameter optimisation

For each experiment, we used nested cross-validation (NCV) rather than simple random cross-validation to obtain unbiased performance estimates, thereby avoiding inflation caused by spatial autocorrelation in ecological data (Roberts et al., 2017). The NCV consists of outer and inner loops; the outer loops define the test partitions and contain hold-out data to evaluate the final model performance, while the inner loops are used for model architecture selection.

To select the model architecture in the inner loop, we performed a hyperparameter optimisation using the Optuna framework (Akiba et al., 2019) with Bayesian optimisation (Tree-structured Parzen Estimator Ozaki et al. (2020)) and Hyperband for early stopping (Li et al., 2018). Tuned hyperparameters included parameters such as learning rate, network depth and width, and the search spaces tailored to each modelling strategy (details of parameters in Table A.4). Each model was trained for a fixed number of epochs, 100 in the inner loop and 150 in the outer loop. The best five configurations from the inner loop were retrained in the outer loop on the full training set and evaluated on held-out data. This separation of model selection and evaluation ensured that observed performance reflected differences in the modelling strategy of the PGNNs rather than effects of tuning or architecture choice.

To make comparisons within experiments consistent, all nested cross-validation partitions share identical test sets, while the training-set length was varied to isolate the influence of data availability on model robustness, generalisation, and data efficiency. For each data split, we recalibrated the process model (PRELES) on the corresponding training subset to avoid information leakage. Further details are provided in Appendix D, including a schematic representation D.17.

### 2.6. Generalisation error

Inspired by Hestness et al. (2017), we estimate how increasing the amount of data in our experiments affects the prediction error of our hybrid modelling strategies. We fit the models to the data using a power-law decay

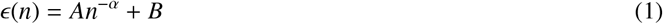

with *ɛ* the prediction error of the model, *n* the number of training data points, *α* is the learning exponent, and *B* the irreducible error, i.e., the component of the error that the model cannot reduce regardless of additional data, while *A* is the initial error of the model.

### 2.7. Evaluation metrics for model performance

#### 2.7.1. Mean Squared Error (MSE) and Root Mean Squared Error (RMSE)

For *N* data samples with observed values *Y_i_*and predicted values *Ŷ_i_*, the Mean Squared Error (MSE) and Root Mean Squared Error (RMSE) are given by

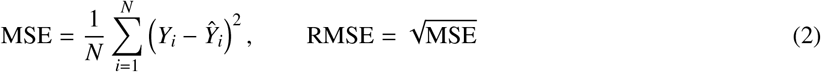

#### 2.7.2. Pearson correlation (r)

We quantified the linear association between observations and predictions using Pearson’s correlation coefficient,

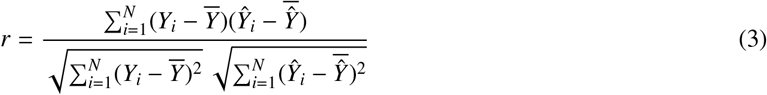

where *Y*^―^ and 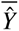 denote the means of the observed and predicted values, respectively.

#### 2.7.3. Nash–Sutcliffe model eflciency coeflcient (NSE)

Finally, the predictive skill of the models was assessed using the NSE metric (Nash and Sutcliffe, 1970), defined as one minus the ratio between the variance of the residuals and the variance of the observed data *Y*, by

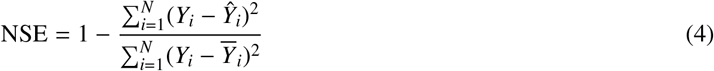

with *Y*^―^ as the mean of the observed values. An NSE value of 1 indicates perfect agreement between modelled and observed data; additionally, the NSE error can be further separated into correlation, amplitude, and mean bias (details in Appendix Appendix A.3.2).

### 2.8. Post-hoc: Model interpretability via ALE

We applied variable importance analysis using Accumulated Local Effects (ALE: Apley and Zhu, 2020) to uncover which features drove the model’s predictions. In detail, ALE measures a feature *X*_1_’s impact on model prediction by computing how small changes alter the prediction (local derivatives) for a subset of the data around the target value of *X*_1_ (hence: local) and averaged across all observed local value combinations for the other features. Those local effects are summed to form the ALE curve. Using only local observed neighbourhoods avoids unrealistic combinations of correlated features. This is particularly advantageous in our domain, where strong dependencies between variables (e.g. between PAR and GPP) are expected.

ALE was implemented using PyALE (Jomar, 2020) to quantify the main effect (*main*) between input features and outputs (GPP, ET). Furthermore, to capture the interaction strength between input feature pairs, ALE quantifies the second-order pairwise effect (*interaction*) by varying both features simultaneously. Consequently, *main* results in a 1D function, while *interaction* results in a 2D surface - both were computed using a grid size of 20. To compare feature importance between modelling strategies, we computed the range of each feature’s effect. Additionally, to keep the representation consistent across model architectures, we applied a ‘chained’ ALE calculation for the *bias* and *parallel* models: we first fed a perturbed meteorological input to the process model (PRELES), whose resulting prediction was then passed on to the hybrid architectures. This provides us with a measure of the relative importance of each environmental driver for every modelling approach. By using the outer-fold training data, we ensure the results reflect the model’s internal state.

## 3. Results

We present the results from our two experiments, an in-distribution (ID) and out-of-distribution (OOD), both of which included nested cross-validations covering a large range of training sizes (see Table 2).

### 3.1. In-distribution experiments

The ID experiments evaluated how well each modelling strategy generalised as the available training data increased at two separate forest stations, Collelongo in Italy and Hyytiälä in Finland (Figure 2).

**Figure 2:**
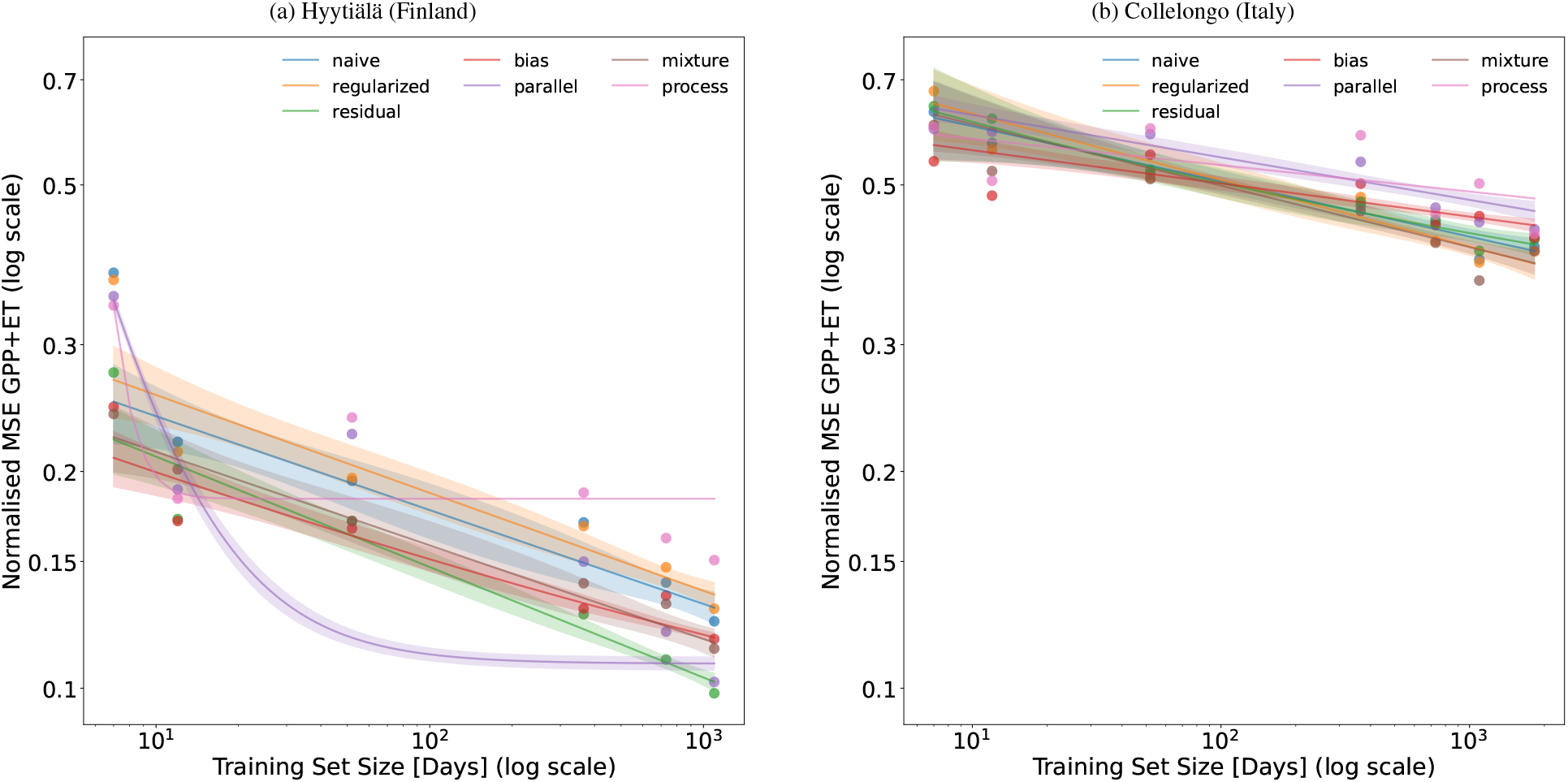
Effects of thinning on training error (MSE 1 standard error) across the best 5 configurations per strategy and fold of different-sized training sets ranging from 7 training samples to full multi-year datasets (up to 1095 samples at Hyytiälä and 1825 at Collelongo). Shading presents the uncertainty of the fit via error propagation Appendix A.4.1 ( 68% CI) excluding the baseline process model. The parameters of the generalisation error can be found in A.8.

Overall, all models improved with additional data at our test sites. Hybrid models clearly outperformed the process model (PRELES) and showed a modest but consistent advantage over the plain ANN (*naive*), although no hybrid architecture consistently performed best. The *bias* model, however, ranked second at both sites (see table A.5 for site-specific NSE rankings).

More specifically, with only seven data points, all hybrid models and the process model (PRELES) outranked the plain ANN (*naive*) at Hyytiälä. This suggests that the process model (PRELES) benefitted from its mechanistic structure under data sparsity at this site. Nevertheless, while it also improved with data, it never matched the performance gain of the *parallel*, *bias* or *residual* models, which benefitted far more strongly from additional data and pulled cleanly ahead of the plain ANN (naive). At Collelongo, performance differences were slight, with the *bias* model outperforming the plain ANN in our most sparse setting. All models showed moderate improvements as data increased, reaching very similar performances. The process model performed less well, ranking last once a weekly data size had been reached. The bias model, which relies exclusively on the process model (PRELES) predictions, matched the overall pattern of the process model (PRELES) error across training lengths (Figure 2). As data size increased, the *mixture* and *residual* models pulled ahead. This suggests that the process model struggles most to predict with noisy data.

Our most stable strategies across both sites were the *bias* and the *parallel* model (see Figure 3b, Figure 3b). However, this was not uniform: at Collelongo, the *parallel* model exhibited a few outliers, resulting in a longer tail (Figure 3b). Overall, both architectures’ dependence on the process model (PRELES) seemed to have a stabilising effect.

**Figure 3:**
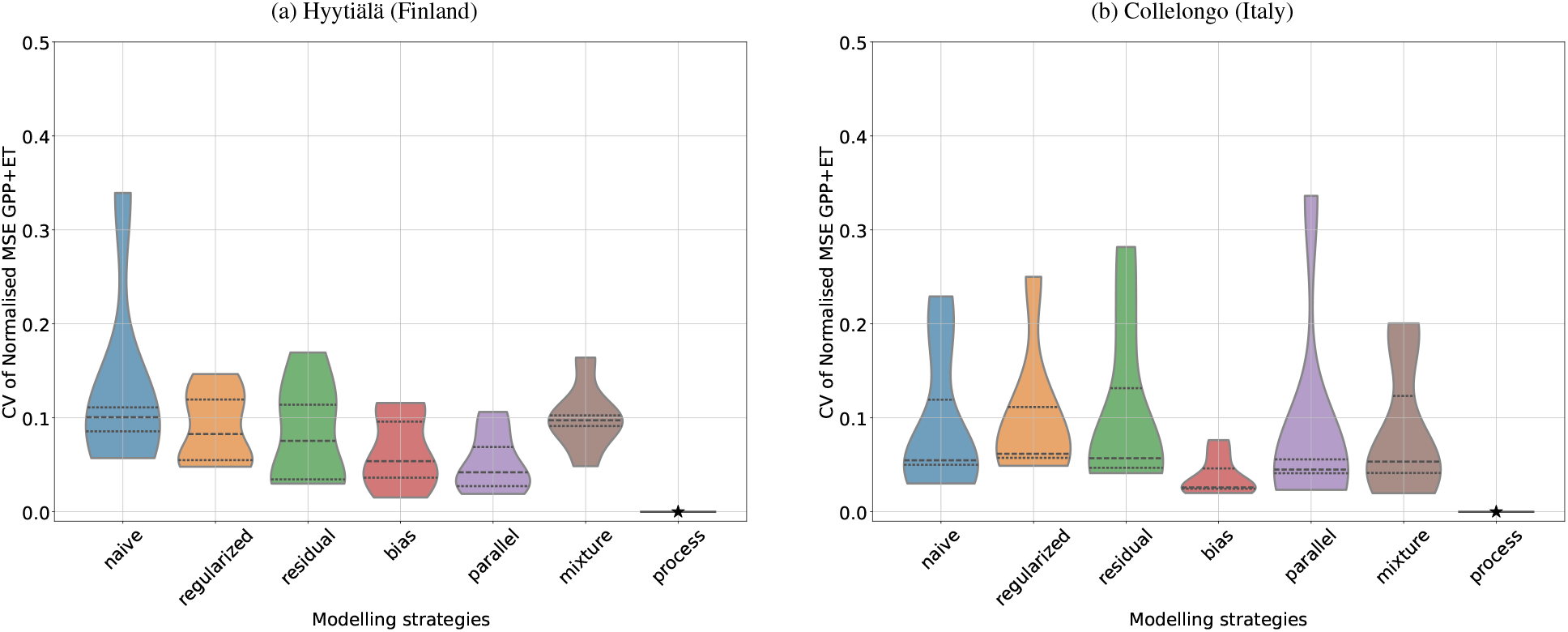
Consistency of model predictions, expressed as coefficient of variation for each modelling strategy, computed across the five different temporal holdouts of test data for every model separately. Within violin plots, the dotted and dashed lines represent the 25/75% quantile and median, respectively. Process model marked by a star.

To summarise, for the ID experiments, the hybrid strategies substantially improved over the process model (PRELES) in average error (MSE), stability (CV), and predictive skill (NSE); the **bias** model offered a consistent advantage over the plain ANN (*naive*) baseline. Addressing our first research question directly: only under extreme data limitations, with seven training samples, did one hybrid (**bias**) strategy show a clear advantage over the process model (PRELES). At Hyytiälä, the first-ranked *residual* achieved a mean NSE score of 0.84 together with the *bias* model, compared to 0.80 by the plain ANN (*naive*) model (NSE +0.04), corresponding to a 4% improvement in predictive skill and 5% over the process model (PRELES). At Collelongo, the *bias* design achieved a mean NSE of 0.51, matching the plain ANN and ranking second only to the *mixture* design (NSE 0.53). However, the data quality issues and high variability in performance across folds make this ranking less conclusive. Figure 4 shows the normalised outputs of the process, plain ANN and *bias* model against the measured GPP and ET signals, with the latter showing indications of sensor errors. We include test folds 2 and 3 in the appendix, Figures A.10a and A.10b.

**Figure 4:**
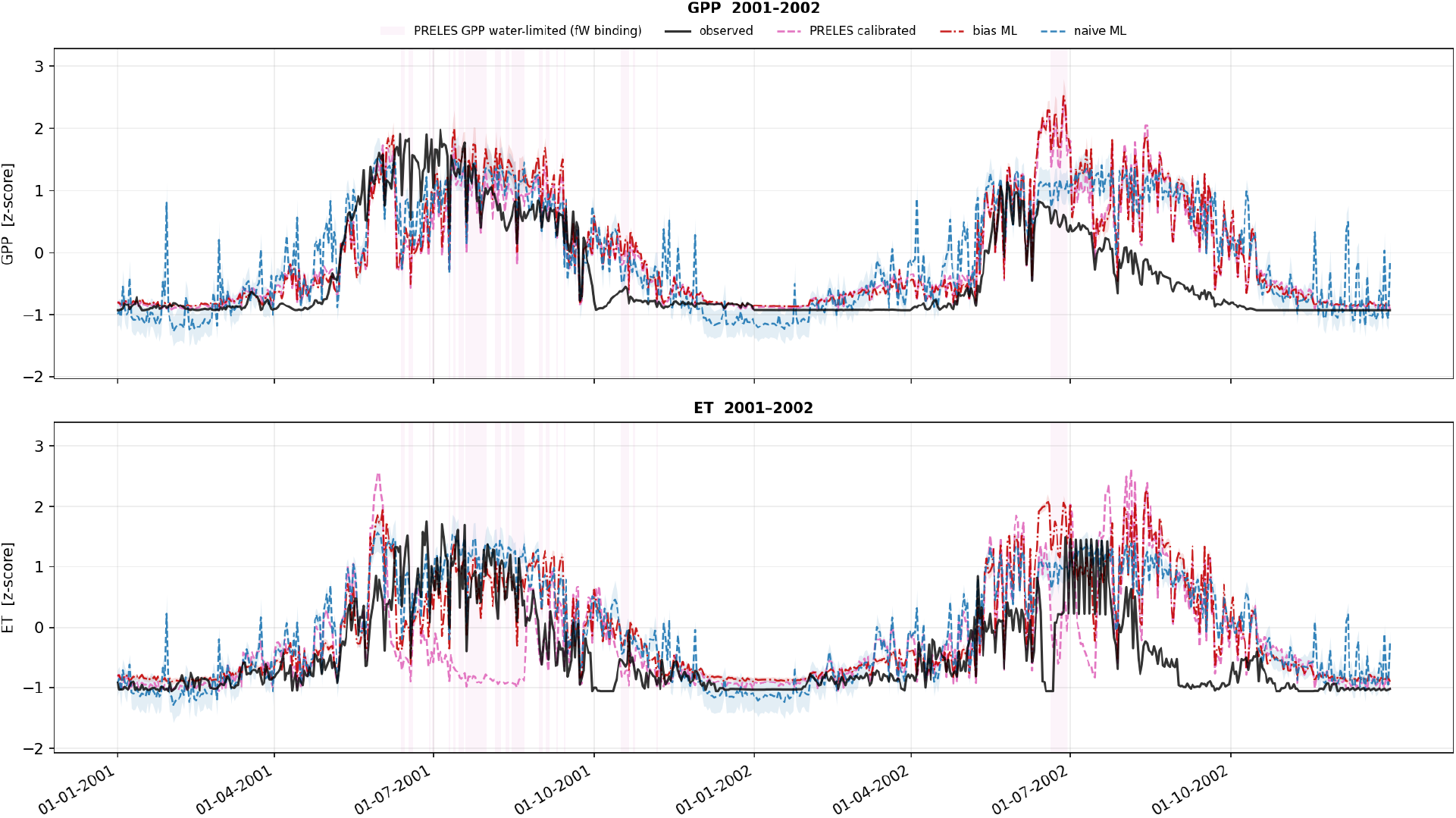
Normalised (*z*-scored) model predictions (GPP and ET) at Collelongo for process model (PRELES), post-calibration, pink), plain ANN (naive, blue) and bias (hybrid, red) models for test fold 1 (2001–2002), trained on 2% thinning data (i.e. 7 training samples). Pink shading indicates where the process model (PRELES)’s fW variable, a soil-water modifier that suppresses GPP when soil water falls below a calibrated threshold.

### 3.2. Out-of-distribution experiments

The OOD experiments assessed how well each modelling strategy extrapolates to unseen environments by training on three forest sites and testing on a fourth, previously unobserved one. Figure 5a reports these results. We again fitted a power-law decay function to the experiments (details of the resulting parameters in Table A.9)

**Figure 5.**
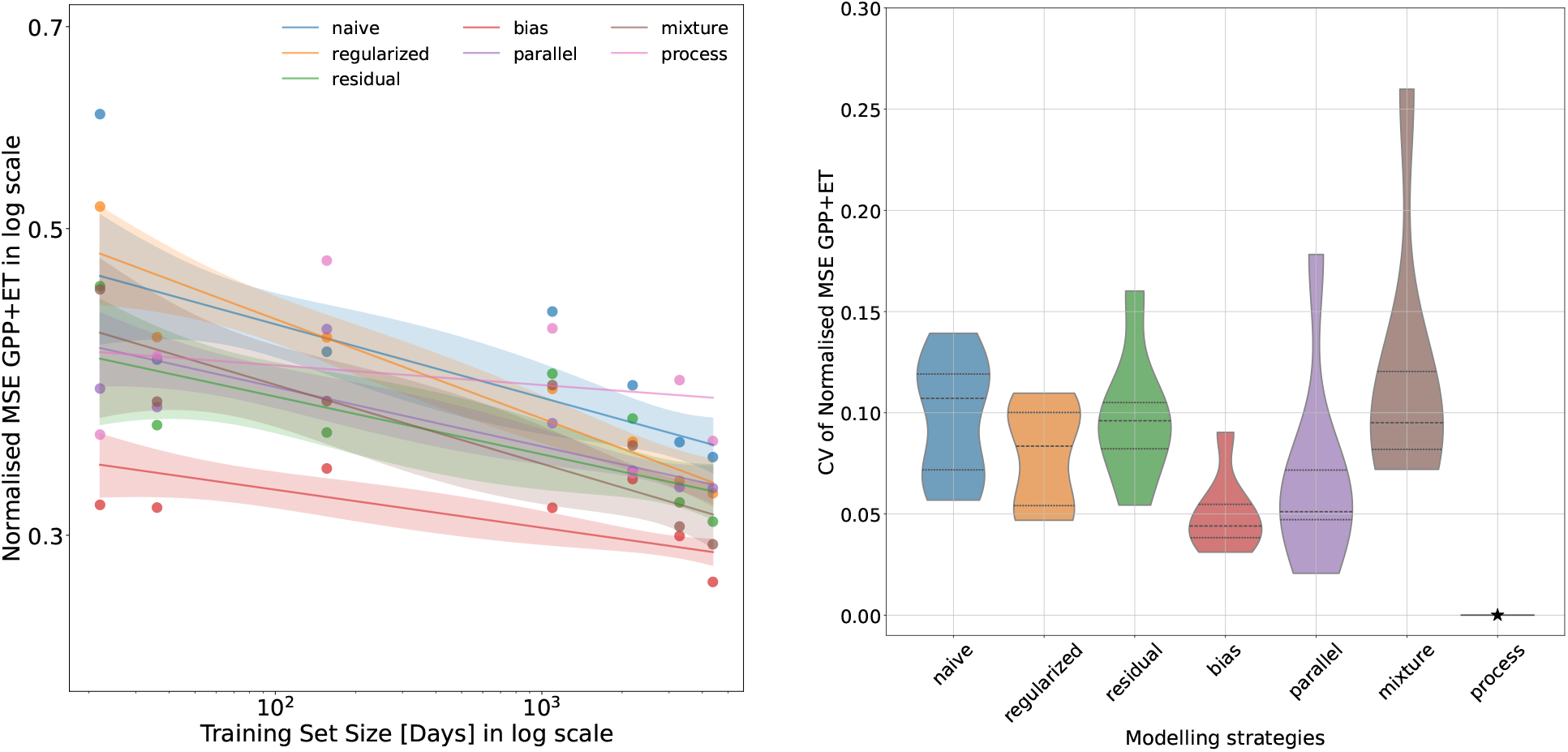
(a) Effects of thinning on test error on normalised (across stations and GPP and ET) (MSE 1 standard error) across the best 5 configurations per strategy of different-sized training sets ranging from 37 training samples to 13140 samples across 3 different training stations and one test station. Shading represents the uncertainty of the fit error propagation Appendix A.4.1 ( 68% CI), excluding the baseline process model. Details of resulting parameters in Table A.9 (b) CV for each modelling strategy is computed across the different hold-outs of test data, taking the average of the MSE and standard deviation of the 5 configurations of each modelling strategy for every hold-out of test data and dividing these averages. The process model (PRELES) has only one configuration per held-out and therefore no standard deviation. Within the violin plots, *…* the 25% and 75% quartiles and median, while the −− line the 50% quartile (median).

As with the ID experiments (Figure 3.1), almost all modelling strategies improved with more data. The *bias* model ranked first overall, highlighting the advantage of a simple non-linear correction added on top of the *process* model (PRELES), particularly when data are scarce. Averaged over training sizes, the *bias* model improved on the plain ANN by 0.11 NSE, equivalent to 11% improvement. For the smallest training size (22 samples), all hybrid models and the process model (PRELES) outperformed the plain ANN (*naive*) on average. The *process* model (PRELES) ranked second in the sparsest-data setting but was overtaken by the plain ANN (*naive*) model as data availability increased. The hybrid models similarly improved, maintaining their advantage across all data settings.

To assess whether the modelling strategies generalised consistently across test stations, we again calculated the coefficient of variation (CV) in Figure 5b. Here, the same hybrid strategies (*bias* and *parallel*) showed the lowest CVs as in the temporal experiment. The *bias* model had the lowest median, followed by the *parallel* model as a top-performing strategy, though the latter showed a large variance, followed by the regularized and *residual* models, and finally by the plain ANN (*naive*). The mixture model showed extreme levels of variability.

Finally, to compare the skill of the modelling strategies at the different test stations and under different data thinning scenarios, Figure 6 reports the average NSE across both targets. Although performance varied across strategies, the *bias* model performed best overall, ranking first at Hyytiälä (NSE 0.89), Collelongo (0.65), and Le Bray (0.55), and second at Sorø (0.68), where the *parallel* model ranked first. The *bias* model also performed consistently well under data-sparse conditions, outperforming the process model (PRELES) at all sites except Le Bray. The process model (PRELES) slightly outperformed the plain ANN at Collelongo and Le Bray, particularly where training data were scarcest. To summarise, the *bias* model performed best under OOD conditions, followed by the *parallel* and *mixture* models.

**Figure 6:**
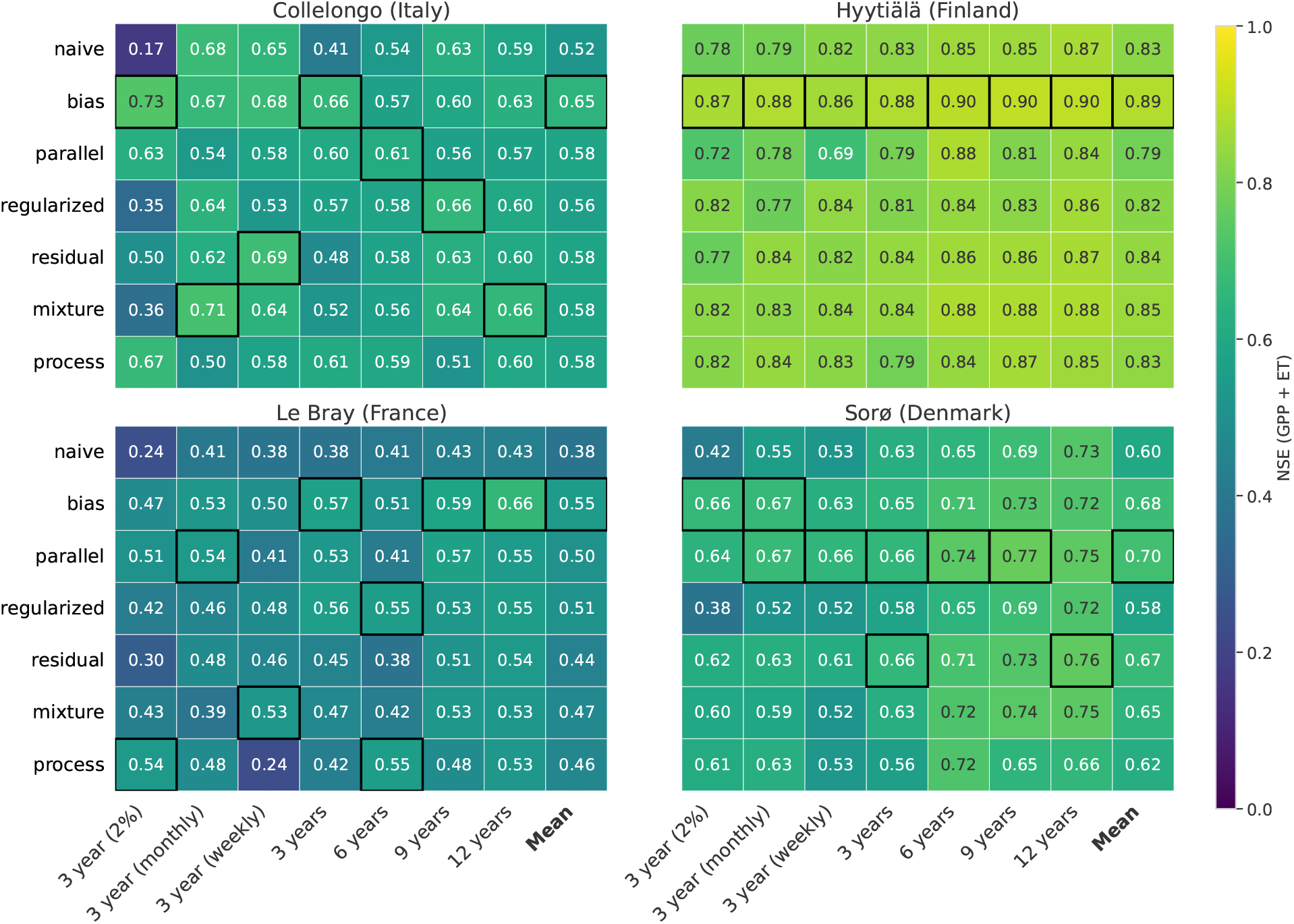
Nash-Sutcliffe efficiency (NSE) of GPP and ET combined for the different hybrid strategies for the 4 test stations sampled over different training data sizes ranging from 22 to 13140 samples. At each station, the final column summarises the mean NSE across all training sizes of every modelling strategy. NSE performance is classified as > 0.7: excellent; 0.5-0.7: good; < 0.5: poor. The best-performing strategy is highlighted by a black frame.

Notably, performance varied widely when examining the GPP and ET predictions separately. The hybrids’ advantage over the plain ANN (*naive*) came mainly from GPP, the target the process model (PRELES) predicted well. At every test site, *bias* exceeded *naive* for GPP (by 0.10–0.26 NSE) but only marginally for ET (0.01–0.06), which reflects the process model (PRELES)’s own performance. While the process model (PRELES) ranked second only to *bias* for GPP, it performed worst for ET. Additional metrics (RMSE and *r*^2^), can be found in the supplementary materials in Tables A.12, A.13.

#### Transfer and generalisation analyses

The process model (PRELES) performed worst on the training data for both GPP and ET, yet achieved among the best test performance (NSE) for GPP (see Figure A.13). In contrast, the PGNNs and the plain ANN (*naive*) showed a larger decline in performance from training to test data. Decomposing NSE into its timing (*r*), amplitude (*α*), and mean-level bias (*β*) components (Appendix A.3.2) allowed us to identify which aspects of the predictions were the most difficult for each model to transfer from training to unseen sites (see Appendix A.14). Interestingly, the sources driving performance loss differed between GPP and ET. For GPP, the neural network models (PGNNs and plain ANN) lost, on average, approximately twice as much performance skill between training and test data as the process model (PRELES) did, with most of this loss attributable to timing and mean-level bias. For ET, in contrast, the neural network models showed a smaller loss in the level component than the process model (PRELES), but a larger loss in timing.

The differences in mean-level bias provide an even clearer illustration of this pattern (see Figure 7). The process model (PRELES) consistently showed the largest deviation in GPP levels in the training data compared to all other models, yet achieved the lowest mean level bias on the GPP test set. Its more rigid structure and therefore more limited fitting to the training data may therefore contribute to the process model (PRELES)’s stronger generalisation for GPP. However, the same pattern did not extend to ET. For ET, the process model (PRELES) exhibited the largest bias error in the training data, and this remained consistent in the test set, whereas the other models performed better in both the training and test sets. This suggests that the process model (PRELES) benefits from its model structure for GPP, allowing it to generalise well despite its poorer training performance, but that this advantage does not extend to ET. This difference is also evident in the average NSE, when separated by output (see Appendix A.11)

**Figure 7:**
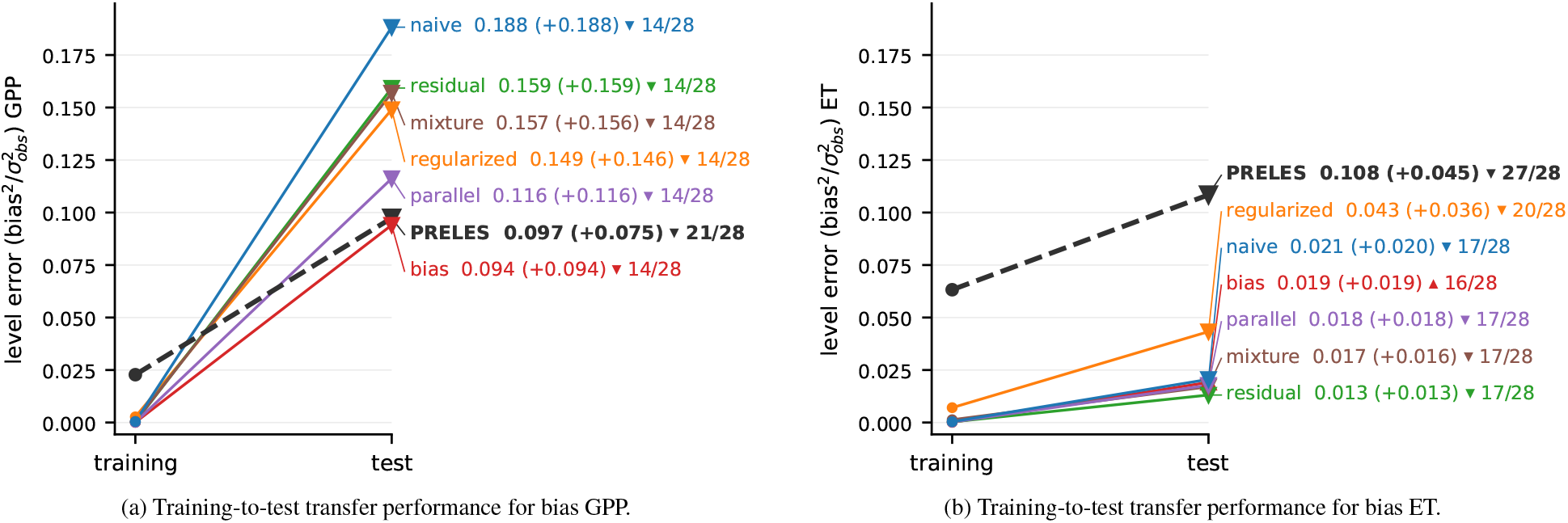
Comparison of training and test-level error (bias) pooled across all OOD experiments and training lengths. The value following each model name indicates the absolute mean bias, while the value in parentheses represents the difference between test and training bias (biastest biastrain). The marker at the test value indicates the direction of the bias: ▴ denotes overestimation at the test site, while ▾ denotes underestimation. The number following the marker indicates the number of cases with the corresponding bias direction out of the total number of cases (overestimating/underestimating / all cases).

## 4. Post-Hoc Analyses: Interpretability

We hypothesised that performance differences at test time could be linked to correlation changes of the input variables. We examined this using Accumulated Local Effects (ALE), which measures how far a fitted model’s prediction moves as that input varies; see section 2.8. This indicates how much influence a variable has on the predicted output. We found that the model architectures adapted to the heterogeneous and diverse training site distributions differently, learning to rely on various combinations of input variables. However, there were no consistent patterns to be found in this comparison that would explain model performance differences, and we refer to the figures of this test in the supplementary in Figures D.18 to D.20 (input to target interaction) and in Figures D.21 to D.23 (input to input interaction). The remainder of this section focuses on Le Bray, the site with the lowest average performance.

### 4.0.1. Diagnosing the difficulties of Le Bray

#### Predicted GPP

Le Bray was tested on the same years (2006–2008) in all experiments, and its poor NSE scores largely reflect a failure to represent Le Bray’s seasonal mean level (bias). All network strategies over-predicted GPP at Le Bray by 0.59–1.48 gC m^−2^ d^−1^ (observed mean 5.01), whereas the process model (PRELES) underpredicted it by 0.71. The correlation (*r*^2^) with observed GPP was similar for the process model (PRELES) and the hybrids ( 0.75), and lowest for naive (0.66) (Appendix table A.13).

Beyond just level errors, Le Bray also showed one anomalous period. The predictions show that the strategies diverged most during summer 2006 (Figure 8, 12-year training set). Observed GPP declined sharply from early July to mid-August and continued to decline till mid-October. The process model (PRELES) predicted a similar decline primarily through its soil-water modifier *f_W_*. *f_W_* is 1 while the model’s own simulated soil-water store stays above a calibrated threshold, then falls linearly to zero as the store approaches the calibrated wilting point. This period therefore plausibly represents a drought event that the process model (PRELES) captured. The hybrids and the plain ANN responded very differently to this period. The plain ANN (*naive*) with no access to the process model (PRELES), or information on soil water states, kept its early-summer level and over-predicted GPP throughout. The *residual* model matched the *naive* in spring but began to decline once the process model (PRELES) did, reproducing only part of the drop and predicting well above the observations. The *bias* model, whose only inputs are the process model (PRELES) outputs, followed the process model (PRELES’) suppression and predicted the observed decline most closely, briefly undershooting it in late August before over-predicting again in October, as *f_W_* returned to 1. The contrast with 2007 shows that this divergence is tied to the *f_W_* response: in 2007, when *f_W_* did not trigger, the three networks predicted similar GPP throughout. Across all training regimes, the process model (PRELES)’s accuracy in predicting this time window varied strongly, and the *bias* reproduced only the 2006 drop where the corresponding calibration did; its advantage over plain ANN (*naive*) at this site therefore derives from the process model (PRELES)’s ability to reproduce this seasonal decline.

**Figure 8:**
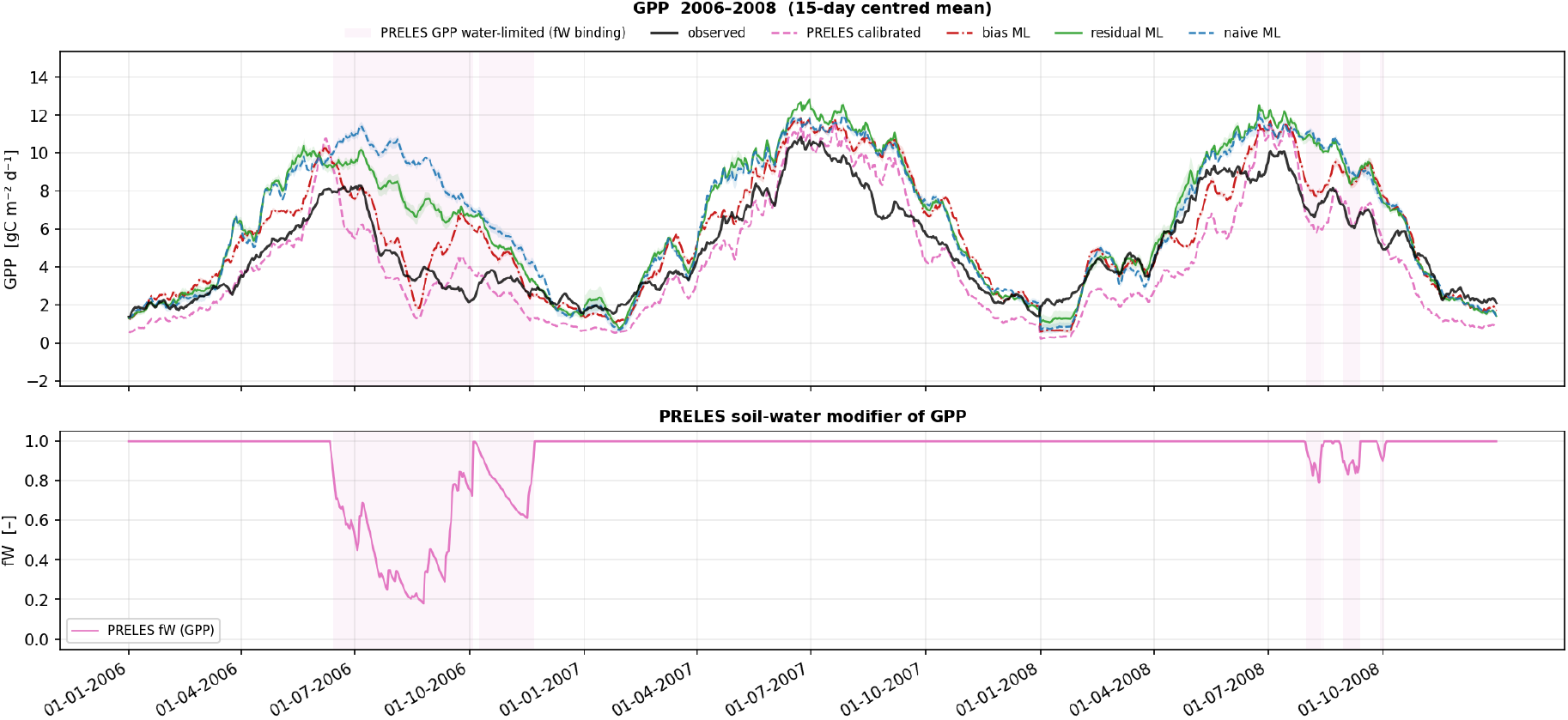
Model predictions for GPP at Le Bray: observations (black), PRELES (post-calibration, pink), plain ANN (naive, blue), *residual* (green) and bias (hybrid, red) models for test years (2006-2008. Pink shading indicates where the process model (PRELES)’s fW variable, a soil-water modifier that suppresses GPP when soil water falls below a calibrated threshold.

#### Predicted ET

ET proved more complex, showing a combination of level error and the lowest *r*^2^ correlation scores (A.13), suggesting poorer seasonal timing. Notably, all strategies reproduced Le Bray’s ET seasonal timing poorly, both as a training and as a test site. The process model (PRELES) additionally already underestimated (mean seasonal level) the ET signal of Le Bray when it was in the training data (appendix Table B.15). That all models struggle even when Le Bray is in their training data suggests that some information required to explain the ET signal is missing altogether.

To see whether the models’ internal representation of the ET drivers revealed any discrepancies, we examined the ALE responses across sites. The ALE curves revealed that the strategies differed most in how ET responds to VPD (see Figure 9). Since VPD is highly correlated with PAR, we also examined the 2nd-order ALE responses (see Appendix Figure B.16). These showed that at low PAR, VPD had little effect on predicted ET in any model. At high PAR, the networks consistently learnt that higher VPD corresponds to a lower ET, at all four test sites. In contrast, the process model (PRELES) showed opposite responses across sites: its ET prediction decreased as VPD increased in the Collelongo and Hyytiälä folds, whereas in the Le Bray and Sorø test folds, its ET prediction increased as VPD increased. The hybrids applied a correction to this wrongly signed response. The ALE curves for *bias* and *parallel*, directly reliant on the process model (PRELES) outputs as inputs, showed that the models applied corrections to the process model (PRELES) from either side: where its response was positive, they corrected it towards neutral or negative; where it was strongly negative, they weakened it slightly. The hybrids thus converged on a flat to moderately negative response, though not always fully (*parallel* in the Le Bray fold remained positive).

**Figure 9:**
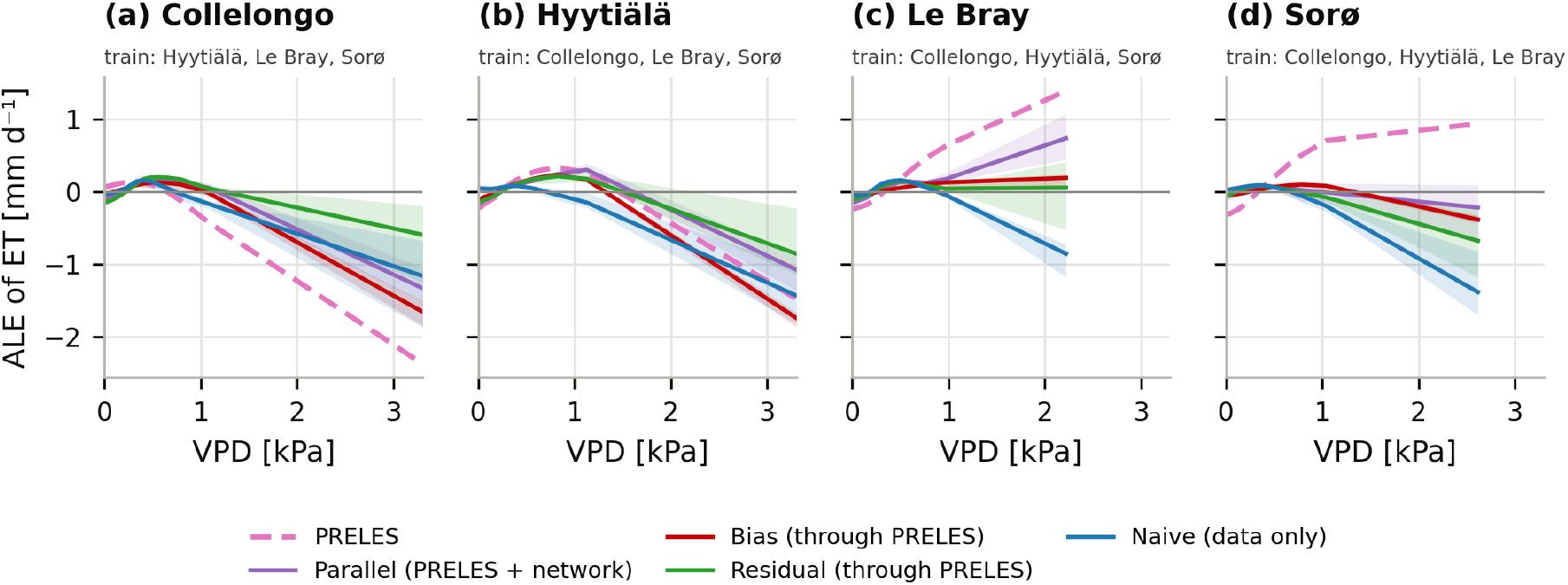
Learnt response of ET to VPD per test site, 12-year configuration: accumulated local effects (ALE) computed on the training days of each fold, mean (line) and range (band) over five configurations. Curves are centred and show how predicted ET changes with VPD, not its level. *Parallel* is the process model (PRELES) plus the network term; *residual* and *bias* include the path through PRELES. Grey: VPD reached at the test site beyond the training range.

That the process model (PRELES) fitted opposite responses suggests that calibration, rather than model structure, sets the response. This follows from how the process model (PRELES) computes ET. The process model (PRELES) model formulation does not include an ET-specific VPD response. Instead, transpiration is derived from GPP, multiplied by a water-use ratio that is calibrated once for all training sites. Transpiration is scaled up with VPD to represent the drying power of the air (evaporative demand). Soil evaporation (not VPD-dependent), is then added, resulting in the final ET prediction. VPD therefore acts on ET in two opposing ways: it lowers ET by reducing GPP, and raises it through evaporative demand.

This structure has three consequences. First, whether ET rises or falls with VPD depends on how the calibration balances these two effects. In the Le Bray and Sorø folds, the VPD point at which ET switches from rising to falling lay at or beyond the upper end of the observed range. This is the fold-dependent response made visible in the process model (PRELES) response surfaces (appendix Figure B.16), where ET continues to rise with VPD. Across training lengths, this switch point ranged from 0.3 to 8.7 kPa for the same training sites, indicating that it is poorly constrained by the training data. Second, the process model (PRELES) limits GPP according to whichever driver imposes the greater constraint: the VPD limitation or the soil-water limitation ( *f_W_* ). Once soil water becomes the stronger limitation, the VPD modifier no longer acts on GPP, correspondingly no longer acts on ET and the only remaining VPD effect is this upward scaling through evaporative demand. In effect, under soil-water limitation, the process model (PRELES)’ ET prediction can only rise with VPD, all else being equal. Finally, because the water-use ratio is shared across sites, the process model (PRELES) assumes that, at a given VPD and soil-water state, every site uses the same amount of water per unit carbon. This is unrealistic, and the observed ET-to-GPP ratio differed between sites, ranging from 0.24 at Sorø to the highest 0.42 at Le Bray, which would explain the process model (PRELES)’ ET error at Le Bray. These structural properties add to the process model (PRELES)’ difficulty in predicting ET at Le Bray, though they cannot fully explain the site’s low ET correlation, which all strategies share. The plain ANN (*naive*) still performs worst here, and does not share the mechanistic inaccuracies.

The full figures showing the ALE effect on both GPP and ET for all input variables are shown in the appendix (Figures B.14 and B.15).

## 5. Discussion

Our results demonstrate that incorporating mechanistic structure into neural networks generally improved predictive performance, particularly under out-of-distribution (OOD) conditions. In the OOD setting, all PGNN variants either outperformed or performed at a similar level to the stand-alone process model (PRELES), while the plain ANN (naive) model performed worst overall. Among the PGNN variants, the *bias* PGNN achieved the best overall performance, indicating that its formulation provided a particularly effective balance between mechanistic constraints and data-driven flexibility. Under in-distribution (ID) conditions, the rankings of the best-performing PGNNs varied across data size and sites, with only the *bias* PGNN consistently outperforming the plain ANN (*naive*). These results suggest that the mechanisms encoded in the process model (PRELES) already capture the main structure of the flux signals, and a simple non-linear correction is sufficient to improve predictions. Notably, the *bias* PGNN also showed greater robustness under data scarcity and achieved the highest predictive accuracy. Together, these findings highlight the potential of the *bias* formulation as a robust strategy when transferability and stability matter.

Compared with the previous study by Wesselkamp et al. (2024), our results show improved predictive accuracy across all modelling strategies, including the process model (PRELES) itself. These improvements likely reflect the extensive hyperparameter optimisation based on Bayesian optimisation applied to each modelling strategy, together with the calibration of the process model parameters (Van Oijen et al., 2005; Sun and Sun, 2015) within the nested cross-validation framework. As a result, observed performance differences were less influenced by suboptimal tuning and more reflective of genuine structural differences between modelling approaches.

### 5.1. Little training data suflces to capture flux dynamics

Unique to our experiments was the systematic evaluation of data size, as data scarcity is frequently identified as a key limitation in (ecological) ML applications (Von Rueden et al., 2021; Willard et al., 2022; Karpatne et al., 2024). All models were surprisingly insensitive to it when judged by correlation (*r*^2^) with observed fluxes. Across both in-distribution (ID) and out-of-distribution (OOD) experiments, roughly 40-50 samples were sufficient to achieve stable predictions. This corresponds to a weekly resolution in the ID and a monthly resolution for the OOD experiments. Errors in flux mean-bias level and amplitude behaved differently and predictions improved with more data but could not reduce them fully; we return to this below. Learning the dynamics is therefore a comparatively simple task as radiation, temperature and atmospheric dryness can describe the seasonal course of GPP and ET very well, although anomalous events, such as drought occurrences, remain difficult to predict (Williams et al., 1997; Tramontana et al., 2016; Jung et al., 2019; Mahnken et al., 2022; Camps-Valls et al., 2023). This finding applies to daily GPP and ET predicted from PAR, air temperature, VPD, precipitation and fAPAR, and we expect it to generalise for similar sets of drivers. For studies with limited observational resources, it means that sparse sampling can be enough to learn when fluxes rise and fall, provided the samples span the seasonal cycle.

### 5.2. The PGNNs are more robust under extrapolation than the plain ANN (naive)

Consistent with other (pure) machine-learning studies that predict carbon fluxes (Dou et al., 2018; Uyekawa et al., 2025), we observed a reduction in predictive accuracy at sites that are ecologically different to the training data. Under these conditions, PGNNs demonstrated their clearest advantage. Across the OOD settings, all PGNNs outperformed the plain ANN (*naive*) at nearly every test site, and that gap was widest at Le Bray, the site with the lowest overall skill. This suggests that incorporating process knowledge into machine learning models can impart information that makes them more robust to changing environmental relationships. This makes them the superior choice when transferring models across climatic zones or when future conditions are expected to deviate from the past.

Although this study focuses on forest ecosystem modelling, our results align with other studies reporting the benefits of embedding knowledge from process models into machine-learning frameworks across ecological domains. For example, Chen et al. (2022) proposed a graph-based meta-learning approach for river networks by incorporating simulated physical properties to model water quantity and quality. In agroecosystem modelling, Liu et al. (2022) used a process model to generate synthetic training data, improving estimates of N_2_O emissions under limited observational data. Yang et al. (2023) later expanded this approach by replacing the process-based model with a neural-network surrogate that represents different biophysical processes. They also used Kalman filtering and particle swarm optimisation to update states and estimate parameters for carbon dynamics and crop yield. Finally, Yu et al. (2024) pre-trained neural networks in a simulated lake environment and incorporated physical constraints, such as mass and energy conservation, to improve predictions of dissolved oxygen concentrations under changing environmental conditions.

Together with our results, these studies show that process-guided learning offers a viable strategy for improving robustness and transferability when data are sparse, systems are heterogeneous, and models must transfer across space or regime.

### 5.3. Le Bray as an example of the problems of transfer

Le Bray showed the lowest average NSE, making it the most divergent of the four test sites. Its errors illustrate three ways in which a process model’s information can transfer, or fail to transfer, to an unseen site. The process model can carry information the input drivers lack, a poorly constrained mechanism that does not help, and a consistent mean-bias level error. At Le Bray, the process model (PRELES)’ soil-water state helped the hybrids predict the 2006 drought decline in GPP; its weakly constrained ET response to VPD was partly corrected by the hybrids, and its structural underestimation of ET was corrected without giving the hybrids an advantage over the plain ANN.

#### When the process model carries information the inputs lack

During the 2006 European heatwave, the process model (PRELES) predicted a GPP decline through its soil-water modifier *f_W_* (Figure 8). This aligns with Stocker et al. (2018), who attributed Le Bray’s GPP decline to a soil-moisture-driven reduction in light-use efficiency. None of the neural networks received soil water as an input, and the plain ANN (*naive*) continued predicting the learnt seasonal trajectories. The process model (PRELES), in contrast, contains its own approximated state variable of soil moisture, and the PGNN architectures inherited the response to that state to different degrees: *bias* tracked the decline most closely, while the *parallel* and *residual* models reproduced it only partially. This state representation is what improved the hybrid’s predictive performance at this site. All hybrids predicted Le Bray’s GPP better than the plain ANN, and *bias* also outperformed the process model (PRELES) itself.

#### The hybrid models can learn to correct mechanistic structure

While hybrid models can inherit process structure that benefits them, they can also inherit structure that does not. Le Bray’s poor ET performance prompted us to examine the models’ learnt functional relationships via ALE, which revealed a weakly constrained VPD response that the hybrid models worked to correct. The process model (PRELES) has no ET-specific VPD response; ET inherits the VPD limitation set for GPP and combines it with evaporative demand. Consequently, ET’s response to VPD is a balance between two calibrated parameters. In the Le Bray fold, the calibration placed the VPD point at which ET begins to fall beyond the observed range, so ET rose with VPD over all observed conditions.

The same structure constrains the process model (PRELES) under soil water limitation. It limits GPP according to the stronger constraint, VPD or soil water. When the soil is dry, the VPD effect on GPP disappears while the drying effect of the air on transpiration remains, so drier air can only increase the process model (PRELES)’ ET. This directly contrasts with the expected response, in which ET falls with high VPD at high PAR, as stomata close (**??**Grossiord et al., 2020). The plain ANN (*naive*) learnt that ET falls with VPD at high PAR in every fold, while the hybrid models counteracted the process model (PRELES)’s positive response in the Le Bray and Sorø folds, neutralising it, except for the *parallel* model in the Le Bray fold. This shows that with sufficiently informative data, the PGNNs can override incorrect mechanisms supplied by a process model. This mechanistic inaccuracy is likely one of several contributing factors that drove the process model (PRELES)’s poor ET *r*^2^, since the plain ANN, which saw no process information, did not perform any better.

#### A consistent process-model bias can be corrected, but not exploited

Neither the meteorological inputs nor the process model (PRELES) transferred Le Bray’s flux level’s reliably; each failed for one target. Other studies using far more training data have shown this to be a persistent problem, as flux level depends on vegetation and site properties that climate alone poorly captures (Reichstein et al., 2014; Tramontana et al., 2016; Besnard et al., 2018; Migliavacca et al., 2021; Mahnken et al., 2022). For GPP, the plain ANN overestimated Le Bray’s level in every test year, not only during the 2006 drought, whereas the process model (PRELES) largely avoided this error. For ET, the roles reversed. The process model (PRELES) computes ET from GPP with ET parameters calibrated jointly on the training pool, and so assumes that all sites use the same amount of water per unit of carbon taken up. Le Bray uses considerably more, so the process model (PRELES) underestimated its ET whether the site was in the training data or not.

This is where the hybrids can capitalise on the process model, but also where they can be misled. The process model (PRELES) underestimated both fluxes at the training sites, and the hybrids learnt to correct this. For ET, PRELES’ underestimation persisted at Le Bray, and the learnt correction transferred. The hybrids roughly halved the process model (PRELES)’ ET level error, although it did not bring the hybrids closer to the observed level than the plain ANN. For GPP, the correction overshot. The process model (PRELES) underestimated Le Bray’s GPP less than that of the training sites, yet the hybrids applied a larger correction at Le Bray than in training, so that *residual* and *parallel* still overestimated the site’s GPP. Effectively, hybrids can exploit a consistent process-model level error, but only as far as both the level error and the learnt correction transfers to the new site.

### 5.4. Limitations

Several limitations of this study warrant acknowledgement. Although our four sites span a meaningful climatic gradient from boreal to Mediterranean conditions, they remain a small and exclusively European sample. Assessing PGNN transferability across a broader range of ecosystems and regions is therefore needed.

A further limitation concerns our use of gap-filled flux and meteorological products as model targets and inputs. GPP, ET, and VPD in the PROFOUND database are partly derived through MDS gap-filling from meteorological drivers similar to those used as model inputs, so we cannot fully attribute predictive skill to real ecosystem responses rather than to the gap-filling procedure itself. A post-hoc check (Appendix C.2) restricted evaluation to directly-measured days at each site. At the one site with a sufficiently large and seasonally representative directly-measured sample (Sorø), predictive skill (RMSE, NSE) showed no detectable difference, and mean bias trended toward less over-prediction on measured days, the opposite of what gap-filling circularity would predict. At the remaining three sites, the measured sample was too small and seasonally skewed to support this comparison; we cannot rule out some influence of gap-filling structure there.

Our fAPAR preprocessing carries a related uncertainty. The rolling-window length and notch threshold used for outlier detection were selected by visual inspection rather than analytical validation. Our sensitivity analysis ( Appendix C.1) shows that plausible alternative parameters reclassify a meaningful fraction of flagged values in both directions and shift the reconstructed series. Therefore, we cannot say which setting most accurately reflects the true signal; we report this as an unresolved source of input uncertainty.

For comparability, all models used the same predictors as the process model, excluding potentially informative variables such as soil moisture, nitrogen availability, and atmospheric CO_2_ concentrations. Additionally, the process model itself showed limited and partly unreliable performance across sites; other state-of-the-art forest stand-scale process models may provide a stronger mechanistic backbone and improve the inductive bias available to hybrid strategies.

Lastly, we deliberately used relatively simple multi-layer perceptrons due to the data constraints tested here. More sophisticated architectures, such as Transformer-based models, may provide additional benefits when larger training datasets are available. As such, our results should not be interpreted as a general assessment of the performance limits.

## 6. Conclusion and Outlook

Process-guided neural networks represent a promising modelling strategy for ecological prediction under data scarcity. Across four European forest types, PGNNs achieved competitive predictive performance with remarkably limited training data. The main findings of this study can be summarised as follows:

1. PGNNs consistently outperformed the plain ANN (*naive*) and generally outperformed the stand-alone process model (PRELES), except for the GPP prediction at Le Bray under out-of-distribution conditions. These results suggest that PGNNs benefit from both the learning capacity of neural networks and the process-based knowledge encoded in the process model (PRELES).
2. PGNNs, specifically the *bias* PGNN, provided the greatest advantage for in-distribution and out-of-distribution settings on average, demonstrating the potential of process-guided learning across a range of environmental settings.
3. Incorporating process-based knowledge improved transferability to climatically novel sites, suggesting that mechanistic structure can provide a useful inductive bias when extrapolating beyond the training domain.
4. ALE-based analyses indicated that both PGNNs and ANNs learnt physically plausible relationships between environmental drivers and predicted fluxes. Combined with the other analyses, these results suggest that PGNNs benefit from the process-based guidance of the process model (PRELES) during drought conditions - information that is not available from the input drivers alone. However, they also inherit some of PRELES’s physically implausible constraints on evapotranspiration (ET), although they can partially correct these limitations.

Despite these advantages, transitions between energy-limited and water-limited regimes remain a persistent challenge. Near these transitions, no model configuration performed consistently well. However, PGNNs that closely followed the process model (PRELES) water-stress modifier ( *f_W_* ) benefited from this process-based state variable. At the same time, PGNNs partially corrected the physically implausible ET constraints inherited from the process model (PRELES), highlighting both the benefits and limitations of process guidance.

To address these limitations, we envision two options: First, more detailed mechanistic process models may provide stronger and more transferable inductive biases for PGNNs. The process model (PRELES), encodes empirical approximations of mechanistic structure tailored to specific forest ecosystems. In multi-site contexts under unfamiliar environmental regimes, not all assumptions may hold. Incorporating more complex or mechanistically rich process models provides stronger and more transferable inductive biases, potentially enhancing structural generalisation, and could further increase the performance of PGNNs.

Second, some regime transitions may be difficult to represent explicitly within fixed process formulations, but could potentially be learned from data. Recent hybrid modelling work (Fang and Gentine, 2024) has shown that selectively replacing uncertain process components, such as ET-specific functions, with simple neural representations can substantially improve site-level predictions of carbon and water fluxes without increasing overall model complexity.

Future hybrid approaches could therefore benefit from explicitly targeting regime-dependent processes through improved representations of water limitation, more flexible parametrisations of uncertain process components, and training data covering a wider range of environmental conditions. Furthermore, the constraints imposed by process-based knowledge could be relaxed selectively for individual outputs. In particular, given the limitations of the process model (PRELES) in predicting ET, allowing more independent ET predictions — for example, by relaxing the process model (PRELES)-based constraints in the regularised loss function — may enable PGNNs to better exploit the learning capacity of neural networks while retaining the benefits of process guidance for other outputs.

## 7. Code Availability

The entire workflow was implemented in Python and PyTorch, allowing for a flexible adaptation to new models and data. The code https://github.com/hannah-habenicht/Process-Guided-Neural-Networks-in-Forest-Carbon-Flux-Modelling and additional data can be found in https://zenodo.org/records/23942881.

## 8. Acknowledgement

Funded by the Deutsche Forschungsgemeinschaft (DFG, German Research Foundation) – Project-ID 499552394 – CRC 1597 “SmallData”. CFD acknowledges partial funding by the German Excellence Initiative Project number 533786343 - FutureForests (EXC 3127), and the DFG Project-ID 459819582 - CRC 1537 “Ecosense”. This research was funded by the Deutsche Forschungsgemeinschaft (DFG, German Research Foundation) under grant number 539134284, through EFRE (FEIH_2698644), and the state of Baden-Württemberg.

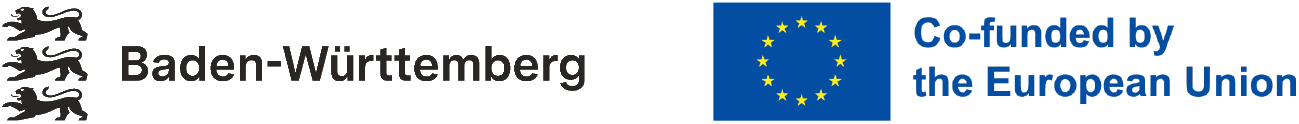

During the preparation of this work, the authors used Claude (Anthropic) for language editing. After using this tool, the authors reviewed and edited as needed and take full responsibility for the content of the published article.

## Appendix A. Modelling strategies and Experiments

### Appendix A.1. Modelling strategies

All architectures optimise a supervised prediction objective based on either mean-squared error (MSE) or mean-absolute error (MAE). We write the neural component as a parametric mapping *f_θ_* : *X* → *Y* from environmental drivers *x* ∊ *X* to predicted fluxes. See Figure 1 for a schematic representation.

The *naive* network minimises the difference between its prediction *ŷ*_NN_ = *f_θ_*(*x*) and the observed target *y*,

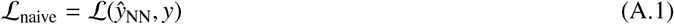

The *bias* and *residual* strategies apply the same loss but differ in how the neural network interacts with the process-model output *ŷ*_process_. The bias model predicts *ŷ*_bias_ by correcting the output of *ŷ*_process_

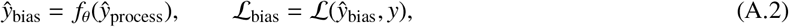

whereas the residual model receives *ŷ*_process_ as an additional input and learns the difference between its predictions *ŷ*_res_ and the true values *y*.

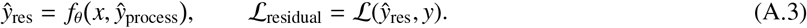

The *parallel* model, has the same inputs as *x* as the *naive* network, but adds the output of the process model *ŷ*_process_ to the output of the neural network,

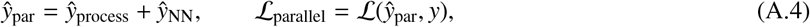

such that the physical prediction augments the learned term.

The *regularized* model adds an explicit penalty that discourages deviations from the process-model behaviour,

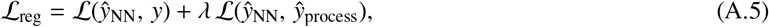

with *λ* controlling the strength of the physics-based constraint.

Finally, the *mixture model* (Shazeer et al., 2017) combines several trainable neural networks with a fixed pre-calibrated process model (PRELES) as experts *e*. A sparsely-gated mixture-of-experts layer *G_e_*(*x*) selects a subset of experts and learns their weights *G_e_*(*x*) based on the input *x*. So the final output *ŷ*_mix_ is produced by

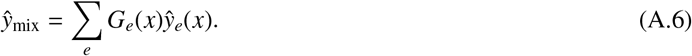

We adopt a noisy Top-K Gating mechanism: only the best performing k experts (via KeepTopK) are selected, noise is added to *H*(*x*) to encourage a balanced use of experts. The gating is finally defined as

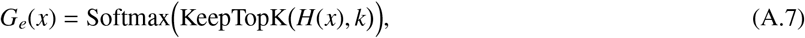

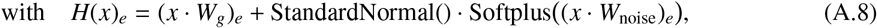

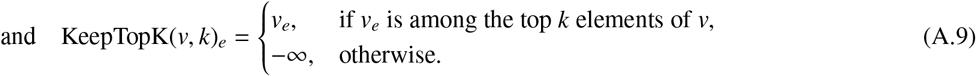

Training minimises the L_total_, which combines *ŷ*_mix_ and an entropy term *E*, that encourages the model to focus on a single expert. The entropy is computed over the expert logits *p_e_*(*x*) = Softmax(*x* · *W_g_*), averaged across each training batch *B*. In practice, we set *α* ∈ [0.005, 0.02], making the entropy term’s influence small.

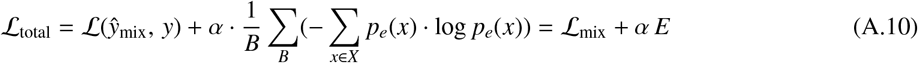

To integrate the process model (PRELES) as a pre-calibrated model, we gradually schedule the integration using a combination of noise term and hard masking at the beginning. This allows the neural network sufficient time to learn before fully incorporating the process model.

**Table A.3:**
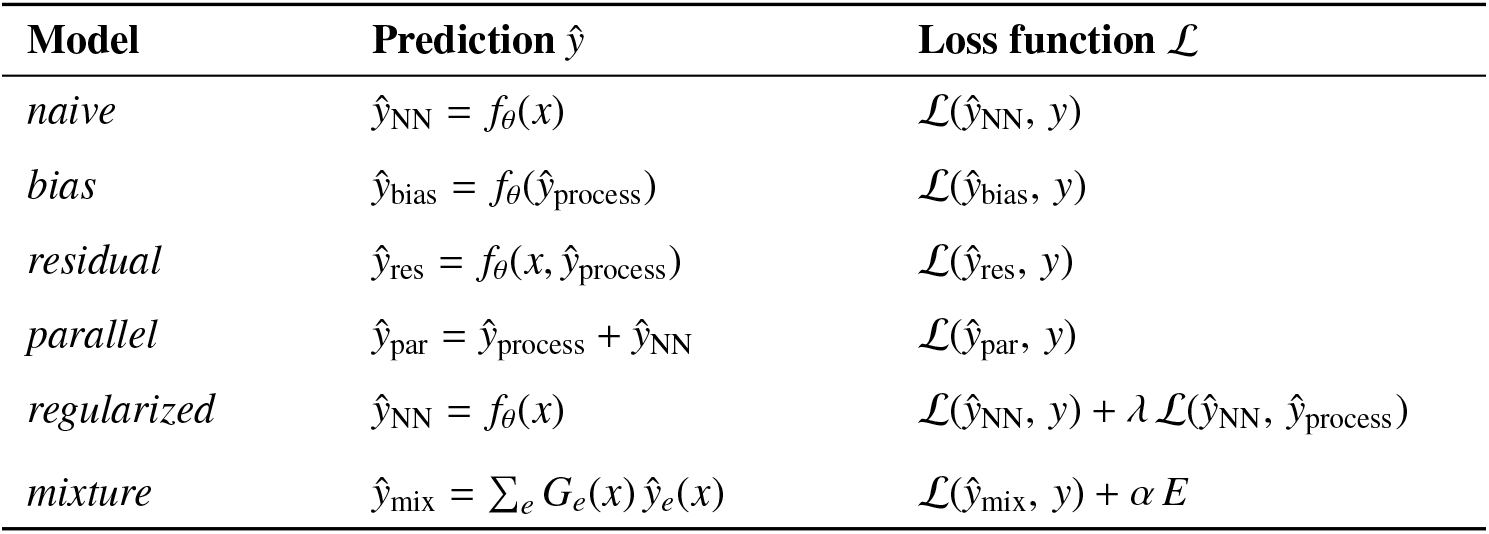
Summary of modelling strategies and corresponding prediction and loss formulations.

| **Model** | **Prediction ŷ** | **Loss function ℒ** |
|---|---|---|
| *naive* | ŷ<sub>NN</sub> = f<sub>θ</sub>(x) | ℒ(ŷ<sub>NN</sub>, y) |
| *bias* | ŷ<sub>bias</sub> = f<sub>θ</sub>(ŷ<sub>process</sub>) | ℒ(ŷ<sub>bias</sub>, y) |
| *residual* | ŷ<sub>res</sub> = f<sub>θ</sub>(x, ŷ<sub>process</sub>) | ℒ(ŷ<sub>res</sub>, y) |
| *parallel* | ŷ<sub>par</sub> = ŷ<sub>process</sub> + ŷ<sub>NN</sub> | ℒ(ŷ<sub>par</sub>, y) |
| *regularized* | ŷ<sub>NN</sub> = f<sub>θ</sub>(x) | ℒ(ŷ<sub>NN</sub>, y) + λ ℒ(ŷ<sub>NN</sub>, ŷ<sub>process</sub>) |
| *mixture* | ŷ<sub>mix</sub> = ∑<sub>e</sub> G<sub>e</sub>(x) ŷ<sub>e</sub>(x) | ℒ(ŷ<sub>mix</sub>, y) + α E |

### Appendix A.2. Hyperparameter Optimisation

Table A.4 provides an overview of all HPO settings used in the nested cross-validation experiments. It lists the shared optimisation logic, the experiment-specific search spaces for the temporal, spatial, and noisy-mixture setups, and the fixed defaults that apply across all configurations. This table serves as a reference for the exact parameter ranges and training constraints used in the study.

#### Appendix A.2.1. Process model (PRELES): Code Integration, Extension and Interface

The original process model (PRELES) was implemented in C++ by the authors Minunno et al. (2016) and made available in R through the Rpreles repository. The developed C++ Python binding was extended from Linux to include Windows and wraps the process model as a callable package. This interface supports integration of the process model (PRELES) within neural networks and allows for seamless coupling with PyTorch-based training and calibration routines.

#### Appendix A.2.2. Process model (PRELES): Calibration

To calibrate the process model (PRELES), we employed a Bayesian optimisation using the Optuna framework (Akiba et al., 2019) with a covariance matrix adaptation evolution strategy (CMA-ES) sampler (Hansen, 2016; Nomura and Shibata, 2024). The model was calibrated against observed gross primary production (GPP) and evapotranspiration (ET) data by minimising the negative log-likelihood of a joint multivariate normal distribution over the residuals. In our experiments, we focused on tuning a subset of site-specific parameters related to water retention and evapotranspiration processes, including soildepth, beta, Smax, and ETkappa, among others (parameter indices 5–11, 14–18, and 31). These 12 parameters were selected for their known influence on site-to-site variability and their physiological relevance in forest ecosystem functioning. Calibration was initialised from a default set for conifer trees taken from the original repository Appendix A.2.1. For each parameter set proposed by the optimiser, PRELES predictions were generated across all training samples from a split (see NCV Appendix D), and the residuals were used to estimate an empirical covariance matrix capturing cross-process dependencies. The resulting log-likelihood served as the scalar objective function for optimisation. The calibration procedure was repeated over 800 trials, with intermediate results stored and the best-performing parameter set selected based on the minimum loss.

**Table A.4:**
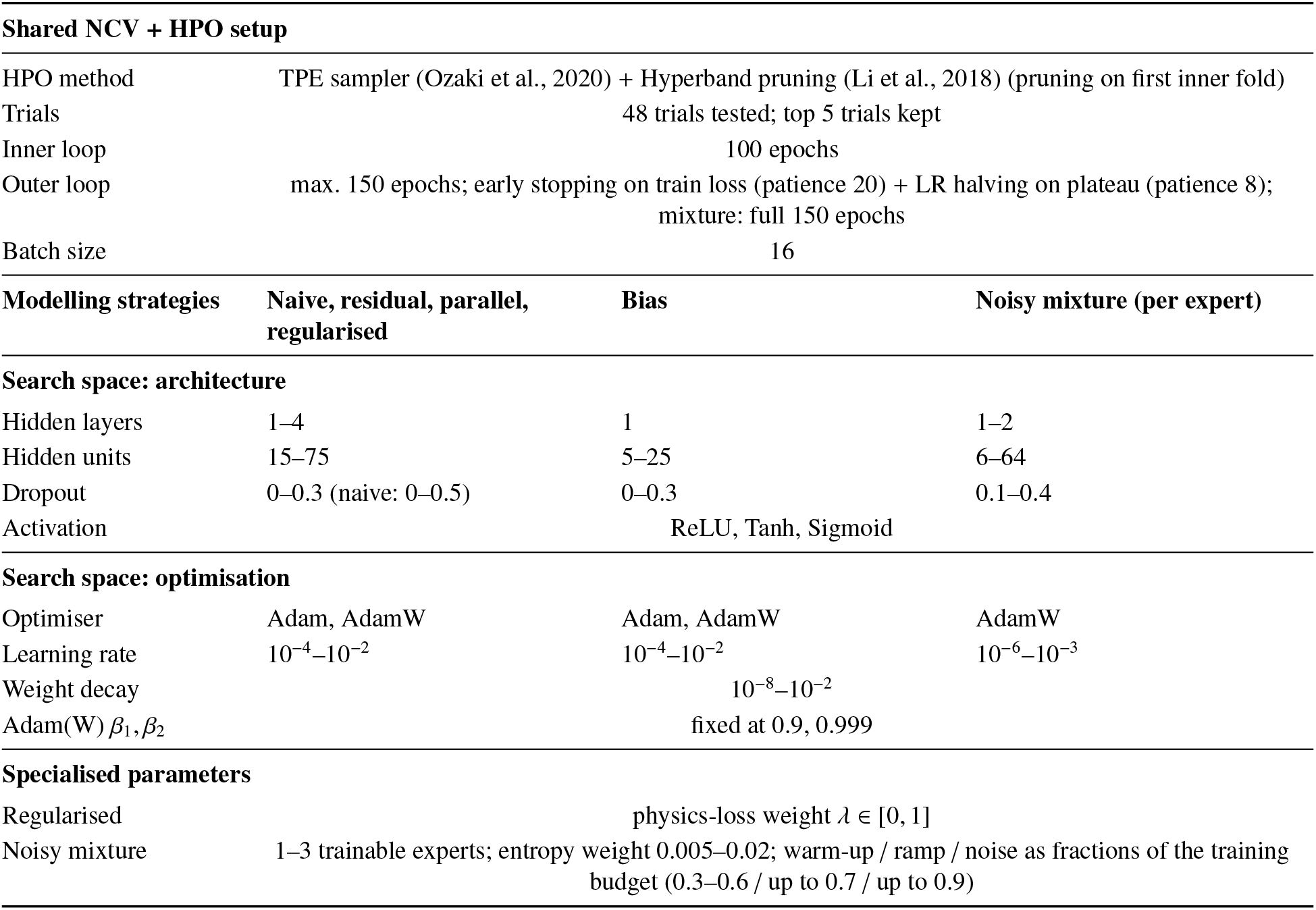
Summary of hyperparameter optimisation (HPO) settings used in the nested cross-validation (NCV) experiments. The search space is identical for the temporal (ID) and spatial (OOD) NCV; shared settings are listed once.

### Appendix A.3. Qualitative results of selected years and station

Figures 4, A.10a and A.10b shows an exemplary comparison between the process model (PRELES) and the best-performing modelling strategy, regularized, at Collelongo at the most thinned-out scenario. For both folds, irregularities are evident in the observed ET data during 2001–2002, in 2004–2005 and in 2006-2007.

#### Appendix A.3.1. In-distribution NSE results

**Figure A.10:**
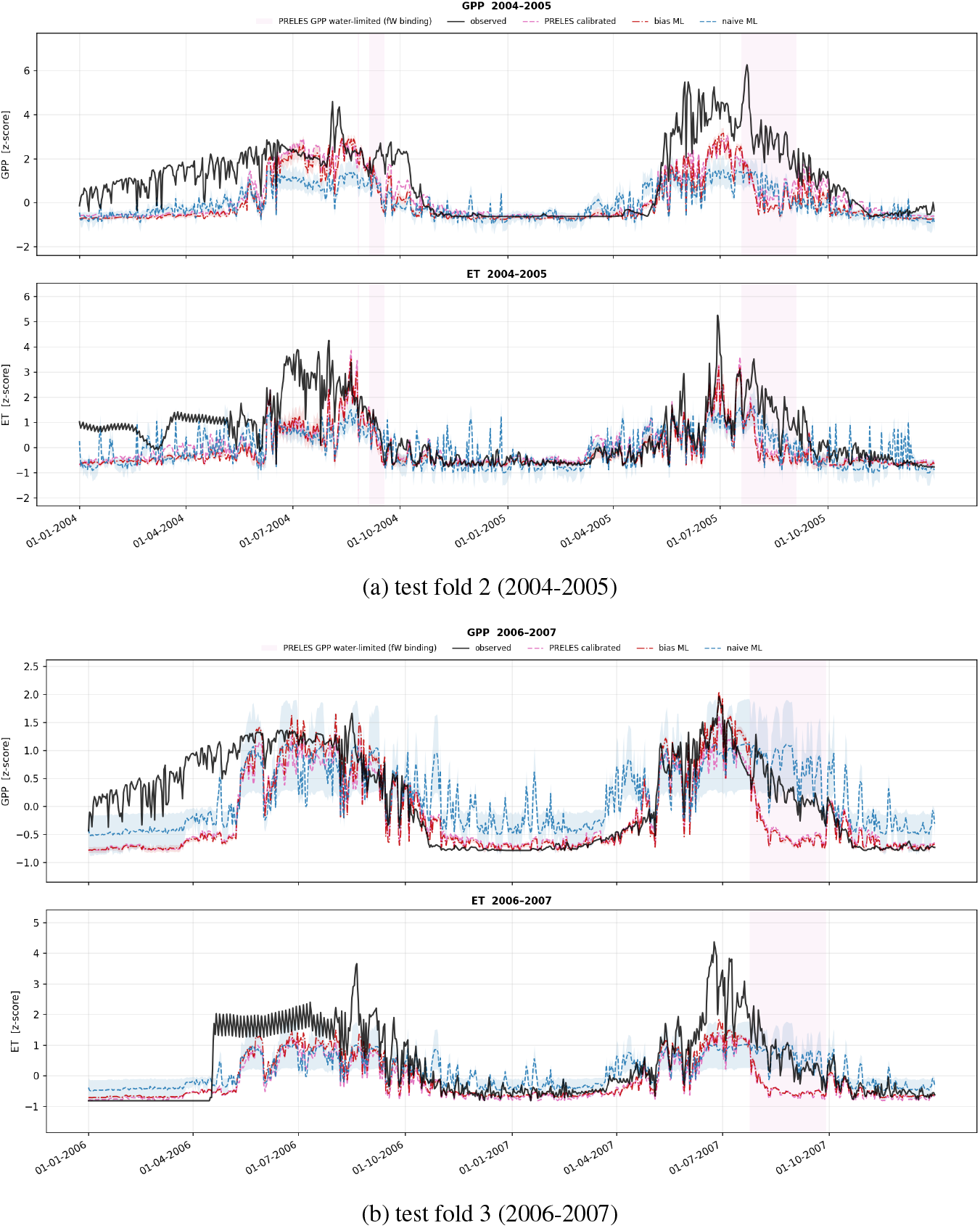
Normalised (*z*-scored) model predictions (GPP and ET) at Collelongo for the process model (PRELES) (post-calibration, pink), plain ANN (naive, blue) and bias (hybrid, red) models for test fold 2 (2004-2005) and test fold 3 (2006-2007), trained on 2% thinning. Pink shading indicates where PRELES’s fW variable, a soil-water modifier that suppresses GPP when soil water falls below a calibrated threshold.

**Table A.5:** Temporal experiments: test **NSE (Nash–Sutcli**ff**e e**ffi**ciency)** on held-out years per prediction strategy, for GPP, ET and their average (GPP+ET). NSE performance is classified as *>* 0.7: excellent; 0.5-0.7: good; *<* 0.5: poor. The five HPO configurations are averaged within each test-year fold and the folds within each data-amount regime (one training year thinned to every 50th, 30th or 7th day, and full daily data with 1, 2, 3 or 5 training years; 5 years only for Collelongo); cells report mean std over those regimes. GPP+ET is the mean of the GPP and ET NSE within each regime. Strategies are ranked separately for each station by the GPP+ET mean NSE; the highest mean per station and column is in bold. Average: mean ± std over the seven strategies.

| Collelongo (Italy) (7 regimes) |  |  |  |  | Hyytiälä (Finland) (6 regimes) |  |  |  |  |
|---|---|---|---|---|---|---|---|---|---|
| Rank | Strategy | GPP | ET | GPP+ET | Rank | Strategy | GPP | ET | GPP+ET |
| 1 | mixture | **0.52** ± **0.06** | **0.54** ± **0.10** | **0.53** ± **0.08** | 1 | residual | **0.85** ± **0.06** | 0.83 ± 0.08 | **0.84** ± **0.06** |
| 2 | bias | 0.50 ± 0.04 | 0.53 ± 0.07 | 0.51 ± 0.05 | 2 | bias | 0.84 ± 0.04 | **0.84** ± **0.06** | 0.84 ± 0.05 |
| 3 | naive | 0.50 ± 0.08 | 0.52 ± 0.09 | 0.51 ± 0.09 | 3 | mixture | 0.84 ± 0.04 | 0.82 ± 0.06 | 0.83 ± 0.05 |
| 4 | regularized | 0.50 ± 0.12 | 0.52 ± 0.09 | 0.51 ± 0.10 | 4 | parallel | 0.82 ± 0.08 | 0.80 ± 0.11 | 0.81 ± 0.09 |
| 5 | residual | 0.47 ± 0.10 | 0.52 ± 0.10 | 0.49 ± 0.09 | 5 | regularized | 0.80 ± 0.08 | 0.79 ± 0.10 | 0.80 ± 0.09 |
| 6 | parallel | 0.48 ± 0.06 | 0.48 ± 0.11 | 0.48 ± 0.07 | 6 | naive | 0.80 ± 0.09 | 0.79 ± 0.09 | 0.80 ± 0.09 |
| 7 | process | 0.51 ± 0.06 | 0.44 ± 0.11 | 0.47 ± 0.07 | 7 | process | 0.80 ± 0.06 | 0.77 ± 0.09 | 0.79 ± 0.07 |
|  | Average | 0.50 ± 0.02 | 0.51 ± 0.03 | 0.50 ± 0.02 |  | Average | 0.82 ± 0.02 | 0.81 ± 0.02 | 0.82 ± 0.02 |

**Table A.6:**
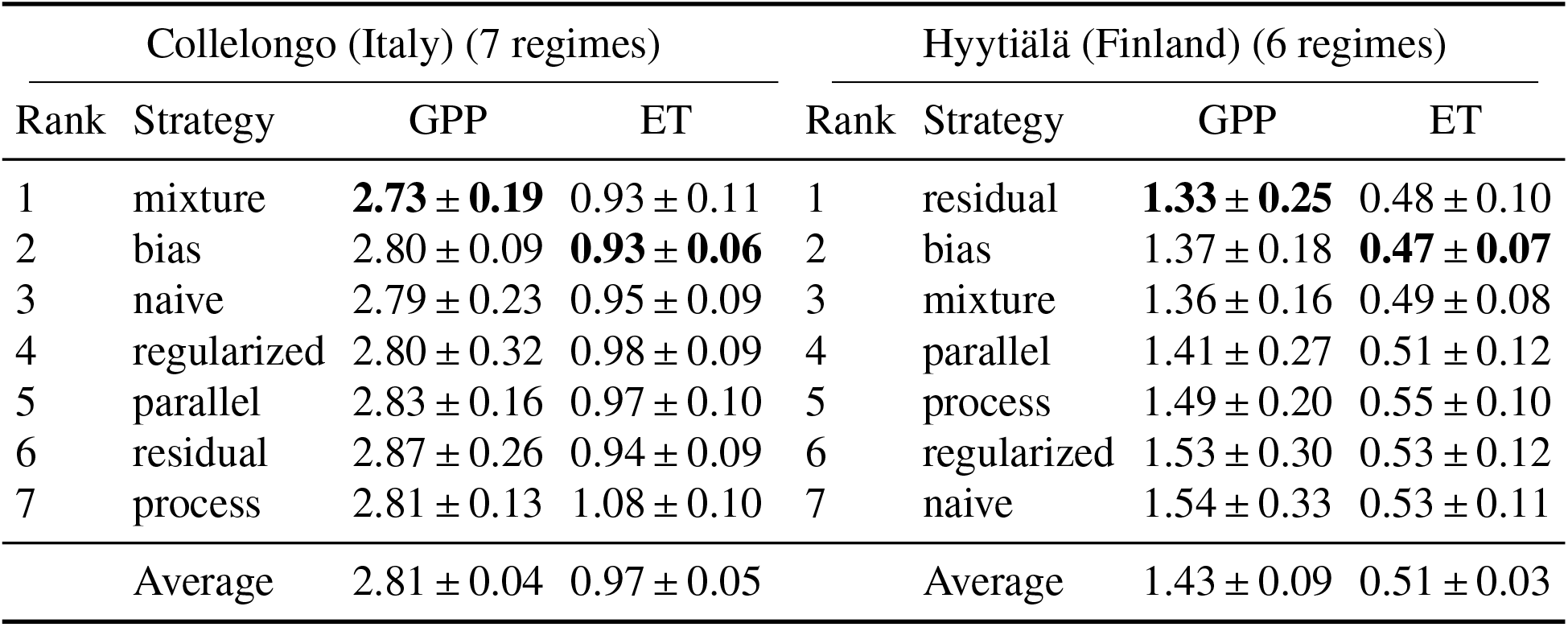
Temporal experiments: test **RMSE** on held-out years per prediction strategy for GPP (g C m^−2^ d^−1^) and ET (mm d^−1^). The five HPO configurations are averaged within each test-year fold and the folds within each data-amount regime (one training year thinned to every 50th, 30th or 7th day, and full daily data with 1, 2, 3 or 5 training years; 5 years only for Collelongo); cells report mean std over those regimes. Strategies are ranked separately for each station by the combined (GPP+ET) mean RMSE; lowest mean per station and target is in bold.

**Table A.7:** Temporal experiments: test *r*^2^ on held-out years (squared Pearson correlation between observed and predicted) per prediction strategy, for GPP and ET. *r*^2^ captures pattern/phase agreement only and ignores bias and scale errors (cf. NSE). The five HPO configurations are averaged within each test-year fold and the folds within each data-amount regime (one training year thinned to every 50th, 30th or 7th day, and full daily data with 1, 2, 3 or 5 training years; 5 years only for Collelongo); cells report mean std over those regimes. Strategies are ranked separately for each station by the combined (GPP+ET) mean *r*^2^; highest mean per station and target is in bold.

| Collelongo (Italy) (7 regimes) |  |  |  | Hyytiälä (Finland) (6 regimes) |  |  |  |
| Rank | Strategy | GPP | ET | Rank | Strategy | GPP | ET |
|---|---|---|---|---|---|---|---|
| 1 | mixture | **0.70** ± **0.02** | **0.70** ± **0.05** | 1 | bias | 0.89 ± 0.02 | **0.88** ± **0.04** |
| 2 | bias | 0.68 ± 0.01 | 0.70 ± 0.02 | 2 | residual | 0.90 ± 0.03 | 0.87 ± 0.06 |
| 3 | naive | 0.68 ± 0.04 | 0.69 ± 0.04 | 3 | mixture | **0.90** ± **0.02** | 0.86 ± 0.06 |
| 4 | regularized | 0.68 ± 0.05 | 0.69 ± 0.04 | 4 | parallel | 0.89 ± 0.03 | 0.85 ± 0.08 |
| 5 | residual | 0.67 ± 0.04 | 0.69 ± 0.04 | 5 | process | 0.88 ± 0.02 | 0.84 ± 0.08 |
| 6 | parallel | 0.67 ± 0.03 | 0.68 ± 0.04 | 6 | naive | 0.87 ± 0.05 | 0.85 ± 0.06 |
| 7 | process | 0.67 ± 0.01 | 0.64 ± 0.05 | 7 | regularized | 0.87 ± 0.05 | 0.84 ± 0.09 |
|  | Average | 0.68 ± 0.01 | 0.68 ± 0.02 |  | Average | 0.89 ± 0.01 | 0.86 ± 0.01 |

#### Appendix A.3.2. Decomposition of NSE

To separate errors in timing from errors in scale, we decomposed NSE into three components (Murphy, 1988; Gupta et al., 2009):

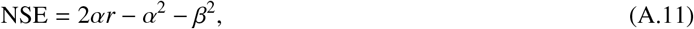

where *r* is the Pearson correlation between predicted and observed values (timing), *α* = *σ_Ŷ_ /σ_Y_*the ratio of their standard deviations (amplitude), and *β* = |*µ_Ŷ_ µ_Y_*| */σ_Y_* the mean bias scaled by the observed standard deviation (level). A perfect prediction has *r* = 1, *α* = 1 and *β* = 0.

### Appendix A.4. Generalisation error of in-distribution (ID) and out-of-distribution (OOD) estimations

Following the generalisation error *ɛ*(*n*) = *An*^−^*^α^* + *B* introduced in section 2.6, we report the estimated parameters for the ID experiments at Collelongo and Hyytiälä (Table A.8), as well as for the out-of-distribution experiment (Table A.9). Since the process model (PRELES) relies on a single fitted model, whereas the other modelling strategies are averaged over five different configurations, the resulting estimates are not directly comparable.

In the ID experiments, Collelongo and Hyytiälä, the initial error *A* of the hybrid strategies is of similar size within each station and is lower at Hyytiälä (*A* 0.2–0.4) than at Collelongo (*A* 0.5–0.8). At Collelongo, the *residual* strategy has the lowest initial error and the highest learning rate (*α* = 0.16), but it is also the only strategy with a non-zero asymptotic error (*B* = 0.26); the other strategies learn slowly (*α* 0.04–0.09), and the large uncertainties of *A* and *B* show that their error curves are nearly flat, so the initial and the asymptotic error cannot be separated well. At Hyytiälä, the *bias* strategy starts from the lowest initial error (*A* = 0.22) and approaches a small asymptotic error (*B* = 0.05), and the *naive*, *regularized*, *residual* and *mixture* strategies show similar, well-determined learning curves (*α* 0.13–0.18, *B* 0). Overall, all hybrid strategies benefit only moderately from additional training data (*α <* 0.2). An exception is the *parallel* strategy at Hyytiälä, which starts from a much larger initial error (*A* = 5.46) and compensates with a steep learning rate (*α* = 1.61), but retains an asymptotic error of *B* = 0.11, indicating a strong dependence on data volume. At Collelongo, the *parallel* strategy behaves like the other strategies. Finally, the power-law fit of the *process* model is unstable at both stations, because its error hardly changes with the training-set size. At Hyytiälä, it has the highest and well-determined asymptotic error (*B* = 0.18). Although not directly comparable to the statistical models, this behaviour suggests structural bias in parts of the process model that cannot be mitigated by additional data.

In the OOD experiments (Table A.9), all strategies show, on average across the four stations, a similar and only weakly decreasing generalisation error: the initial errors are comparable (*A* 0.37–0.57), the learning rates are low (*α* 0.01–0.07), and the asymptotic error *B* is close to zero but with uncertainties larger than the initial error. Over the range of training-set sizes considered, additional data therefore reduce the error at an unseen station only slightly, and the power law cannot separate the initial from the asymptotic error. The *bias* strategy has the lowest average initial error (*A* = 0.37), the *regularized* strategy the highest (*A* = 0.57). The fit of the process model (PRELES) is unstable on average, because its error hardly changes with the training-set size. The per-station fits (Table A.10) show that this average combines different behaviour. At Collelongo, the *naive*, *regularized* and *residual* strategies drop steeply at the smallest training sets (*α* 4, *A* at the bound of the fit) and then level off at a high asymptotic error (*B* 0.35–0.39), whereas the *bias* strategy and the process model (PRELES) reach lower floors (*B* = 0.24 and 0.28) with well-determined fits; the *mixture* fit is undetermined at this station. At Hyytiälä and Sorø, all hybrid strategies show smooth learning curves with low learning rates (*α* 0.05–0.22) and no detectable asymptotic error. At Le Bray, the *naive* strategy retains the highest asymptotic error (*B* = 0.55), followed by the *mixture* (*B* = 0.47, with an unstable *A*) and *parallel* (*B* = 0.37) strategies, while the *residual* fit is unstable. Also, the *bias* here has a comparatively large fitted initial error (*A* = 0.75) but keeps slowly improving with additional data (*α* = 0.09, no detectable asymptotic error) and reaches the lowest error of all strategies at this station for the largest training set (nMSE = 0.34), mainly through a reduced GPP error. The process model is the only strategy whose fit is unstable at three of the four stations (Hyytiälä, Le Bray, Sorø), because its calibration barely benefits from more training data; at Collelongo, its error is nearly constant at *B* = 0.28. Although not directly comparable to the other models, this suggests a structural limit of the process model that cannot be overcome with additional data, while the hybrid strategies improve only slowly with more data when transferred to unseen stations.

**Table A.8:** Parameters of the generalisation error of the modelling strategies for the in-distribution experiments at Collelongo and Hyytiälä. The test error is fitted as a power law of the training-set size *D*, *E*(*D*) = *A D*^−^*^α^* + *B*, with *B* the error floor. *E* is the normalised mean squared error (nMSE): for GPP and ET separately, the MSE on the held-out test years is divided by the variance of the observations of the same test years, and the two values are averaged 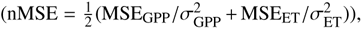 so that both fluxes contribute equally and the error is unitless (nMSE = 1 corresponds to predicting the mean of the test period). Possible artefacts or unstable estimates (*σ >* 5 and *σ >* 3 |value|) are marked in italics.

| In-distribution experiments |  |  |  |  |  |  |
|---|---|---|---|---|---|---|
|  | Collelongo |  |  | Hyytiälä |  |  |
| Strategy | *A* ± *σ* | *α* ± *σ* | *B* ± *σ* | *A* ± *σ* | *α* ± *σ* | *B* ± *σ* |
| naive | 0.72 ± 1.479 | 0.08 ± 0.269 | 0.00 ± 1.659 | 0.32 ± 0.315 | 0.13 ± 0.326 | 0.00 ± 0.423 |
| regularized | 0.78 ± 1.186 | 0.09 ± 0.259 | 0.00 ± 1.359 | 0.35 ± 0.227 | 0.14 ± 0.243 | 0.00 ± 0.317 |
| residual | 0.51 ± 0.203 | 0.16 ± 0.345 | 0.26 ± 0.444 | 0.30 ± 0.089 | 0.15 ± 0.162 | 0.00 ± 0.145 |
| bias | 0.62 ± 2.138 | 0.05 ± 0.207 | 0.00 ± 2.201 | 0.22 ± 0.041 | 0.18 ± 0.168 | 0.05 ± 0.085 |
| parallel | 0.72 ± 1.553 | 0.06 ± 0.177 | 0.00 ± 1.616 | 5.46 ± 1.585 | 1.61 ± 0.145 | 0.11 ± 0.002 |
| mixture | 0.74 ± 0.932 | 0.09 ± 0.203 | 0.00 ± 1.089 | 0.29 ± 0.247 | 0.13 ± 0.282 | 0.00 ± 0.328 |
| process | *0.63 ± 14.485* | 0.04 ± 1.014 | *0.00 ± 14.675* | *1.0e5 ± 4.03e6* | *6.87 ± 20.733* | 0.18 ± 0.019 |

**Table A.9:**
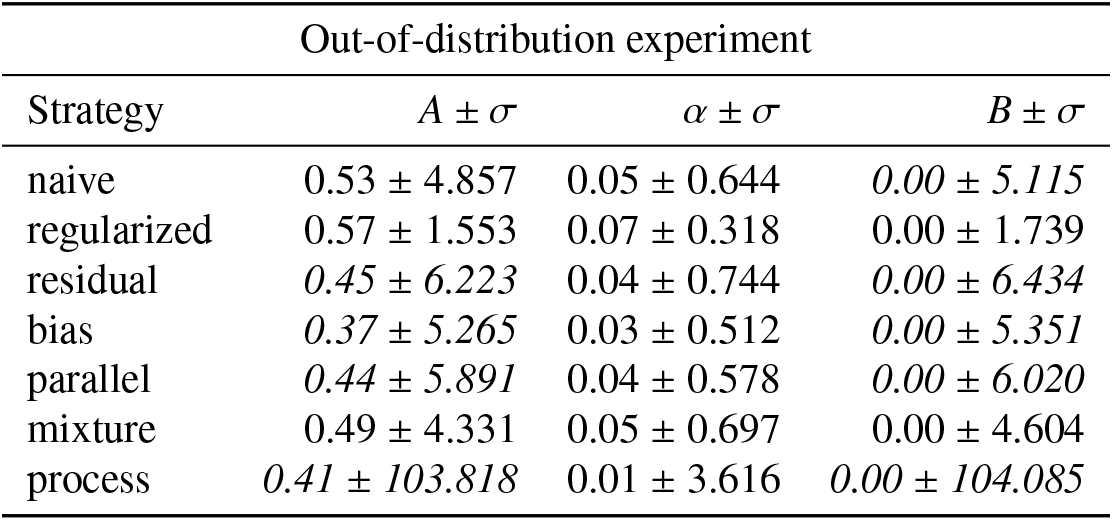
Parameters of the generalisation error for the OOD experiment, averaged across all four forest stations. The test error is fitted as a power law of the training-set size *D*, *E*(*D*) = *A D*^−^*^α^* + *B*, with *B* the error floor. *E* is the normalised mean squared error (nMSE): for GPP and ET separately, the MSE at the held-out station is divided by the variance of that station’s observations over its test years, and the two values are averaged 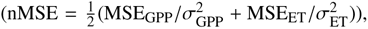 so that both fluxes and all stations contribute equally and the error is unitless (nMSE = 1 corresponds to predicting the station’s mean of the test period). Possible artefacts or unstable estimates (*σ >* 5 and *σ >* 3 value ) are marked in italics.

**Table A.10:** Parameters of the generalisation error for the OOD experiments at the four stations (each station held out in turn). The test error is fitted as a power law of the training-set size *D*, *E*(*D*) = *A D*^−^*^α^* + *B*, with *B* the error floor. *E* is the normalised mean squared error (nMSE): for GPP and ET separately, the MSE at the held-out station is divided by the variance of that station’s observations over its test years, and the two values are averaged 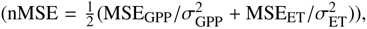 so that both fluxes contribute equally and the error is unitless (nMSE = 1 corresponds to predicting the station’s mean of the test period). Large values are given in scientific notation; possible artefacts or unstable estimates (*σ >* 5 and *σ >* 3 |value|, or *A*at the upper bound of the fit) are marked in italics.

| Out-of-distribution experiments |  |  |  |  |  |  |
|---|---|---|---|---|---|---|
|  | Collelongo |  |  | Hyytiälä |  |  |
| Strategy | *A* ± *σ* | *α* ± *σ* | *B* ± *σ* | *A* ± *σ* | *α* ± *σ* | *B* ± *σ* |
| naive | *1.00e5* ± *1.28e6* | 4.47 ± 3.90 | 0.35 ± 0.01 | 0.31 ± 0.37 | 0.09 ± 0.24 | 0.00 ± 0.46 |
| regularized | *1.00e5* ± *6.52e5* | 4.18 ± 2.10 | 0.39 ± 0.01 | 0.23 ± 1.22 | 0.06 ± 0.44 | 0.00 ± 1.30 |
| residual | *6.53e4* ± *1.58e6* | 4.21 ± 7.83 | 0.36 ± 0.01 | 0.23 ± 1.43 | 0.06 ± 0.59 | 0.00 ± 1.55 |
| bias | 0.11 ± 0.02 | 0.00 ± 0.04 | 0.24 ± 0.02 | 0.22 ± 0.07 | 0.10 ± 0.09 | 0.00 ± 0.10 |
| parallel | 0.21 ± 0.10 | 0.16 ± 0.61 | 0.33 ± 0.31 | 0.96 ± 0.18 | 0.22 ± 0.10 | 0.00 ± 0.10 |
| mixture | *0.00* ± *7.69e6* | *0.00* ± *2.14e3* | *0.38* ± *7.69e6* | 0.23 ± 1.03 | 0.08 ± 0.60 | 0.00 ± 1.17 |
| process | 0.13 ± 0.04 | 0.00 ± 0.10 | 0.28 ± 0.04 | *0.19* ± *18.66* | 0.02 ± 2.57 | *0.00* ± *18.81* |
|  | Le Bray |  |  | Sorø |  |  |
| Strategy | *A* ± *σ* | *α* ± *σ* | *B* ± *σ* | *A* ± *σ* | *α* ± *σ* | *B* ± *σ* |
| naive | 0.37 ± 1.09 | 0.34 ± 1.14 | 0.55 ± 0.15 | 0.88 ± 0.25 | 0.13 ± 0.16 | 0.00 ± 0.48 |
| regularized | 0.41 ± 0.42 | 0.11 ± 0.32 | 0.28 ± 0.62 | 1.00 ± 0.09 | 0.15 ± 0.15 | 0.00 ± 0.38 |
| residual | *0.65* ± *9.68* | 0.04 ± 0.69 | *0.00* ± *9.92* | 0.51 ± 0.99 | 0.08 ± 0.28 | 0.00 ± 1.12 |
| bias | 0.75 ± 0.47 | 0.09 ± 0.12 | 0.00 ± 0.58 | 0.42 ± 1.25 | 0.05 ± 0.21 | 0.00 ± 1.31 |
| parallel | 0.14 ± 0.02 | 0.00 ± 0.04 | 0.37 ± 0.02 | 0.45 ± 0.47 | 0.07 ± 0.13 | 0.00 ± 0.53 |
| mixture | *65.39* ± *486.62* | 1.91 ± 2.26 | 0.47 ± 0.02 | 0.53 ± 0.82 | 0.08 ± 0.24 | 0.00 ± 0.94 |
| process | *0.59* ± *197.13* | *0.01* ± *5.35* | *0.00* ± *197.77* | *0.45* ± *24.86* | 0.03 ± 1.99 | *0.00* ± *25.22* |

#### Appendix A.4.1. Error propagation by delta method

We approximated the variance *σ*^2^ of the generalisation error function *ɛ_θ_*(*n*) = *An*^−^*^α^* + *B* (Tables A.9 and A.10) using the delta method (Oehlert, 1992). For the generalisation error function *ɛ*(*θ̂*), the gradient is evaluated at the optimised parameter values *θ̂*.

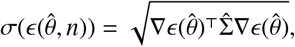

where *θ̂* = (*Â, B̂, α̂*)^⊤^, and hence ∇*ɛ* = (*n*^−^*^α̂^*, 1, − ln(*n*)*Ân*^−^*^α̂^*)^⊤^.

#### Appendix A.5. Additional results for OOD experiments

Tables A.11, A.12, and A.13 report NSE, RMSE, and *r*^2^ for the same four stations, and read together they show RMSE ranking the stations differently from the other two metrics. Hyytiälä holds the strongest position on every measure: the lowest mean RMSE (GPP 1.80, ET 0.52, averaged across strategies), the highest NSE (0.76/0.80), and the best *r*^2^ for most strategies. Collelongo sits in the middle throughout, with mid-range RMSE (GPP 2.41) and the second-lowest NSE (0.66/0.62).

Le Bray and Sorø invert this trend for GPP. Le Bray shows the lowest mean GPP RMSE of the four stations (1.76), yet its GPP NSE (0.61) is the worst. Sorø shows the reverse pairing: its mean GPP RMSE (3.01) is the highest by a wide margin, while its GPP NSE (0.69) ranks second-best. The two stations swap places between the absolute and the relative metric.

The process model’s *r*^2^ diverges from its NSE and RMSE rank at two of the four stations. At Le Bray it shows the best GPP and ET *r*^2^ of any strategy (0.75 and 0.71), and at Hyytiälä it matches the *residual* for the best GPP *r*^2^ (0.89). At Collelongo the pattern reverses: process *r*^2^ is among the weakest of any strategy, particularly for ET (0.50, the lowest score at that station), and at Sorø its ET *r*^2^ (0.74) trails the hybrid strategies by a similar margin.

**Figure A.11:**
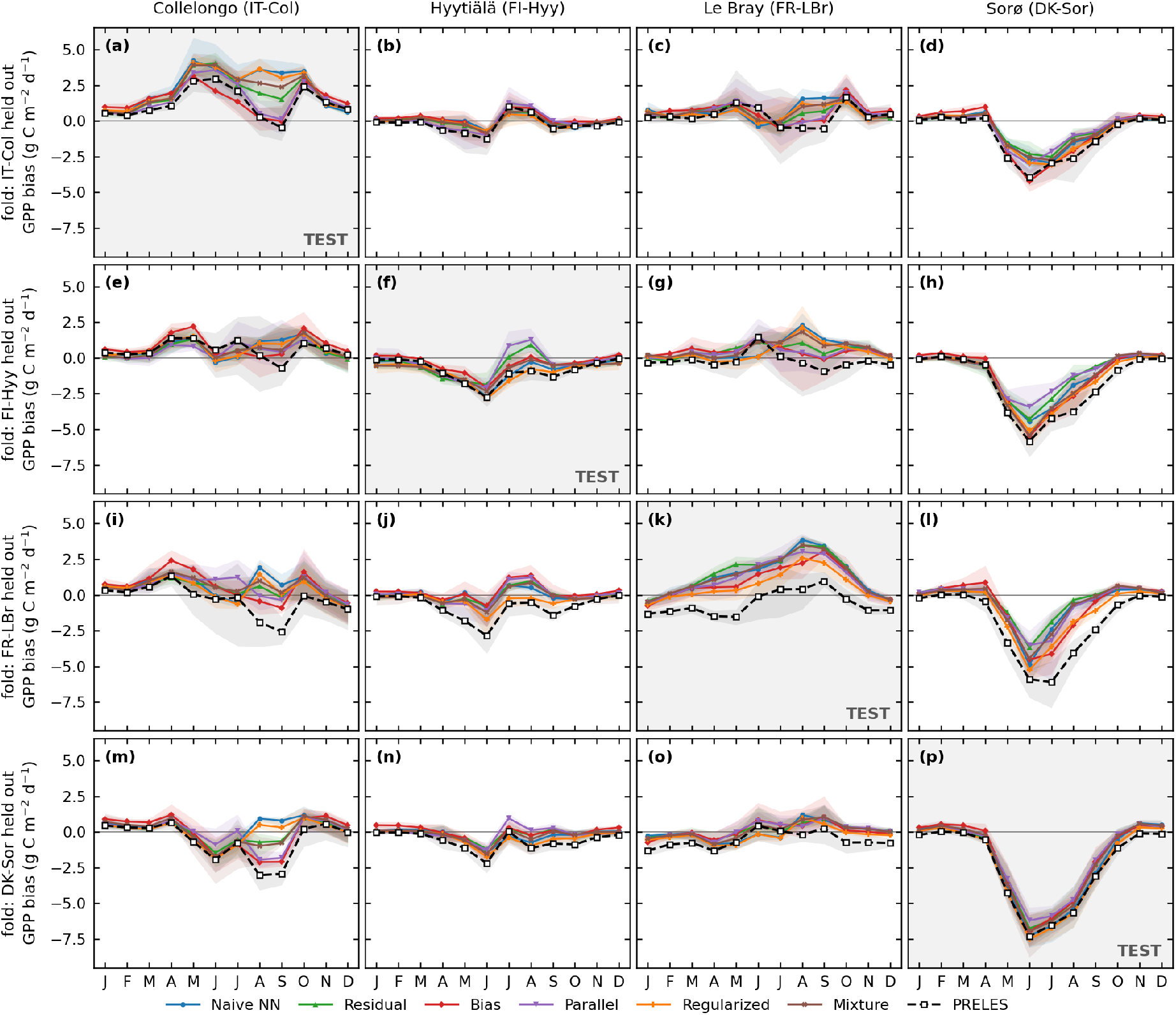
Monthly bias of daily **GPP** for all four held-out stations of the spatial NCV across full training lengths shown for training and test (rows: Collelongo 2011–2013, Hyytiälä 2010–2012, Le Bray 2006–2008, Soro 2006–2008). Bias = mean(pred - obs) per calendar month, pooled over each station’s three test years; positive values indicate over-prediction. For each NN variant, the line is the mean over five HPO configurations and the band spans min–max across configurations; PRELES (dashed) is a single calibrated run. Note the differing y-axis ranges between rows.

**Figure A.12:**
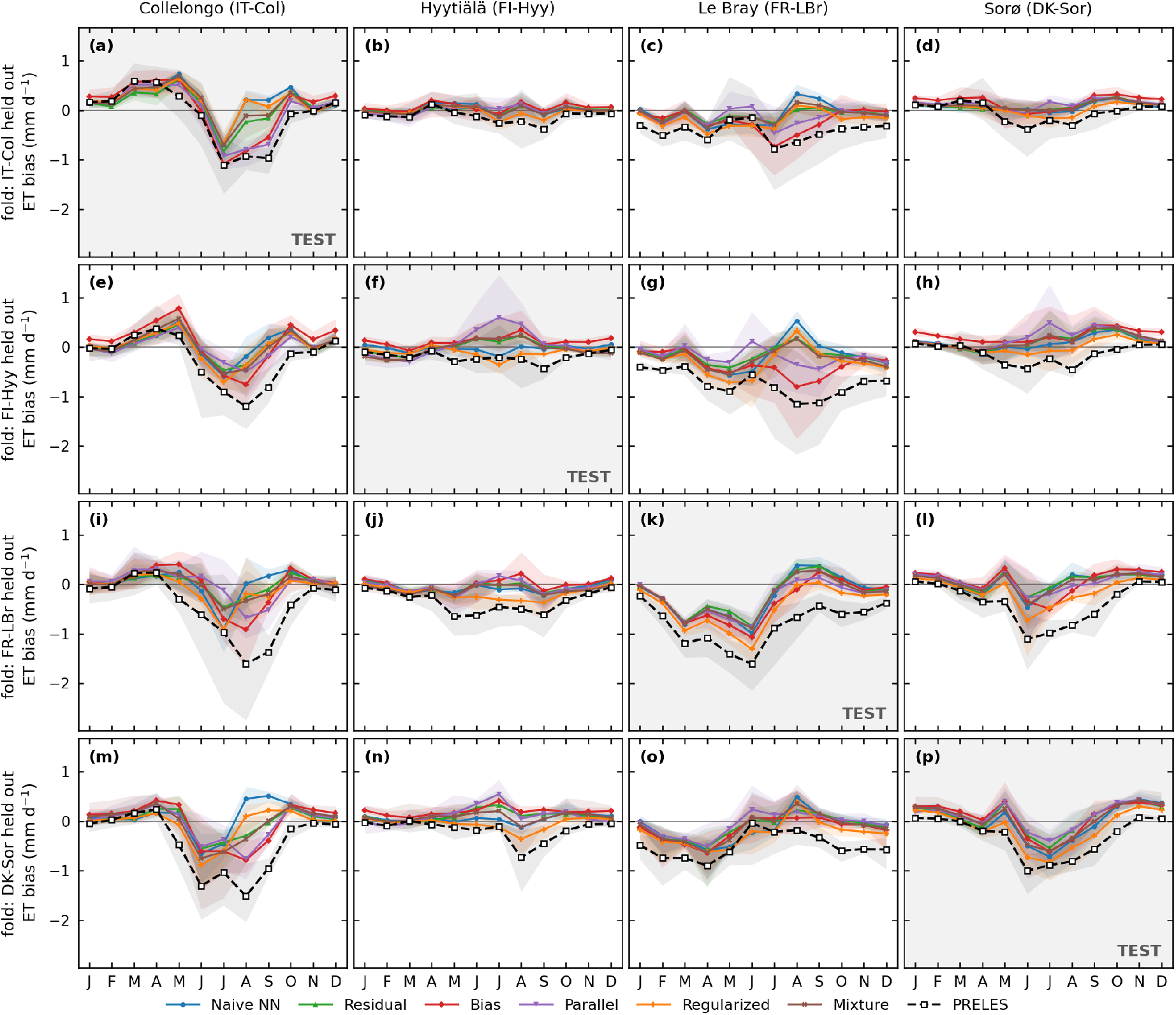
Monthly bias of daily **ET** for all four held-out stations of the spatial NCV across full training lengths shown for training and test (rows: Collelongo 2011–2013, Hyytiälä 2010–2012, Le Bray 2006–2008, Soro 2006–2008). Bias = mean(pred - obs) per calendar month, pooled over each station’s three test years; positive values indicate over-prediction. For each NN variant, the line is the mean over five HPO configurations and the band spans min–max across configurations; PRELES (dashed) is a single calibrated run. Note the differing y-axis ranges between rows.

**Table A.11:** Spatial OOD experiments: mean test **NSE** (Nash–Sutcliffe efficiency) across folds per prediction strategy and held-out station, for GPP, ET and their average (GPP+ET). NSE performance is classified as *>* 0.7: excellent; 0.5-0.7: good; *<* 0.5: poor. The five HPO configurations are averaged within each training-length regime; cells report mean std over those regimes (the All-stations column pools over stations and regimes). GPP+ET is the mean of the GPP and ET NSE within each regime. Strategies are ranked by the overall combined (GPP+ET, all-stations) mean NSE; the highest mean per column is in bold. Average: mean ± std over the seven strategies.

| Rank | Strategy | Collelongo (Italy) |  |  | Hyytiälä (Finland) |  |  | Le Bray (France) |  |  | Sorø (Denmark) |  |  | All stations |  |  |
|---|---|---|---|---|---|---|---|---|---|---|---|---|---|---|---|---|
|  |  | GPP | ET | GPP+ET | GPP | ET | GPP+ET | GPP | ET | GPP+ET | GPP | ET | GPP+ET | GPP | ET | GPP+ET |
| 1 | bias | **0.61 ± 0.08** | 0.69 ± 0.04 | **0.65 ± 0.05** | **0.91 ± 0.02** | **0.86 ± 0.03** | **0.89 ± 0.02** | 0.44 ± 0.13 | 0.65 ± 0.04 | **0.55 ± 0.06** | 0.57 ± 0.08 | 0.79 ± 0.02 | 0.68 ± 0.04 | 0.63 ± 0.19 | 0.75 ± 0.09 | **0.69 ± 0.13** |
| 2 | parallel | 0.51 ± 0.06 | 0.66 ± 0.08 | 0.58 ± 0.03 | 0.82 ± 0.04 | 0.75 ± 0.12 | 0.79 ± 0.07 | 0.35 ± 0.12 | 0.65 ± 0.09 | 0.50 ± 0.07 | **0.60 ± 0.09** | **0.80 ± 0.03** | **0.70 ± 0.05** | 0.57 ± 0.19 | 0.71 ± 0.10 | 0.64 ± 0.12 |
| 3 | mixture | 0.45 ± 0.19 | 0.72 ± 0.08 | 0.58 ± 0.12 | 0.85 ± 0.03 | 0.85 ± 0.03 | 0.85 ± 0.02 | 0.31 ± 0.11 | 0.64 ± 0.08 | 0.47 ± 0.06 | 0.53 ± 0.12 | 0.77 ± 0.06 | 0.65 ± 0.09 | 0.54 ± 0.24 | 0.74 ± 0.10 | 0.64 ± 0.16 |
| 4 | residual | 0.44 ± 0.13 | **0.72 ± 0.06** | 0.58 ± 0.07 | 0.84 ± 0.03 | 0.83 ± 0.05 | 0.84 ± 0.03 | 0.23 ± 0.14 | **0.66 ± 0.04** | 0.44 ± 0.08 | 0.56 ± 0.09 | 0.79 ± 0.03 | 0.67 ± 0.06 | 0.52 ± 0.25 | **0.75 ± 0.08** | 0.63 ± 0.16 |
| 5 | process | 0.58 ± 0.05 | 0.58 ± 0.13 | 0.58 ± 0.06 | 0.85 ± 0.02 | 0.82 ± 0.04 | 0.83 ± 0.03 | **0.61 ± 0.08** | 0.31 ± 0.15 | 0.46 ± 0.11 | 0.51 ± 0.06 | 0.74 ± 0.06 | 0.62 ± 0.06 | **0.64 ± 0.14** | 0.61 ± 0.22 | 0.62 ± 0.15 |
| 6 | regularized | 0.42 ± 0.15 | 0.70 ± 0.09 | 0.56 ± 0.10 | 0.80 ± 0.04 | 0.84 ± 0.02 | 0.82 ± 0.03 | 0.51 ± 0.09 | 0.51 ± 0.09 | 0.51 ± 0.05 | 0.44 ± 0.13 | 0.72 ± 0.10 | 0.58 ± 0.12 | 0.54 ± 0.19 | 0.69 ± 0.14 | 0.62 ± 0.15 |
| 7 | naive | 0.37 ± 0.25 | 0.68 ± 0.15 | 0.52 ± 0.18 | 0.81 ± 0.04 | 0.85 ± 0.03 | 0.83 ± 0.03 | 0.18 ± 0.08 | 0.59 ± 0.07 | 0.38 ± 0.06 | 0.46 ± 0.14 | 0.73 ± 0.08 | 0.60 ± 0.11 | 0.45 ± 0.27 | 0.71 ± 0.13 | 0.58 ± 0.19 |
|  | Average | 0.48 ± 0.09 | 0.68 ± 0.05 | 0.58 ± 0.04 | 0.84 ± 0.04 | 0.83 ± 0.04 | 0.84 ± 0.03 | 0.38 ± 0.16 | 0.57 ± 0.13 | 0.47 ± 0.05 | 0.52 ± 0.06 | 0.76 ± 0.03 | 0.64 ± 0.04 | 0.56 ± 0.06 | 0.71 ± 0.05 | 0.63 ± 0.03 |

**Table A.12:**
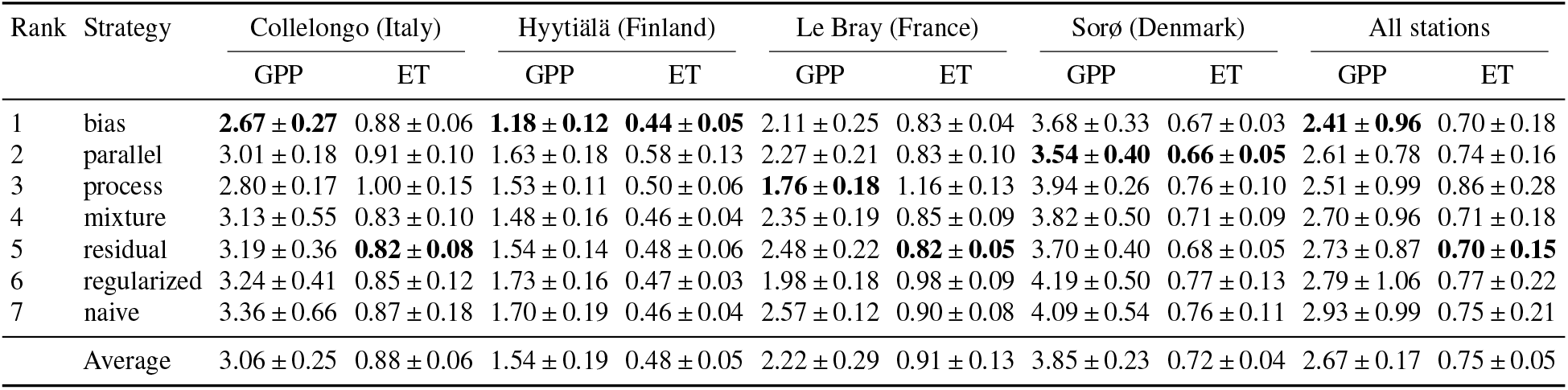
Spatial OOD experiments: mean across folds test **RMSE** per prediction strategy for GPP (g C m^−2^ d^−1^) and ET (mm d^−1^). The five HPO configurations are averaged within each training-length regime; cells report mean std over those regimes (the All-stations column pools over stations and regimes). Strategies are ranked by the overall combined (GPP+ET, all-stations) mean RMSE; lowest mean per column is in bold.

| Rank | Strategy | Collelongo (Italy) |  | Hyytiälä (Finland) |  | Le Bray (France) |  | Sorø (Denmark) |  | All stations |  |
|---|---|---|---|---|---|---|---|---|---|---|---|
|  |  | GPP | ET | GPP | ET | GPP | ET | GPP | ET | GPP | ET |
| 1 | bias | **2.67 ± 0.27** | 0.88 ± 0.06 | **1.18 ± 0.12** | **0.44 ± 0.05** | 2.11 ± 0.25 | 0.83 ± 0.04 | 3.68 ± 0.33 | 0.67 ± 0.03 | **2.41 ± 0.96** | 0.70 ± 0.18 |
| 2 | parallel | 3.01 ± 0.18 | 0.91 ± 0.10 | 1.63 ± 0.18 | 0.58 ± 0.13 | 2.27 ± 0.21 | 0.83 ± 0.10 | **3.54 ± 0.40** | **0.66 ± 0.05** | 2.61 ± 0.78 | 0.74 ± 0.16 |
| 3 | process | 2.80 ± 0.17 | 1.00 ± 0.15 | 1.53 ± 0.11 | 0.50 ± 0.06 | **1.76 ± 0.18** | 1.16 ± 0.13 | 3.94 ± 0.26 | 0.76 ± 0.10 | 2.51 ± 0.99 | 0.86 ± 0.28 |
| 4 | mixture | 3.13 ± 0.55 | 0.83 ± 0.10 | 1.48 ± 0.16 | 0.46 ± 0.04 | 2.35 ± 0.19 | 0.85 ± 0.09 | 3.82 ± 0.50 | 0.71 ± 0.09 | 2.70 ± 0.96 | 0.71 ± 0.18 |
| 5 | residual | 3.19 ± 0.36 | **0.82 ± 0.08** | 1.54 ± 0.14 | 0.48 ± 0.06 | 2.48 ± 0.22 | **0.82 ± 0.05** | 3.70 ± 0.40 | 0.68 ± 0.05 | 2.73 ± 0.87 | **0.70 ± 0.15** |
| 6 | regularized | 3.24 ± 0.41 | 0.85 ± 0.12 | 1.73 ± 0.16 | 0.47 ± 0.03 | 1.98 ± 0.18 | 0.98 ± 0.09 | 4.19 ± 0.50 | 0.77 ± 0.13 | 2.79 ± 1.06 | 0.77 ± 0.22 |
| 7 | naive | 3.36 ± 0.66 | 0.87 ± 0.18 | 1.70 ± 0.19 | 0.46 ± 0.04 | 2.57 ± 0.12 | 0.90 ± 0.08 | 4.09 ± 0.54 | 0.76 ± 0.11 | 2.93 ± 0.99 | 0.75 ± 0.21 |
|  | Average | 3.06 ± 0.25 | 0.88 ± 0.06 | 1.54 ± 0.19 | 0.48 ± 0.05 | 2.22 ± 0.29 | 0.91 ± 0.13 | 3.85 ± 0.23 | 0.72 ± 0.04 | 2.67 ± 0.17 | 0.75 ± 0.05 |

**Table A.13:**
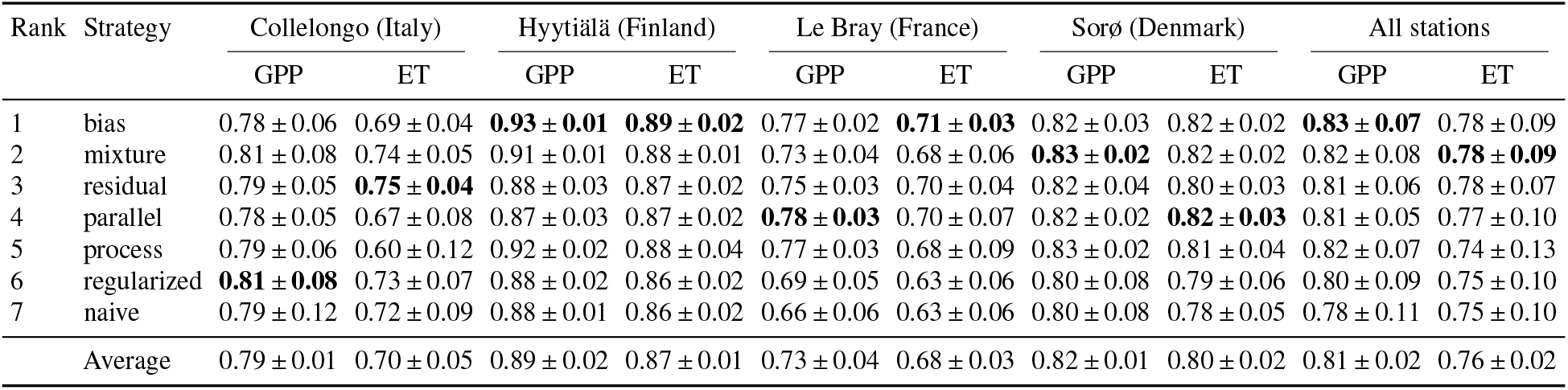
Spatial OOD experiments: mean across folds test *r*^2^ (squared Pearson correlation between observed and predicted) per prediction strategy and held-out station, for GPP and ET. *r*^2^ captures pattern/phase agreement only and ignores bias and scale errors (cf. NSE). The five HPO configurations are averaged within each training-length regime; cells report mean std over those regimes (the All-stations column pools over stations and regimes). Strategies are ranked by the overall combined (GPP+ET, all-stations) mean *r*^2^; highest mean per column is in bold.

**Table A.14:** Loss of performance when transferring to an unseen station (OOD experiments): increase of the error 1 − NSE from the training data to the held-out station, split into its additive components 1 − NSE = level + amplitude + timing, with 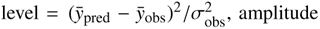 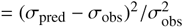 and 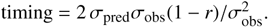 Values are test minus training, averaged over the seven training-set sizes and the five HPO configurations, with all four held-out stations weighted equally; in parentheses the share of each component in the total increase. The smallest increase per column is in bold. PGNN mean: average of the physics-guided strategies (regularized, residual, bias, parallel, mixture); PGNN process: additional loss of the PGNNs compared with the process model, in parentheses the relative difference (PGNN process)*/*process. Training values are normalised by the variance of the pooled training stations, test values by that of the held-out station.

| Strategy | Level | Amplitude | Timing | Total |
|---|---|---|---|---|
| *GPP* |  |  |  |  |
| process | **0.075** (54%) | **0.058** (42%) | **0.006** (4%) | **0.138** |
| naive | 0.188 (47%) | 0.099 (25%) | 0.113 (28%) | 0.400 |
| regularized | 0.146 (47%) | 0.079 (25%) | 0.088 (28%) | 0.313 |
| residual | 0.159 (43%) | 0.097 (27%) | 0.110 (30%) | 0.366 |
| bias | 0.094 (49%) | 0.072 (37%) | 0.027 (14%) | 0.193 |
| parallel | 0.116 (40%) | 0.092 (32%) | 0.083 (28%) | 0.290 |
| mixture | 0.156 (50%) | 0.089 (28%) | 0.068 (22%) | 0.314 |
| PGNN mean | 0.134 (45%) | 0.086 (29%) | 0.075 (25%) | 0.295 |
| PGNN − process | +0.060 (+80%) | +0.028 (+49%) | +0.069 (+1133%) | +0.157 (+113%) |
| *ET* |  |  |  |  |
| process | 0.045 (64%) | **0.007** (10%) | **0.018** (26%) | 0.071 |
| naive | 0.020 (15%) | 0.022 (16%) | 0.093 (69%) | 0.135 |
| regularized | 0.036 (24%) | 0.022 (15%) | 0.093 (61%) | 0.151 |
| residual | **0.013** (11%) | 0.017 (15%) | 0.089 (75%) | 0.119 |
| bias | 0.019 (34%) | 0.015 (27%) | 0.022 (39%) | **0.056** |
| parallel | 0.018 (14%) | 0.034 (27%) | 0.073 (59%) | 0.125 |
| mixture | 0.016 (16%) | 0.022 (21%) | 0.064 (63%) | 0.101 |
| PGNN mean | 0.020 (18%) | 0.022 (20%) | 0.068 (62%) | 0.110 |
| PGNN − process | −0.025 (−55%) | +0.015 (+212%) | +0.050 (+272%) | +0.040 (+56%) |

**Figure A.13:**
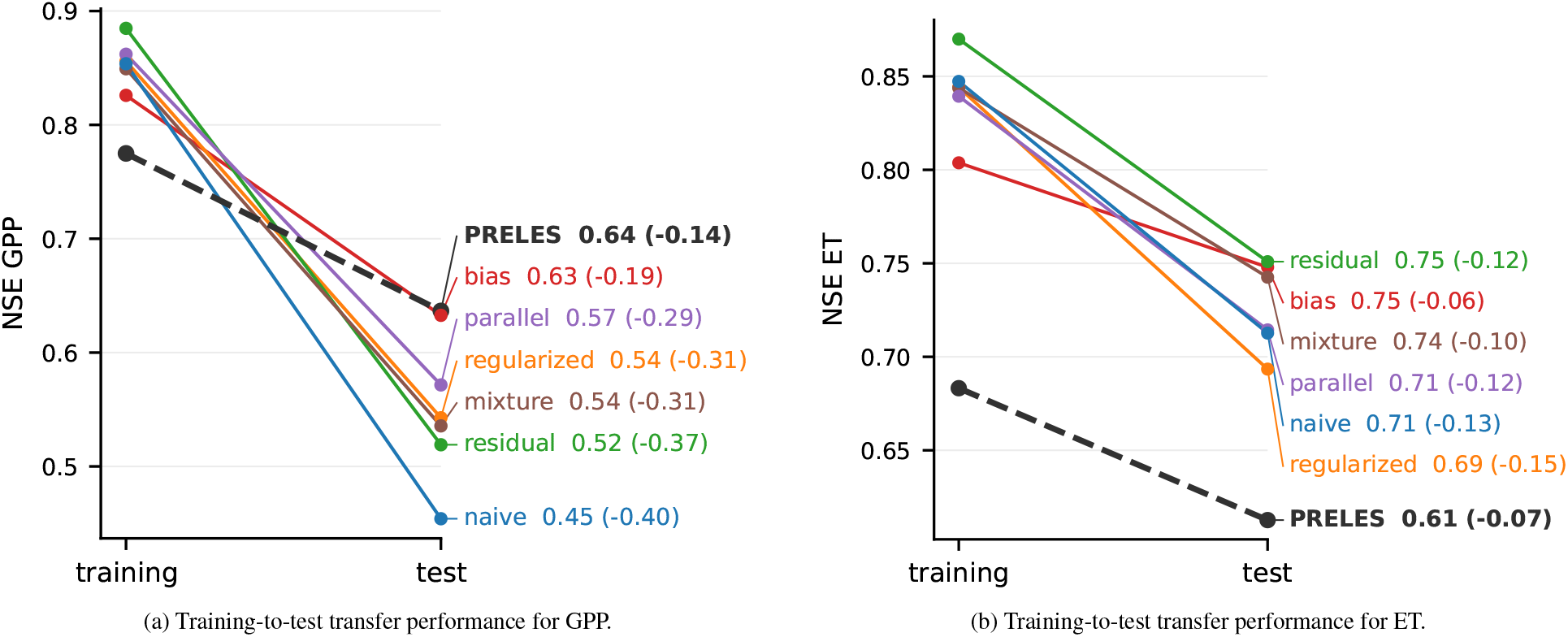
Comparison of training and test NSE across all OOD experiments. The value following each model name indicates the absolute test NSE, while the value in parentheses represents the difference between test and training NSE (NSE_test_ − NSE_train_).

## Appendix B. Post-hoc Analyses

### Appendix B.1. 1D ALE response curves

The process model (PRELES) and the network models all produced similar response curves for most inputs (Figures B.14 and B.15). GPP and ET increased with PAR in every fold. A slight divergence was observed at high light levels, where several networks flattened while the process model (PRELES) stayed close to linear. Both fluxes also increased with fAPAR, except in one case. ET in the Le Bray fold was almost insensitive to fAPAR.

The models disagreed most over the VPD response. For GPP, the models disagreed on the strength of the response, not its direction. Predicted GPP fell with VPD in every model, and the reductions agreed closely in the Collelongo and Hyytiälä folds. For the Sorø and Le Bray folds, the process model (PRELES) responded more weakly than the networks. For ET, the models also disagreed on direction (see main section 4.0.1). The process model (PRELES) ET fell with VPD in the Collelongo and Hyytiälä folds but rose in the Le Bray and Sorø folds. The *naive* model’s ET fell in all four. The hybrids generally lay between the two. The joint VPD PAR surfaces show the same pattern (Figure B.16). ET peaked at high PAR and moderate VPD in most panels. In the process model (PRELES) for the Le Bray and Sorø folds, and in parallel in the Le Bray fold, ET instead stayed high or kept rising towards the highest VPD range.

We interpret the shared PAR and fAPAR responses, and the negative effect of VPD on GPP, as common light and stomatal control that is physically plausible. The ET divergence follows from the process model (PRELES) calibration. Observed warm-season ET, by contrast, fell with VPD at every training site after controlling for Tair and PAR. This supports the sign of the networks’ ET response.

Air temperature and precipitation account for the remaining differences. The process model (PRELES) responded little to same-day Tair. The naive model, in contrast, increased sigmoidally around 10^◦^C, and the parallel and residual models increased less steeply. Same-day precipitation had no effect on process model (PRELES)’s GPP and almost none on ET. The networks responded weakly, with wide bands across draws at high rainfall. The Tair and precipitation differences should not be read as physical disagreement. The process model (PRELES) passes these inputs mainly through state variables (temperature acclimation, soil water, snow). The ALE holds those states at their unperturbed start-of-day values, so the process model (PRELES) curves understate the total effect of Tair and precipitation. The network components carry no such states and map these inputs directly onto the fluxes, so their same-day ALE captures effects that the process model (PRELES) cannot express within a single day.

### Appendix B.2. 2D ALE Response heatmap

**Figure B.14:**
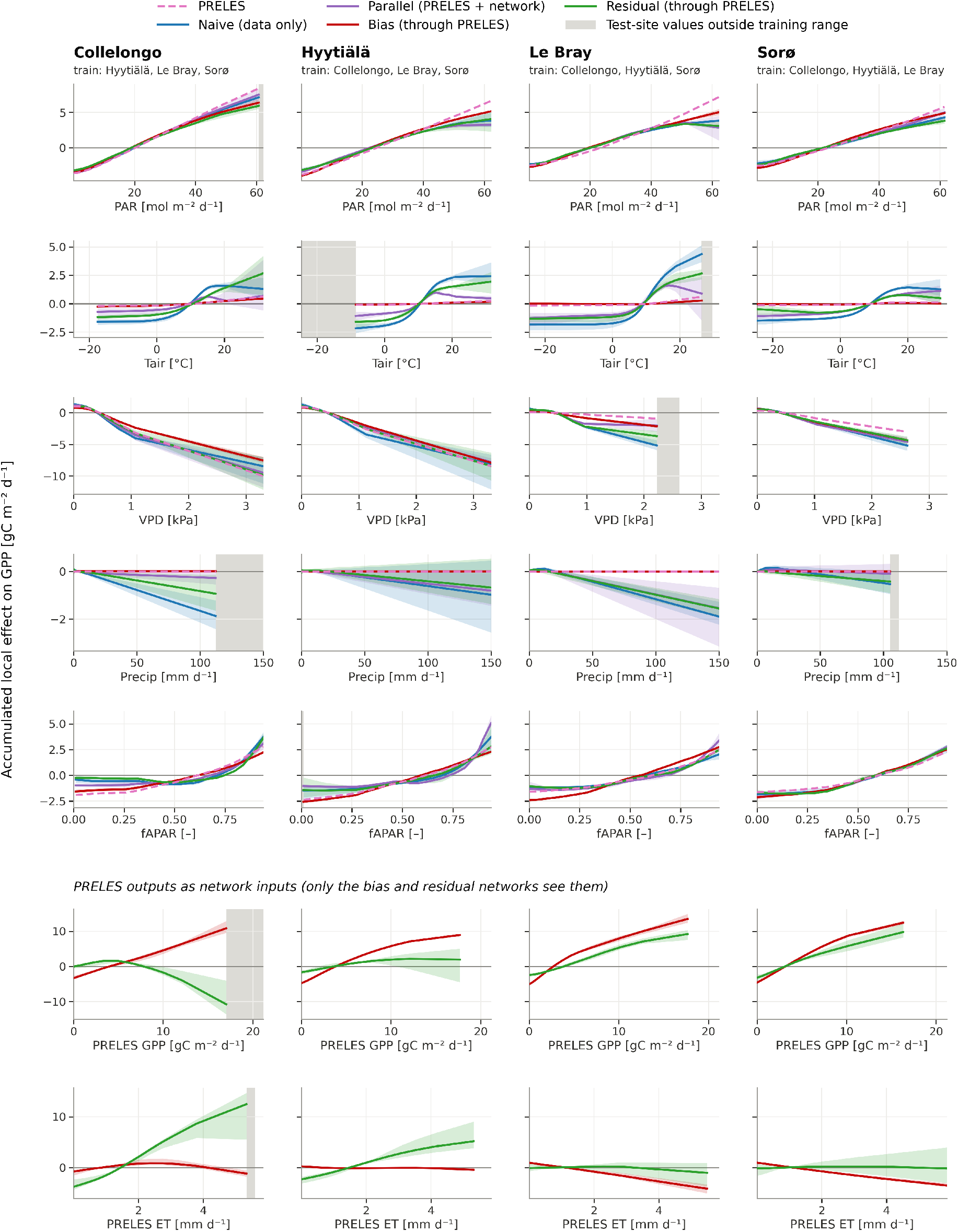
Model response (ALE) in predicting daily GPP. based on each input variable across the four held-out stations of the out-of-distribution experiments (columns: test site Collelongo, Hyytiala, Le Bray, Soro; each fold is trained on the other three sites). Curves are accumulated local effects (ALE) computed on the fold’s training days, in physical units (gC m^−2^ d^−1^). ALE curves are centred, showing how the prediction changes with the input. PRELES (dashed) is the process model’s (PRELES) response. Parallel = PRELES ALE plus the network’s ALE. Bias and residual: the perturbed input day is run through PRELES and then through the network, so their curves include the response passed on through PRELES. plain ANN (naive). Bottom two rows: response of the bias and residual networks to PRELES’ GPP and ET outputs, which only these two networks take as inputs (PRELES output perturbed, network re-run). Lines: mean over five configurations; bands: min–max across configurations. Grey shading: input values that occur on the test site’s days but lie outside the training range (extrapolation).

**Figure B.15:**
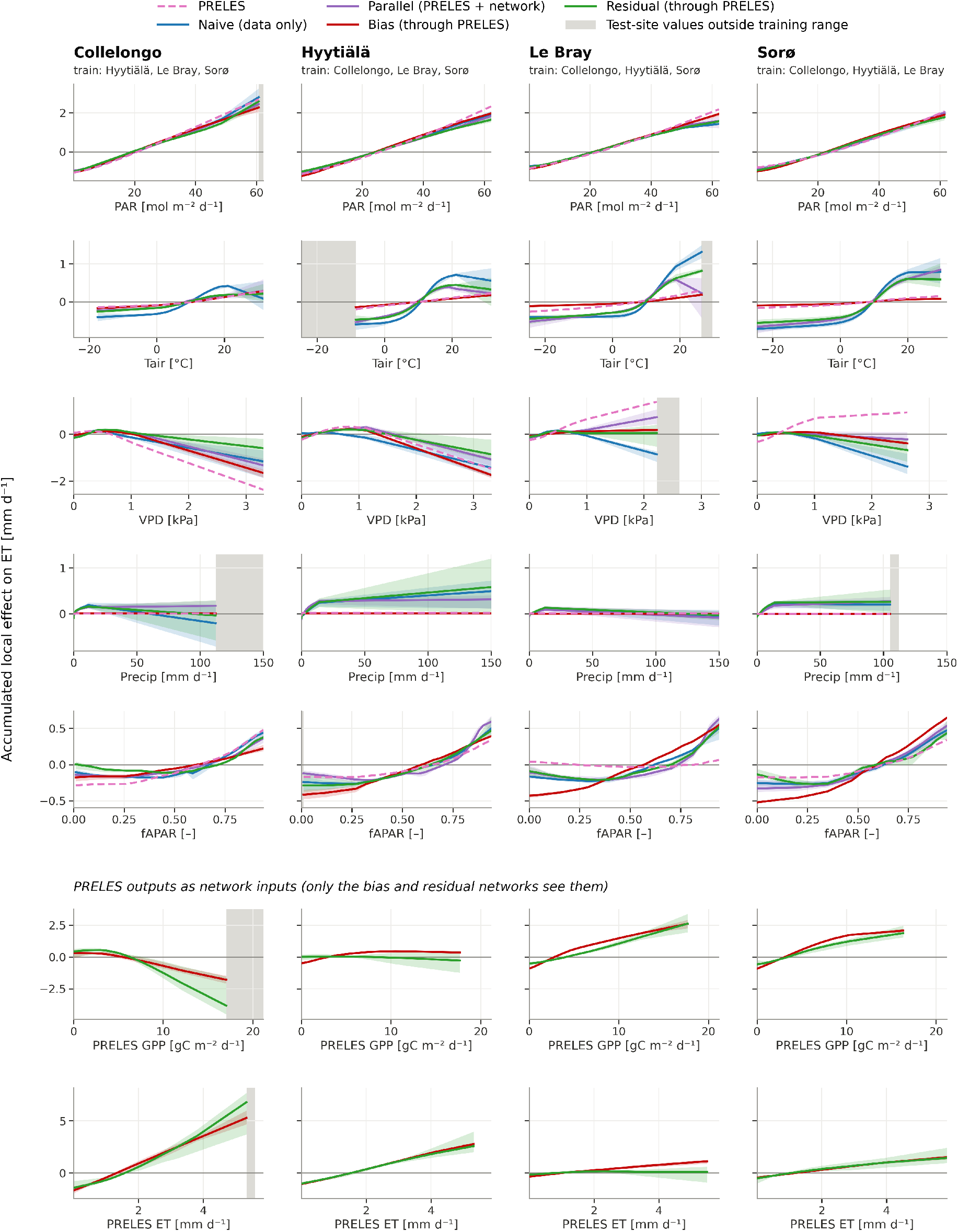
Model response (ALE) in predicting daily ET. based on each input variable across the four held-out stations of the out-of-distribution experiments (columns: test site Collelongo, Hyytiala, Le Bray, Soro; each fold is trained on the other three sites). Curves are accumulated local effects (ALE) computed on the fold’s training days, in physical units (mm d^−1^). ALE curves are centred, showing how the prediction changes with the input. PRELES (dashed) is the process model’s (PRELES) response. Parallel = PRELES ALE plus the network’s ALE. Bias and residual: the perturbed input day is run through PRELES and then through the network, so their curves include the response passed on through PRELES. Plain ANN (naive). Bottom two rows: response of the bias and residual networks to PRELES’ GPP and ET outputs, which only these two networks take as inputs (PRELES output perturbed, network re-run). Lines: mean over five configurations; bands: min–max across configurations. Grey shading: input values that occur on the test site’s days but lie outside the training range (extrapolation).

**Figure B.16:**
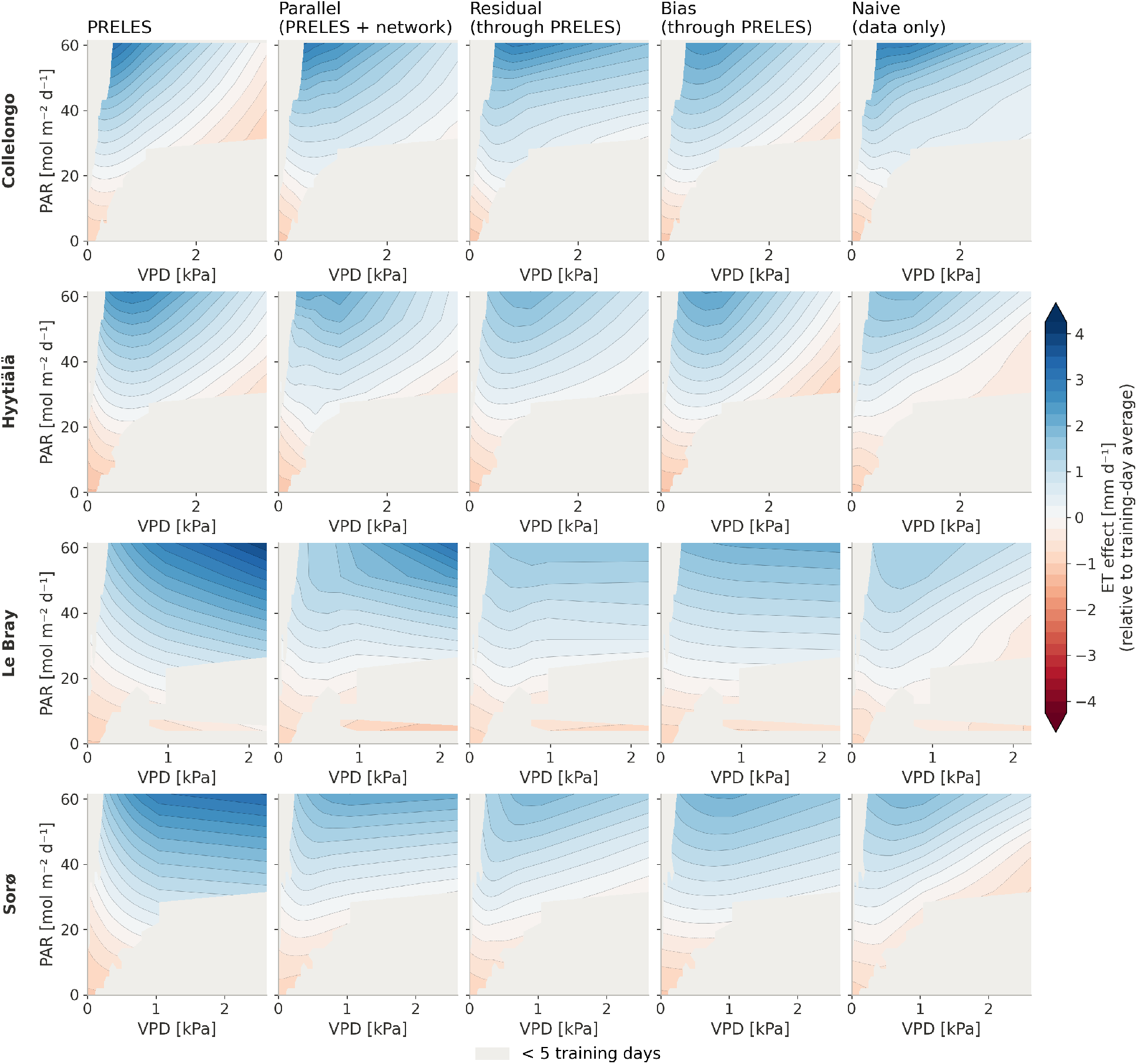
2nd-Order ALE of PAR-VPD on predicted daily ET. based on each input variable across the four held-out stations of the out-of-distribution experiments. Each surface is the sum of the main effects of PAR and VPD and their centred pairwise interaction, computed on the fold’s training days (all months) in physical units. Surfaces are centred: colours show ET above (blue) or below (red) the training-day average, not the ET level. The hybrid surfaces include only the same-day response through the process model (PRELES); effects carried over through the process model (PRELES)’ state variables (soil water, snow, acclimation) are not included. NN surfaces are the mean over five configurations; the process model (PRELES) is a single calibrated run.

**Table B.15:** ET skill at each site as a training site and as the held-out test site, across training regimes. The four training regimes differ in training length (1, 2, 3 and 4 years per training site, every day used); the test years are the same in every regime. <u>As training site</u>: computed over the site’s training days and averaged over the three folds in which the site is used for training. <u>As test site</u>: computed over the three test years. Daily *r*^2^: squared Pearson correlation between observed and predicted daily ET. <u>Level</u>: 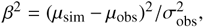 the NSE lost to the mean offset alone, from the decomposition 1 NSE = *β*^2^ + (*α* 1)^2^ + 2*α*(1 *r*) with *α* = *σ*_sim_*/σ*_obs_. Process model (PRELES): the fold’s calibrated model. Networks: per strategy the median over five HPO configurations, then the median over the six strategies (naive, regularised, residual, bias, parallel, mixture). Values are the mean ± standard deviation over the four training lengths (*n* = 4).

|  | As training site |  |  |  | As test site |  |  |  |
|---|---|---|---|---|---|---|---|---|
|  | Daily *r*<sup>2</sup> |  | Level |  | Daily *r*<sup>2</sup> |  | Level |  |
| Site | PRELES | Networks | PRELES | Networks | PRELES | Networks | PRELES | Networks |
| Collelongo | 0.74 ± 0.05 | 0.85 ± 0.02 | 0.05 ± 0.02 | 0.00 ± 0.00 | 0.61 ± 0.10 | 0.77 ± 0.01 | 0.01 ± 0.01 | 0.01 ± 0.01 |
| Hyytiälä | 0.86 ± 0.02 | 0.90 ± 0.00 | 0.04 ± 0.01 | 0.00 ± 0.00 | 0.87 ± 0.05 | 0.88 ± 0.01 | 0.04 ± 0.01 | 0.00 ± 0.00 |
| Le Bray | 0.66 ± 0.02 | 0.76 ± 0.03 | 0.20 ± 0.04 | 0.01 ± 0.00 | 0.70 ± 0.07 | 0.71 ± 0.04 | 0.31 ± 0.06 | 0.03 ± 0.01 |
| Sorø | 0.81 ± 0.04 | 0.86 ± 0.02 | 0.03 ± 0.01 | 0.01 ± 0.00 | 0.82 ± 0.03 | 0.83 ± 0.01 | 0.04 ± 0.04 | 0.01 ± 0.00 |

### Supplementary Materials

## Appendix C. Data preprocessing

### Appendix C.1. fAPAR sensitivity analysis

Implausible fAPAR values were identified using two complementary checks, one based on the BIDS (Viovy et al., 1992), which flags a value if it is below a fixed threshold of its immediate neighbours. The second is a robust outlier detection method (Pearson et al., 2016) where rolling windows compute the local median and the local MAD for each point. A composite is flagged if it falls below 3 times the MAD-based standard deviation of the local median. Based on comparisons with alternative settings, we selected a window step size of 7 and a threshold of 0.20. To characterise the effect of choosing a different combination of parameters, we compared the resulting cleaned series with alternative window sizes (3, 5, 9, 11) and fixed thresholds (0.15, 0.25 and 0.30) for each of the four sites.

We report how much the set of raw measurements (16-day resolution) classified as implausible changes (raw count), and how much this changes the daily fAPAR series actually used by the models (daily level), under these different parameter settings. Using our settings (window length = 7, threshold = 0.20) across all sites, an average of 29 measurements were flagged as implausible. Under less stringent alternatives, up to 8 would no longer be flagged; conversely, the stricter alternatives could flag up to 19 additional measurements not flagged under our settings. Since each raw measurement is held constant across all days until the next valid value, a single reclassification can propagate to many days downstream, meaning that the cleaned forward-filled daily fAPAR values under alternative settings could shift. From our settings, a mean of 536 of the 5403 days ^2^ per site (413–687 across the 4 sites) were affected. The mean absolute difference between the fAPAR values of the raw, uncleaned and cleaned series is 0.37 (0.28–0.41 across sites) on a daily level. In contrast, when using stricter parameters, an average of 658 (635-676 across the 4 sites) days were affected, with a mean absolute difference of 0.35 (0.25–0.41) fAPAR units. Using a more relaxed setting affected an average of 422 (302-493 across the 4 sites) days, with a mean absolute difference of 0.38 (0.29-0.42). Our chosen parameters therefore represent a conservative middle ground. We acknowledge this as a limitation of the reconstructed fAPAR series but do not explore it further, as our primary interest lies in evaluating and comparing the models themselves rather than characterising their sensitivity to the preprocessing choices of the input data.

### Appendix C.2. Results: Sensitivity to gap-filled GPP, LE, and VPD

To assess whether the reported model skill is inflated by structural agreement with the meteorology-driven gap-filling procedure (MDS) used to construct GPP, latent heat flux (LE), and VPD, we evaluated a small number of our models on exclusively measured data. For each of the four sites we identified the subset of days on which GPP, LE, and VPD were all fully measured rather than gap-filled. A day was classified as directly measured only if every half-hourly value contributing to its daily aggregate carried a measured (non-gap-filled) quality flag for all three variables. This criterion is particularly demanding for GPP, since it is derived by daytime partitioning of net ecosystem exchange and a single gap-filled half-hour anywhere in the diel cycle disqualifies that day’s value. Across each site’s available full multi-year record, only 8 (Collelongo), 24 (Hyytiälä), 20 (Le Bray), and 658 (Sorø) days met the joint criterion. Sorø’s measured subset is large and close to seasonally representative of the full record (winter 31%, spring 27%, summer 15%, autumn 27%), spanning all eleven years (2002–2012) allowing us to make a comparison of model performance between full gap-filled heavy and measured-only datasets.

The plain ANN (*naive*) and *residual* models evaluated here are the spatial_len3 out-of-distribution checkpoints: each trained on one year of data from each of the three sites not held out as the test site (three years of training data in total), and evaluated on the fourth, unseen site. Note that these checkpoints are distinct from the twelve-year regime used in the main ALE analysis (section 2.8). We re-evaluated these models against their originally reported test-set performance using RMSE, mean bias (observed predicted), and NSE, with 90% bootstrap confidence intervals (5,000 resamples). For each site, target, and model, days were resampled with replacement, and the 5th and 95th percentiles of the resulting metric distributions were used as the 90% confidence intervals.

Our model performance comparison showed no loss of skill on the measured-only subset at Sorø for either model (plain ANN *naive* and *residual*, see Table C.16). For ET, RMSE and NSE were statistically indistinguishable between the full test set and the measured-only subset. For GPP, both models performed better on the measured-only subset, with non-overlapping 90% CIs for RMSE. The mean bias was positive for both models and both targets, i.e. both models under-predicted, and this under-prediction was smaller on the measured subset for GPP. For ET, the bias did not differ between the two sets. The direction is the opposite of what would be expected if the models were reproducing the gap-filling procedure’s structure, under which closer agreement on gap-filled days, not measured ones, would be the pattern of concern.

At the three smaller sites, the measured-only subsets are too small and seasonally skewed to support meaningful comparisons with the full test sets. Collelongo has only 8 days, all outside summer; Hyytiälä has 24 days, half in winter and none in summer. At Le Bray, 13 of these days fall in 2002, a year in which the GPP record is zero on every day (a missing year encoded as zero rather than a measurement); these days were excluded from the GPP evaluation, leaving 7 days (two in summer) for GPP and 20 for ET. Thus, differences between the full and measured-only performance cannot be separated from seasonal effects or sampling variability. NSE is particularly unstable at these sample sizes because it is normalised by the subset’s observed variance, resulting in very wide bootstrap intervals (Table C.16). We therefore report RMSE and bias for completeness but do not interpret these results as evidence of model skill on directly observed data.

**Table C.16:** Sensitivity of the plain ANN (*naive*) and *residual* model performance to gap-filled GPP, ET, and VPD. RMSE, mean bias (observed predicted; positive = under-prediction), and NSE for the full test set (<u>all</u>), and the directly-measured subset (<u>meas.</u>) at each site, with 90% bootstrap confidence intervals (CI) (5,000 resamples). Brackets show 90% bootstrap CIs. Models evaluated here were originally trained with three years of data (one year per site) and evaluated under a fourth unseen site (see Table 2). Due to the insufficient amount of measured subsets (n = 7–24) at Collelongo (Colle.), Hyytiälä (Hyy), and Le Bray; neither the RMSE/mean bias point estimates nor their CIs should be interpreted as evidence of differences between the full and measured-only performance here. CI’s (measured) are omitted for these sites^†^. Only Sorø (n = 658) supports a meaningful comparison on any metric.

| Site | Model | Target | *n*<sub>all</sub>/*n*<sub>meas.</sub> | RMSE<sub>all.</sub>[CI] | RMSE<sub>meas.</sub>[CI] | Bias<sub>all</sub>[CI] | Bias<sub>meas.</sub>[CI] | NSE<sub>all</sub>[CI] | NSE<sub>meas.</sub>[CI] |
|---|---|---|---|---|---|---|---|---|---|
| Colle. | naive | GPP | 1096 / 8 | 4.09 [3.94, 4.25] | 4.36 [1.95, 6.59] | −2.90 [−3.05, −2.76] | −3.16 [−5.09, −1.65] | 0.10 [0.02, 0.17] | −1.69<sup>†</sup> |
|  | naive | ET | 1096 / 8 | 0.77 [0.74, 0.81] | 0.98 [0.68, 1.21] | −0.17 [−0.21, −0.13] | −0.73 [−1.10, −0.34] | 0.76 [0.74, 0.78] | −2.51<sup>†</sup> |
|  | residual | GPP | 1096 / 8 | 3.71 [3.56, 3.86] | 3.23 [0.84, 5.34] | −2.16 [−2.31, −2.01] | −1.45 [−3.37, −0.01] | 0.26 [0.19, 0.32] | −0.48<sup>†</sup> |
|  | residual | ET | 1096 / 8 | 0.80 [0.75, 0.84] | 0.66 [0.49, 0.80] | 0.03 [−0.01, 0.07] | −0.44 [−0.71, −0.16] | 0.74 [0.72, 0.76] | −0.57<sup>†</sup> |
| Hyy. | naive | GPP | 1096 / 24 | 2.95 [2.80, 3.11] | 1.60 [0.42, 2.47] | 1.64 [1.52, 1.76] | −0.34 [−0.90, 0.13] | 0.43 [0.38, 0.48] | −2.48<sup>†</sup> |
|  | naive | ET | 1096 / 24 | 0.55 [0.52, 0.58] | 0.26 [0.16, 0.34] | 0.25 [0.22, 0.27] | −0.10 [−0.18, −0.03] | 0.78 [0.77, 0.80] | 0.21<sup>†</sup> |
|  | residual | GPP | 1096 / 24 | 2.55 [2.41, 2.68] | 1.39 [0.49, 2.21] | 1.45 [1.35, 1.55] | 0.01 [−0.52, 0.40] | 0.57 [0.53, 0.61] | −1.62<sup>†</sup> |
|  | residual | ET | 1096 / 24 | 0.47 [0.45, 0.50] | 0.26 [0.18, 0.34] | 0.16 [0.14, 0.18] | −0.07 [−0.15, 0.01] | 0.84 [0.82, 0.85] | 0.20<sup>†</sup> |
| Le Bray | naive | GPP | 1096 / 7 | 2.50 [2.39, 2.60] | 2.71 [1.12, 3.98] | −1.07 [−1.19, −0.96] | −1.34 [−2.88, 0.07] | 0.23 [0.13, 0.31] | −0.46<sup>†</sup> |
|  | naive | ET | 1096 / 20 | 0.90 [0.87, 0.94] | 0.96 [0.75, 1.14] | 0.27 [0.23, 0.31] | −0.32 [−0.65, 0.02] | 0.59 [0.56, 0.61] | 0.40<sup>†</sup> |
|  | residual | GPP | 1096 / 7 | 2.42 [2.32, 2.50] | 1.31 [0.69, 1.85] | −1.38 [−1.48, −1.28] | −0.87 [−1.48, −0.28] | 0.28 [0.20, 0.35] | 0.66<sup>†</sup> |
|  | residual | ET | 1096 / 20 | 0.79 [0.75, 0.83] | 0.64 [0.42, 0.82] | 0.20 [0.17, 0.24] | −0.14 [−0.36, 0.10] | 0.69 [0.66, 0.71] | 0.74<sup>†</sup> |
| Sorø | naive | GPP | 1096 / 658 | 4.11 [3.93, 4.28] | 3.34 [3.04, 3.64] | 2.34 [2.17, 2.51] | 1.54 [1.28, 1.81] | 0.47 [0.44, 0.50] | 0.56 [0.52, 0.60] |
|  | naive | ET | 1096 / 658 | 0.71 [0.68, 0.75] | 0.65 [0.59, 0.71] | 0.12 [0.09, 0.16] | 0.09 [0.04, 0.15] | 0.77 [0.75, 0.79] | 0.74 [0.70, 0.77] |
|  | residual | GPP | 1096 / 658 | 3.84 [3.66, 4.01] | 3.07 [2.79, 3.36] | 2.22 [2.07, 2.38] | 1.49 [1.26, 1.73] | 0.53 [0.51, 0.56] | 0.63 [0.59, 0.66] |
|  | residual | ET | 1096 / 658 | 0.65 [0.62, 0.69] | 0.59 [0.53, 0.64] | 0.06 [0.03, 0.09] | 0.06 [0.02, 0.11] | 0.81 [0.79, 0.82] | 0.79 [0.75, 0.82] |

<sup>†</sup> CI omitted due to sample size (n = 7–24). NSE itself is shown for transparency, but is not interpretable as a performance measure.

## Appendix D. Nested cross validation (NCV)

The following sections describe the NCV strategies in detail for (1) artificially thinned data and (2) in combination with OOD settings. To ensure reliable evaluation and eliminate biases from random data splits or favourable hyper-parameter configurations, we employ a nested cross-validation (Roberts et al., 2017) (Leave-One-Station-Out Cross-Validation (LOSO-NCV)) framework. Figure D.17 shows a systematic LOSO-NCV of this OOD experiment. This framework consists of an inner and outer loop: in the outer loop, we evaluate final model performance by testing on one completely unseen station; in the inner loop, we search for the optimal model architecture configuration using a subset of the training data from the outer loop split. This approach is applied consistently across all model types: the various PGNN variants (bias, parallel, regularized, residual and mixture), the stand-alone process model (PRELES), and a plain artificial neural network (ANN). For hyper-parameter tuning (details in Table A.4), we use Hyperband with successive halving (Li et al., 2018) to efficiently explore the hyper-parameter space.

1. Outer Loop (Station split): Each forest site (station) is left out in turn as the outer test set, while the remaining three stations are used for training. These three are then passed to the inner loop for model selection.
2. Inner Loop (Station split): Within each fold, the three training stations are further split: two are used for training, and one is used for validation and testing. This process is repeated so that each station serves as the validation/test set once.
3. Inner Loop (Hyper-parameter Optimisation): To evaluate various hyper-parameter configurations, each trial is tested across all inner folds: two stations are used for training, while the third is split into a test set for hyper-parameter tuning and a validation set for computing average validation scores. These scores guide the selection of the best configuration for final evaluation in the outer loop.
4. Outer loop (Final performance): Based on the average validation scores from the inner loop, the best model configuration are selected, retrained on the three training stations and evaluated on the hold-out station. Final performance is reported as the average across all hold-out stations.

To ensure fairness, a fixed number of years (1-4) was used per station, accounting for different data availabilities. We repeated this setup with varying lengths of training data. The final model performance was averaged across all test stations using metrics such as MSE, R², and RMSE.

### Domain-Shifts

To investigate the performance differences amongst our modelling strategies (pure ANN, PGNNs, process model), we quantified domain-shifts by observing differences in the correlation structures between variables from the training to the test sites. We then examined whether such shifts coincided with changes in model performance using variable importance analysis (ALE).

### Appendix D.0.1. Variable-association diagnostics

To quantify the changes in environmental conditions, we assessed domain-shifts by computing Pearson’s correlation for all pairs of input and output, and evaluated differences in the correlation of these pairs. This identified whether variable associations strengthened (positive correlation difference) or weakened (negative correlation difference) at the test site. All diagnostics were integrated into the evaluation pipeline and applied to each outer fold of the nested cross-validation procedure, ensuring consistent domain-shift assessment across experimental scenarios.

**Figure D.17:**
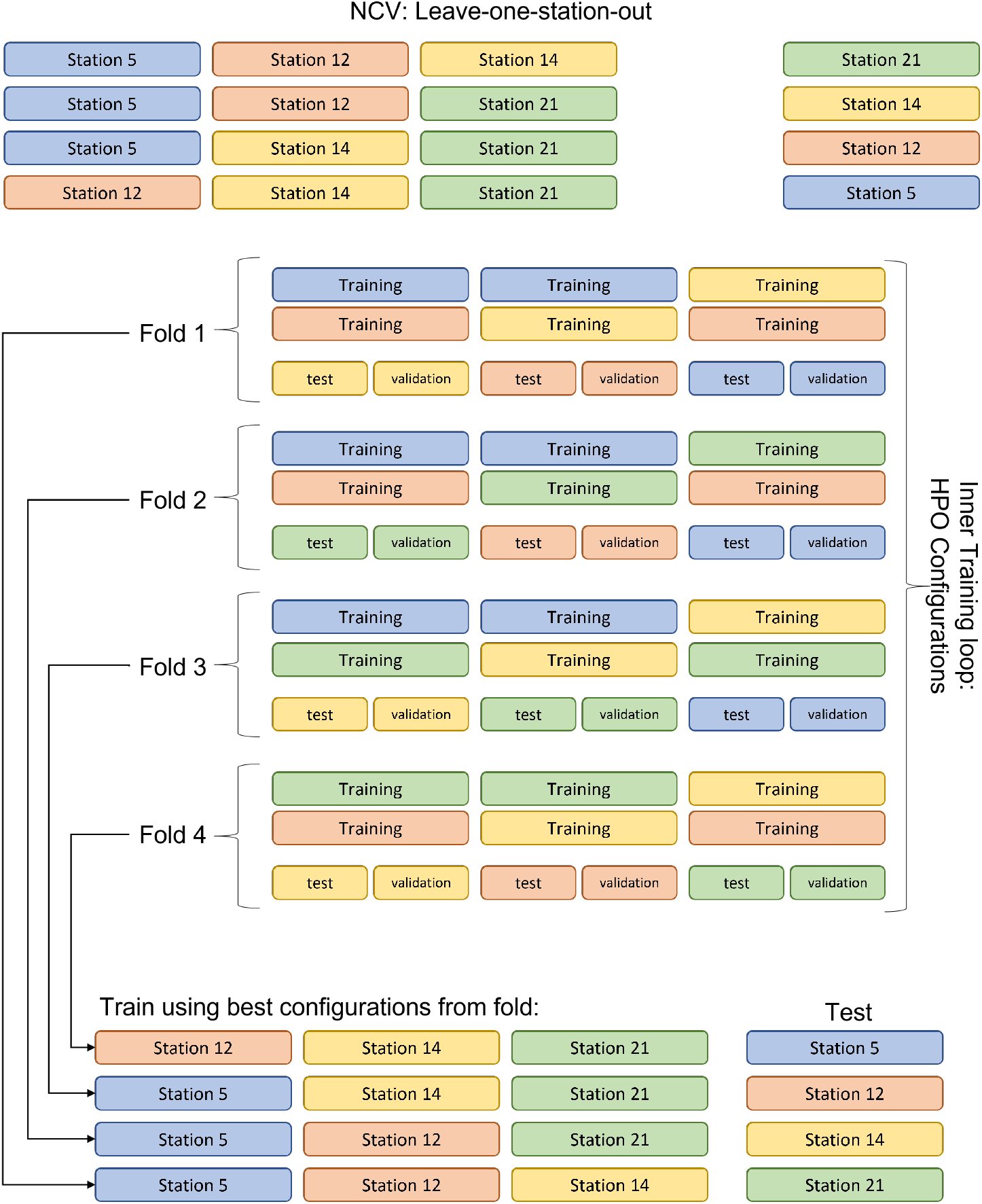
Schematic visualisation of spatial nested-cross validation across the four forest stations, Hyytiälä (12), Le Bray (14), Sorø (21), Collelongo (5)

**Figure D.18:**
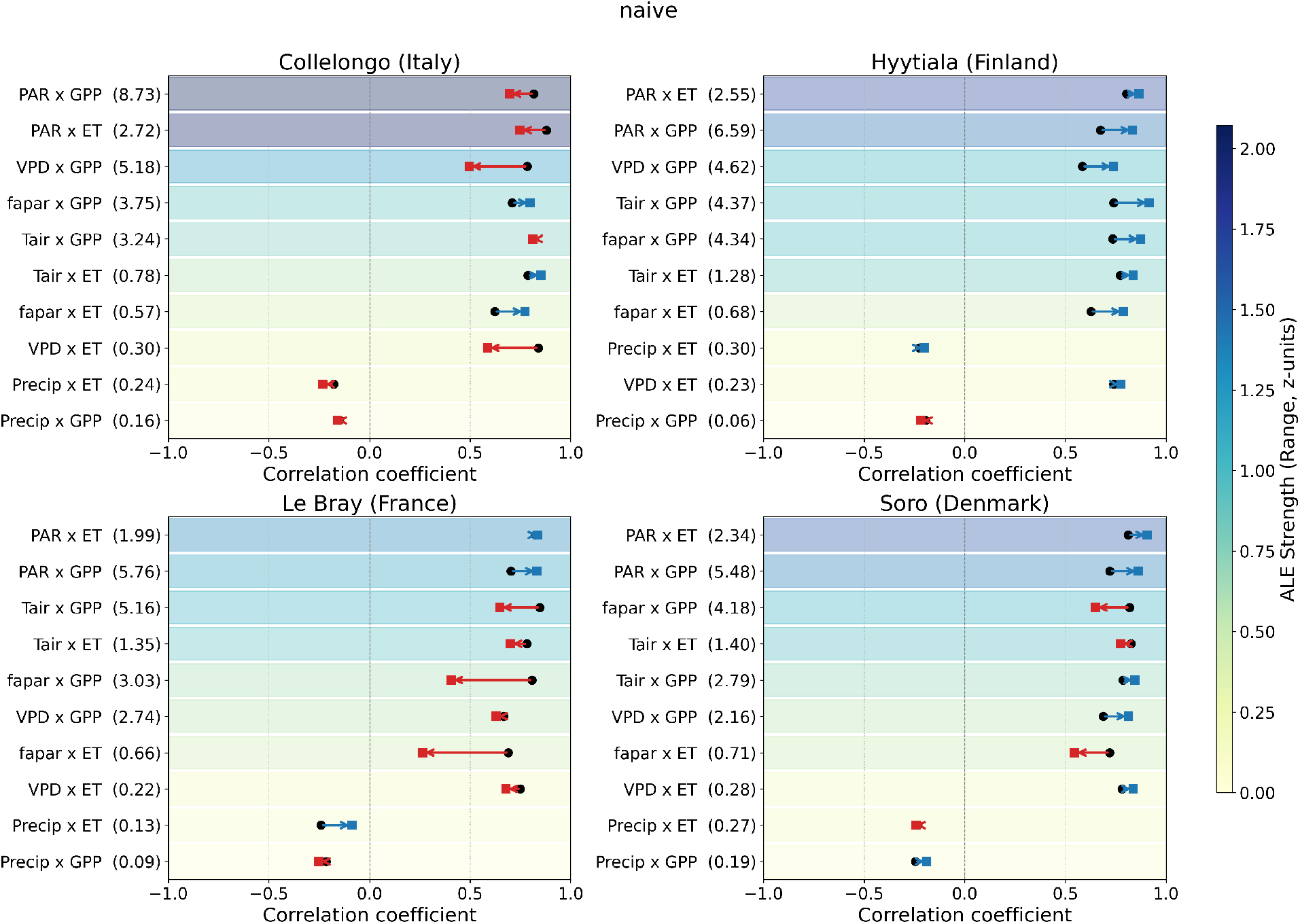
Domain-Shift ***Naive*** model: Predictor-target correlation shifts ranked by first-order ALE values (naive) of main effects on ET and GPP for each forest test station. Black circles represent the correlation coefficient in the training data, while the arrows show the correlation change at the test site, with red indicating a loss in correlation strength and blue an increase. The ALE range reflects the importance of each feature pair, and correlation pairs are ranked from strongest to weakest ALE amplitudes, with underlying heatmap values shown. Heatmap colours are drawn in z-scored units to keep GPP and ET comparable. The labels show the ALE-calculated influence in physical units per target.

**Figure D.19:**
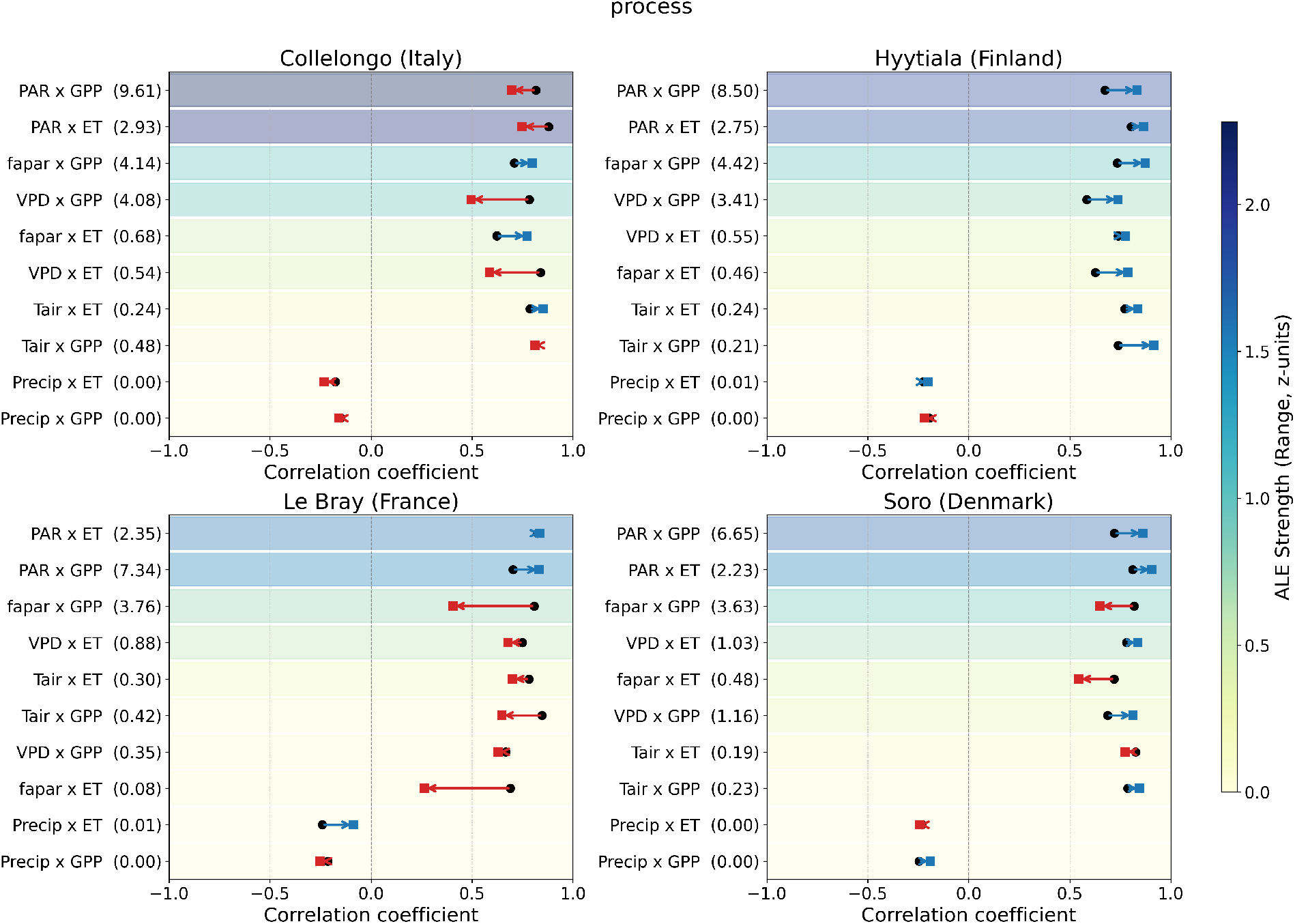
Domain-Shift ***process model(PRELES)***: Predictor-target correlation shifts ranked by first-order ALE values (process model) of main effects on ET and GPP for each forest test station. Black circles represent the correlation coefficient in the training data, while the arrows show the correlation change at the test site, with red indicating a loss in correlation strength and blue an increase. The ALE range reflects the importance of each feature pair, and correlation pairs are ranked from strongest to weakest ALE amplitudes, with underlying heatmap values shown. Heatmap colours are drawn in z-scored units to keep GPP and ET comparable. The labels show the ALE-calculated influence in physical units per target.

**Figure D.20:**
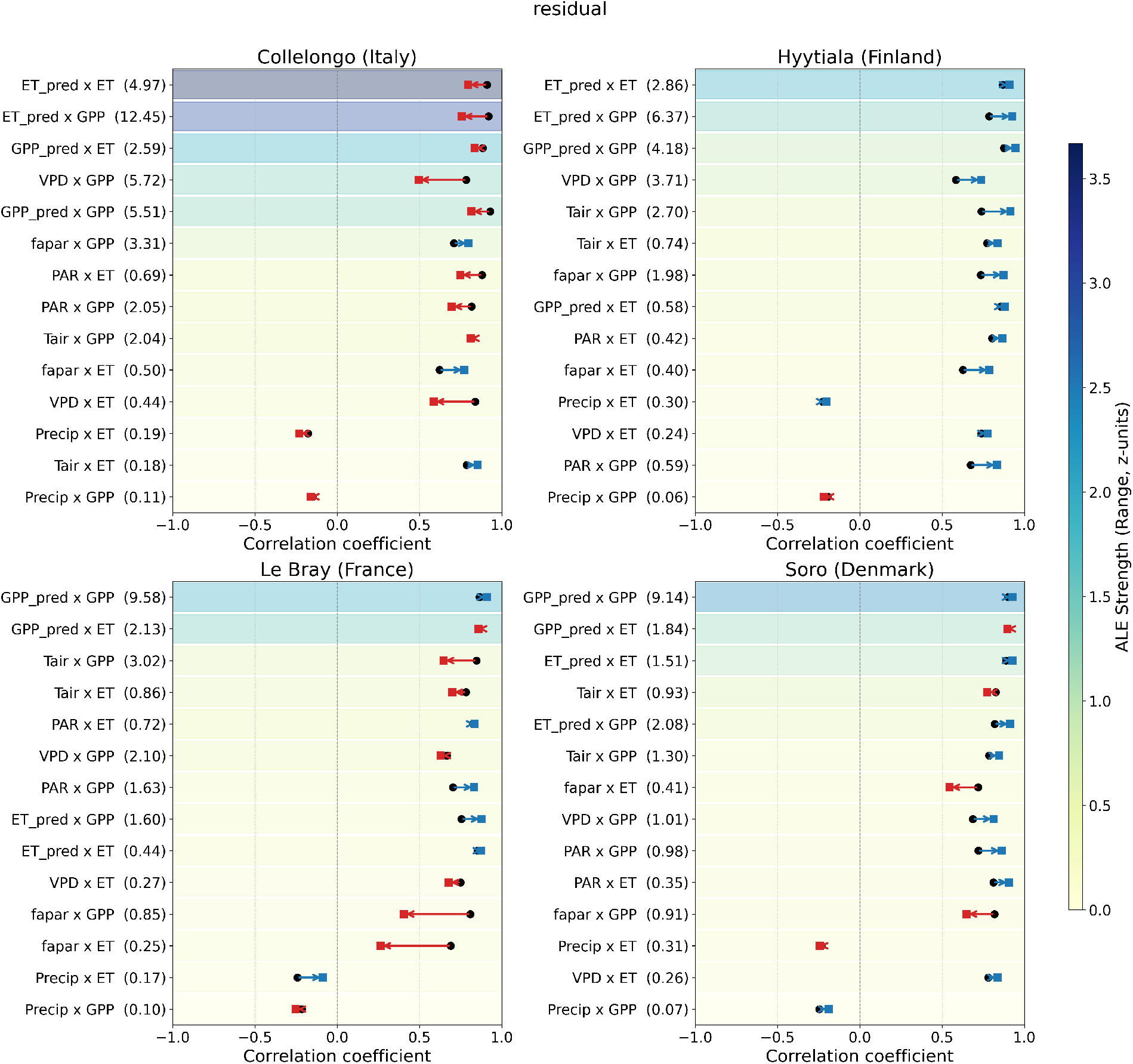
Domain-Shift ***Residual*** model: Predictor-target correlation shifts ranked by first-order ALE values (residual) of main effects on ET and GPP for each forest test station. Black circles represent the correlation coefficient in the training data, while the arrows show the correlation change at the test site, with red indicating a loss in correlation strength and blue an increase. The ALE range reflects the importance of each feature pair, and correlation pairs are ranked from strongest to weakest ALE amplitudes, with underlying heatmap values shown. Heatmap colours are drawn in z-scored units to keep GPP and ET comparable. The labels show the ALE-calculated influence in physical units per target.

**Figure D.21:**
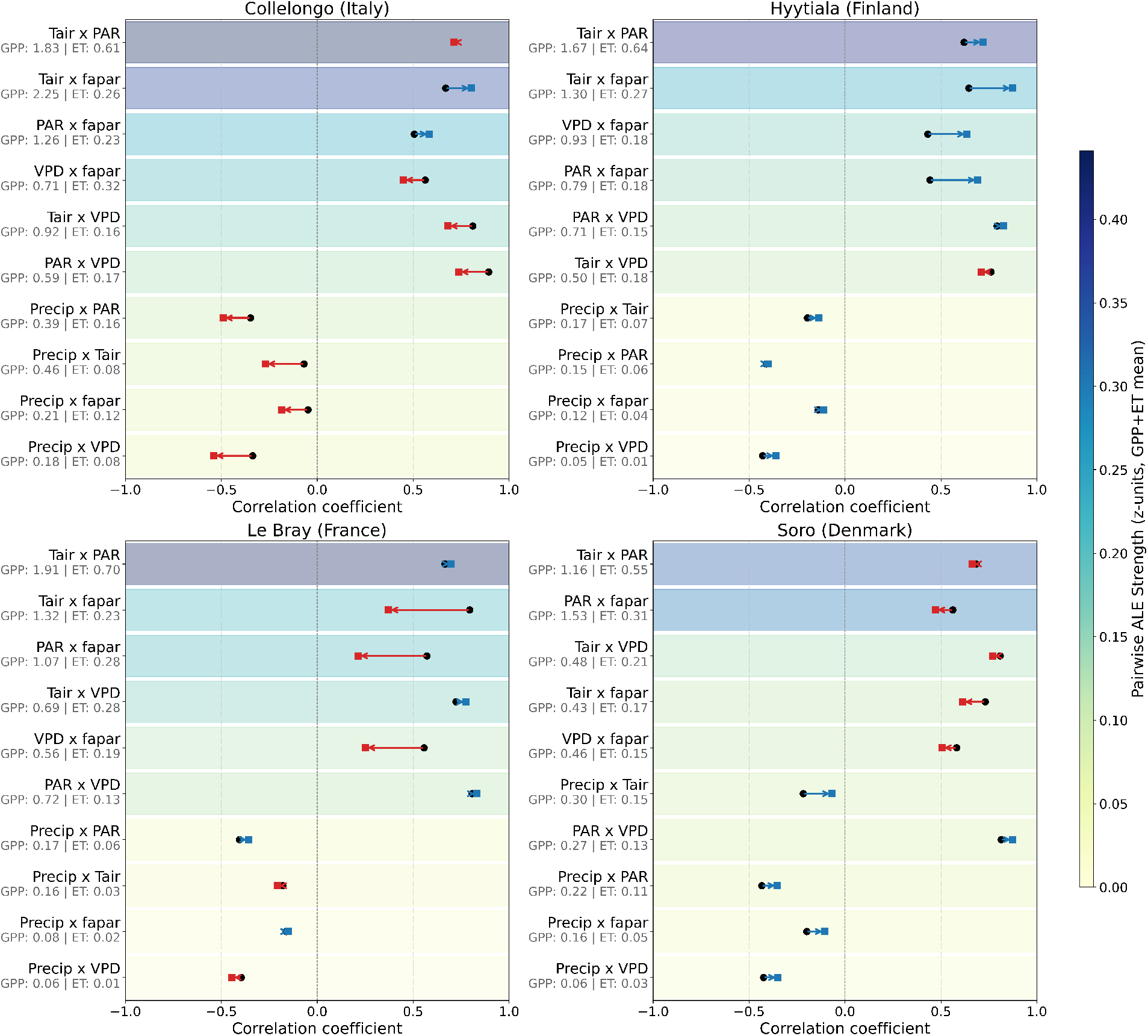
Domain-Shift ***Naive*** model: Predictor-predictor correlation shifts ranked by 2nd-Order ALE values showing the interaction effect on the targets (mean) per modelling strategy at Le Bray. Black circles represent the correlation coefficient within the training data, and the arrow indicates how this correlation changed at the test site. ALE range reflects the amplitude range of that feature pair. Correlation pairs filtered by strongest to weakest ALE amplitudes. Heatmap colours are drawn in z-scored units to keep GPP and ET comparable. The labels show the ALE-calculated influence in physical units per target.

**Figure D.22:**
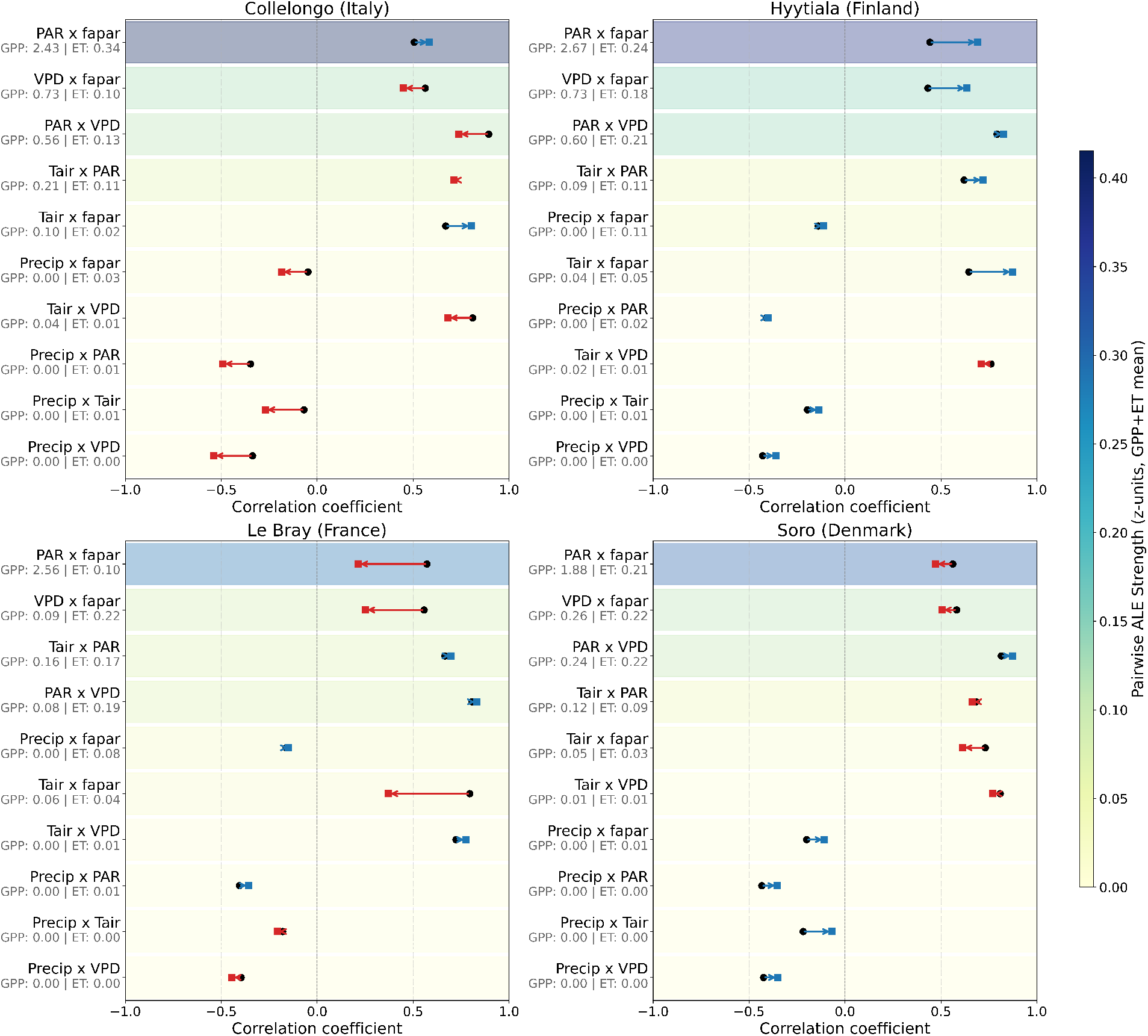
Domain-Shift ***Process*** model (PRELES): Predictor-predictor correlation shifts ranked by 2nd-Order ALE values showing the interaction effect on the targets (mean) per modelling strategy at Le Bray. Black circles represent the correlation coefficient within the training data, and the arrow indicates how this correlation changed at the test site. ALE range reflects the amplitude range of that feature pair. Correlation pairs filtered by strongest to weakest ALE amplitudes. Heatmap colours are drawn in z-scored units to keep GPP and ET comparable. The labels show the ALE-calculated influence in physical units per target.

**Figure D.23:**
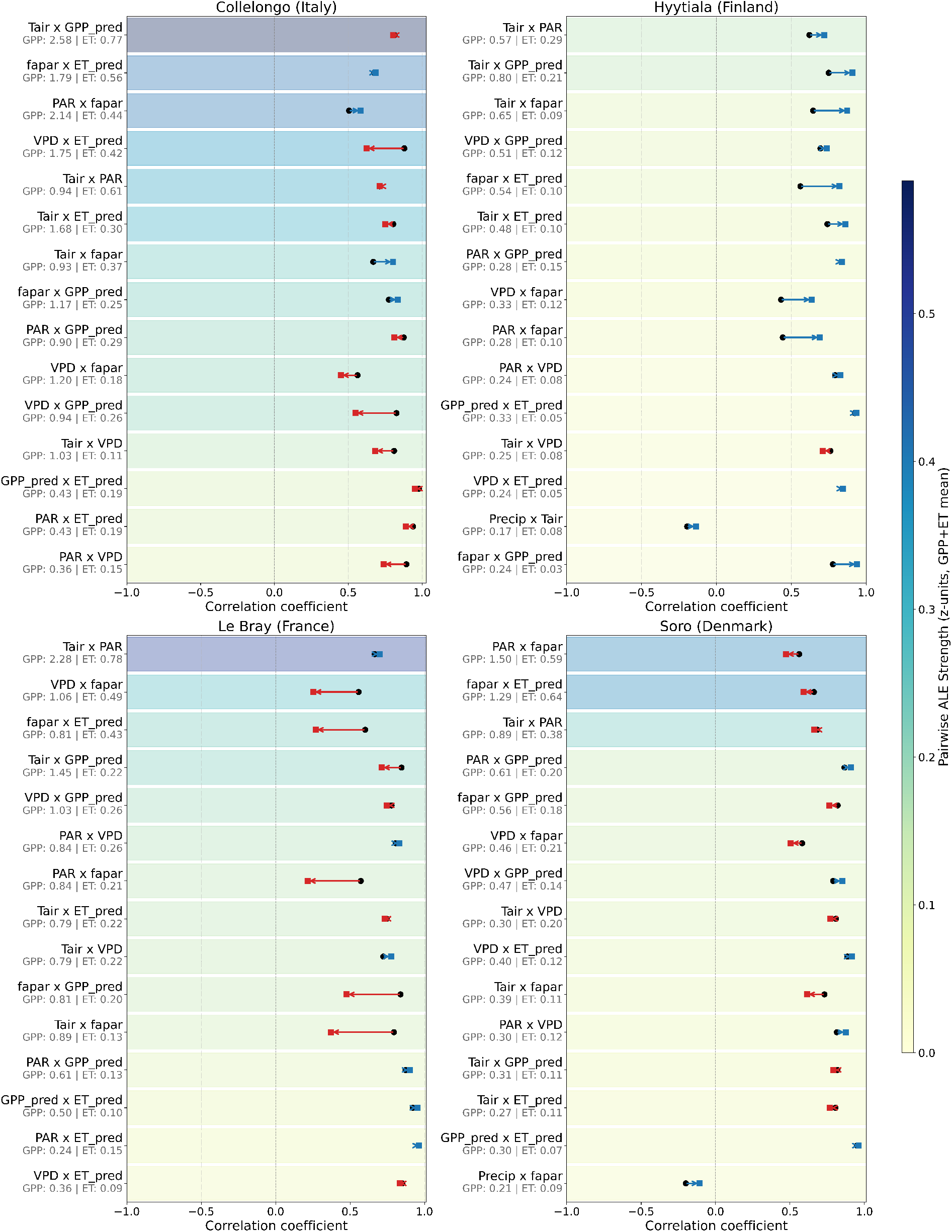
Domain-Shift ***Residual*** model: Predictor-predictor correlation shifts ranked by 2nd-Order ALE values showing the interaction effect on the targets (mean) per modelling strategy at Le Bray. Black circles represent the correlation coefficient within the training data, and the arrow indicates how this correlation changed at the test site. ALE range reflects the amplitude range of that feature pair. Correlation pairs filtered by strongest to weakest ALE amplitudes. Heatmap colours are drawn in z-scored units to keep GPP and ET comparable. The labels show the ALE-calculated influence in physical units per target.

## Footnotes

2 5403 days is the final span of the full available fAPAR record at each site; the period actually used for model training and evaluation after requiring all input variables to be jointly available is shorter and site-specific (2922–4748 days, Table 1), so the affected-day counts above are an upper bound over the full record rather than the modelled period specifically.

## References

Akiba, T., Sano, S., Yanase, T., Ohta, T., Koyama, M., 2019. Optuna: A next-generation hyperparameter optimization framework, in: Proceedings of the 25th ACM SIGKDD international conference on knowledge discovery & data mining, pp. 2623–2631.

Apley, D.W., Zhu, J., 2020. Visualizing the effects of predictor variables in black box supervised learning models. Journal of the Royal Statistical Society Series B: Statistical Methodology 82, 1059–1086. doi:10.1111/rssb.12377.

Besnard, S., Carvalhais, N., Arain, M.A., Black, A., De Bruin, S., Buchmann, N., Cescatti, A., Chen, J., Clevers, J.G., Desai, A.R., et al., 2018. Quantifying the effect of forest age in annual net forest carbon balance. Environmental Research Letters 13, 124018.

Camps-Valls, G., Gerhardus, A., Ninad, U., Varando, G., Martius, G., Balaguer-Ballester, E., Vinuesa, R., Diaz, E., Zanna, L., Runge, J., 2023. Discovering causal relations and equations from data. Physics Reports 1044, 1–68.

Chen, S., Zwart, J.A., Jia, X., 2022. Physics-guided graph meta learning for predicting water temperature and stream-flow in stream networks, in: Proceedings of the 28th ACM SIGKDD Conference on Knowledge Discovery and Data Mining, pp. 2752–2761.

Dormann, C.F., Elith, J., Bacher, S., Buchmann, C., Carl, G., Carré, G., Marquéz, J.R.G., Gruber, B., Lafourcade, B., Leitão, P.J., et al., 2013. Collinearity: a review of methods to deal with it and a simulation study evaluating their performance. Ecography 36, 27–46.

Dou, X., Yang, Y., Luo, J., 2018. Estimating forest carbon fluxes using machine learning techniques based on eddy covariance measurements. Sustainability 10, 203.

Fang, J., Gentine, P., 2024. Exploring optimal complexity for water stress representation in terrestrial carbon models: A hybrid-machine learning model approach. Journal of Advances in Modeling Earth Systems 16, e2024MS004308.

Grossiord, C., Buckley, T.N., Cernusak, L.A., Novick, K.A., Poulter, B., Siegwolf, R.T., Sperry, J.S., McDowell, N.G., 2020. Plant responses to rising vapor pressure deficit. New phytologist 226, 1550–1566.

Gupta, H.V., Kling, H., Yilmaz, K.K., Martinez, G.F., 2009. Decomposition of the mean squared error and nse performance criteria: Implications for improving hydrological modelling. Journal of hydrology 377, 80–91.

Hackenberg, M., Connor, S.G., Kabus, F., Brawner, J., Markham, E., Hardalupas, M., Chowdhury, A., Backofen, R., Köttgen, A., Rohde, A., et al., 2025. Small Data Explainer–The impact of small data methods in everyday life. arXiv preprint arXiv:2507.11773.

Hansen, N., 2016. The CMA evolution strategy: A tutorial. arXiv preprint arXiv:1604.00772.

Hestness, J., Narang, S., Ardalani, N., Diamos, G., Jun, H., Kianinejad, H., Patwary, M.M.A., Yang, Y., Zhou, Y., 2017. Deep learning scaling is predictable, empirically. arXiv doi:10.48550/arXiv.1712.00409.

Hettige, K.H., Ji, J., Xiang, S., Long, C., Cong, G., Wang, J., 2024. Airphynet: Harnessing physics-guided neural networks for air quality prediction. arXiv preprint arXiv:2402.03784.

Hsieh, W.W., 2009. Machine learning methods in the environmental sciences: Neural networks and kernels. Cambridge university press.

Jia, X., Willard, J., Karpatne, A., Read, J.S., Zwart, J.A., Steinbach, M., Kumar, V., 2021. Physics-guided machine learning for scientific discovery: An application in simulating lake temperature profiles. ACM/IMS Transactions on Data Science 2, 1–26.

Jomar, D., 2020. PyALE: A Python implementation of accumulated local effect plots. URL: https://github.com/DanaJomar/PyALE.

Jung, M., Koirala, S., Weber, U., Ichii, K., Gans, F., Camps-Valls, G., Papale, D., Schwalm, C., Tramontana, G., Reichstein, M., 2019. The FLUXCOM ensemble of global land-atmosphere energy fluxes. Scientific data 6, 74.

Karimi, D., Dou, H., Warfield, S.K., Gholipour, A., 2020. Deep learning with noisy labels: Exploring techniques and remedies in medical image analysis. Medical image analysis 65, 101759.

Karpatne, A., Jia, X., Kumar, V., 2024. Knowledge-guided machine learning: Current trends and future prospects. arXiv preprint arXiv:2403.15989.

Koppa, A., Rains, D., Hulsman, P., Poyatos, R., Miralles, D.G., 2022. A deep learning-based hybrid model of global terrestrial evaporation. Nature communications 13, 1912.

Li, L., Jamieson, K., DeSalvo, G., Rostamizadeh, A., Talwalkar, A., 2018. Hyperband: A novel bandit-based approach to hyperparameter optimization. Journal of Machine Learning Research 18, 1–52.

Liu, L., Xu, S., Tang, J., Guan, K., Griffis, T.J., Erickson, M.D., Frie, A.L., Jia, X., Kim, T., Miller, L.T., et al., 2022. KGML-ag: a modeling framework of knowledge-guided machine learning to simulate agroecosystems: a case study of estimating N 2 O emission using data from mesocosm experiments. Geoscientific model development 15, 2839–2858.

Mahnken, M., Cailleret, M., Collalti, A., Trotta, C., Biondo, C., d’Andrea, E., Dalmonech, D., Marano, G., Mäkelä, A., Minunno, F., et al., 2022. Accuracy, realism and general applicability of European forest models. Global Change Biology 28, 6921–6943.

Mäkelä, A., Pulkkinen, M., Kolari, P., Lagergren, F., Berbigier, P., Lindroth, A., Loustau, D., Nikinmaa, E., Vesala, T., Hari, P., 2008. Developing an empirical model of stand GPP with the LUE approach: analysis of eddy covariance data at five contrasting conifer sites in Europe. Global change biology 14, 92–108.

Migliavacca, M., Musavi, T., Mahecha, M.D., Nelson, J.A., Knauer, J., Baldocchi, D.D., Perez-Priego, O., Christiansen, R., Peters, J., Anderson, K., et al., 2021. The three major axes of terrestrial ecosystem function. Nature 598, 468–472.

Minunno, F., Peltoniemi, M., Launiainen, S., Aurela, M., Lindroth, A., Lohila, A., Mammarella, I., Minkkinen, K., Mäkelä, A., 2016. Calibration and validation of a semi-empirical flux ecosystem model for coniferous forests in the Boreal region. Ecological Modelling 341, 37–52. 10.1016/j.ecolmodel.2016.09.020.

Murphy, A.H., 1988. Skill scores based on the mean square error and their relationships to the correlation coefficient. Monthly weather review 116, 2417–2424.

Nash, J.E., Sutcliffe, J.V., 1970. River flow forecasting through conceptual models part I—A discussion of principles. Journal of hydrology 10, 282–290.

Nomura, M., Shibata, M., 2024. cmaes: A Simple yet Practical Python Library for CMA-ES. arXiv preprint arXiv:2402.01373.

Oehlert, G.W., 1992. A note on the delta method. The American Statistician 46, 27–29. doi:10.1080/00031305.1992.10475842.

Ozaki, Y., Tanigaki, Y., Watanabe, S., Onishi, M., 2020. Multiobjective tree-structured parzen estimator for computationally expensive optimization problems, in: Proceedings of the 2020 genetic and evolutionary computation conference, pp. 533–541.

Pearson, R.K., Neuvo, Y., Astola, J., Gabbouj, M., 2016. Generalized hampel filters. EURASIP Journal on Advances in Signal Processing 2016, 87.

Peltoniemi, M., Pulkkinen, M., Aurela, M., Pumpanen, J., Kolari, P., Makela, A., 2015. A semi-empirical model of boreal-forest gross primary production, evapotranspiration, and soil water - calibration and sensitivity analysis. Boreal Environment Research 20, 151–171.

Pichler, M., Hartig, F., 2023. Machine learning and deep learning — A review for ecologists. Methods in Ecology and Evolution 14, 994–1016.

Reichstein, M., Bahn, M., Mahecha, M.D., Kattge, J., Baldocchi, D.D., 2014. Linking plant and ecosystem functional biogeography. Proceedings of the national academy of sciences 111, 13697–13702.

Reichstein, M., Camps-Valls, G., Stevens, B., Jung, M., Denzler, J., Carvalhais, N., Prabhat, F., 2019. Deep learning and process understanding for data-driven Earth system science. Nature 566, 195–204.

Reyer, C.P., Silveyra Gonzalez, R., Dolos, K., Hartig, F., Hauf, Y., Noack, M., Lasch-Born, P., Rötzer, T., Pretzsch, H., Meesenburg, H., et al., 2020. The PROFOUND Database for evaluating vegetation models and simulating climate impacts on European forests. Earth System Science Data 12, 1295–1320.

Roberts, D.R., Bahn, V., Ciuti, S., Boyce, M.S., Elith, J., Guillera-Arroita, G., Hauenstein, S., Lahoz-Monfort, J.J., Schröder, B., Thuiller, W., et al., 2017. Cross-validation strategies for data with temporal, spatial, hierarchical, or phylogenetic structure. Ecography 40, 913–929.

Shazeer, N.M., Mirhoseini, A., Maziarz, K., Davis, A., Le, Q.V., Hinton, G.E., Dean, J., 2017. Outrageously Large Neural Networks: The Sparsely-Gated Mixture-of-Experts Layer. ArXiv abs/1701.06538.

Stocker, B.D., Zscheischler, J., Keenan, T.F., Prentice, I.C., Peñuelas, J., Seneviratne, S.I., 2018. Quantifying soil moisture impacts on light use efficiency across biomes. New Phytologist 218, 1430–1449.

Stull, R.B., 1988. An Introduction to Boundary Layer Meteorology. Atmospheric and Oceanographic Sciences Library, Kluwer Academic, Dordrecht, Netherlands. p.641.

Sun, N.Z., Sun, A., 2015. Model calibration and parameter estimation: for environmental and water resource systems. Springer. 10.1007/978-1-4939-2323-6.

Taiz, L., 2015. Plant physiology and development. Sinauer Associates. Incorporated. p.172–174.

Tramontana, G., Jung, M., Schwalm, C.R., Ichii, K., Camps-Valls, G., Ráduly, B., Reichstein, M., Arain, M.A., Cescatti, A., Kiely, G., et al., 2016. Predicting carbon dioxide and energy fluxes across global FLUXNET sites with regression algorithms. Biogeosciences 13, 4291–4313.

Uyekawa, J., Leland, J., Bergl, D., Liu, Y., Richardson, A.D., Lucas, B., 2025. Machine Learning-Based Prediction of Ecosystem-Scale CO2 Flux Measurements. Land 14, 124.

Van Oijen, M., Rougier, J., Smith, R., 2005. Bayesian calibration of process-based forest models: bridging the gap between models and data. Tree physiology 25, 915–927.

Verma, Y., Heinonen, M., Garg, V., 2024. ClimODE: Climate and weather forecasting with physics-informed neural ODEs. arXiv preprint arXiv:2404.10024.

Viovy, N., Arino, O., Belward, A., 1992. The best index slope extraction (bise): A method for reducing noise in ndvi time-series. International Journal of remote sensing 13, 1585–1590.

Von Rueden, L., Mayer, S., Beckh, K., Georgiev, B., Giesselbach, S., Heese, R., Kirsch, B., Pfrommer, J., Pick, A., Ramamurthy, R., et al., 2021. Informed machine learning - a taxonomy and survey of integrating prior knowledge into learning systems. IEEE Transactions on Knowledge and Data Engineering 35, 614–633.

Wesselkamp, M., Moser, N., Kalweit, M., Boedecker, J., Dormann, C.F., 2024. Process-Informed Neural Networks: A Hybrid Modelling Approach to Improve Predictive Performance and Inference of Neural Networks in Ecology and Beyond. Ecology Letters 27, e70012.

Willard, J., Jia, X., Xu, S., Steinbach, M., Kumar, V., 2022. Integrating scientific knowledge with machine learning for engineering and environmental systems. ACM Computing Surveys 55, 1–37.

Williams, M., Rastetter, E.B., Fernandes, D.N., Goulden, M.L., Shaver, G.R., Johnson, L.C., 1997. Predicting gross primary productivity in terrestrial ecosystems. Ecological Applications 7, 882–894.

Yang, Q., Liu, L., Zhou, J., Ghosh, R., Peng, B., Guan, K., Tang, J., Zhou, W., Kumar, V., Jin, Z., 2023. A flexible and efficient knowledge-guided machine learning data assimilation (KGML-DA) framework for agroecosystem prediction in the US Midwest. Remote sensing of environment 299, 113880.

Yu, R., Qiu, C., Ladwig, R., Hanson, P.C., Xie, Y., Li, Y., Jia, X., 2024. Adaptive process-guided learning: An application in predicting lake do concentrations, in: 2024 IEEE International Conference on Data Mining (ICDM), IEEE. pp. 580–589.

Zhou, K., Liu, Z., Qiao, Y., Xiang, T., Loy, C.C., 2022. Domain generalization: A survey. IEEE transactions on pattern analysis and machine intelligence 45, 4396–4415.

